# Discovery of a DNA-based Optical Nanotube Sensor for Glucose Using Clustering and Deep Learning Algorithms

**DOI:** 10.1101/2025.05.06.652529

**Authors:** Yahya Rabbani, Joey Bregy, Benjamin Rousseau, Sayyed Hashem Sajjadi, Lou De Benedittis, Sara Behjati, Mohammed Mouhib, Subhasis Dehury, Ardemis A. Boghossian

**Affiliations:** Ecole Polytechnique Fédérale de Lausanne (EPFL), 1015-Lausanne, Switzerland

**Keywords:** near-infrared fluorescence (NIR-II), single-walled carbon nanotubes (SWNTs or SWCNTs), continuous glucose monitoring (CGM), directed evolution, clustering, deep neural network learning, pattern recognition

## Abstract

Glucose sensing is vital for managing diabetes. However, current sensors are invasive and secrete enzymatic byproducts that are inflammatory and toxic. Though DNA-wrapped single-walled carbon nanotubes (DNA-SWCNTs) emit fluorescence that is ideal for enzyme-free optical sensing, existing approaches have yet to identify a DNA sequence that can elicit a fluorescence response to glucose. We develop an approach based on clustering to design a diverse library of 90 DNA sequences to screen for a glucose response. The most responsive sequence was further improved based on favorable mutations predicted by deep learning and pattern recognition. This combination of experimental screening and supervised and unsupervised machine learning represents a generalizable approach to developing DNA-SWCNT sensors for even the most elusive analytes.

**Graphical Abstract:** 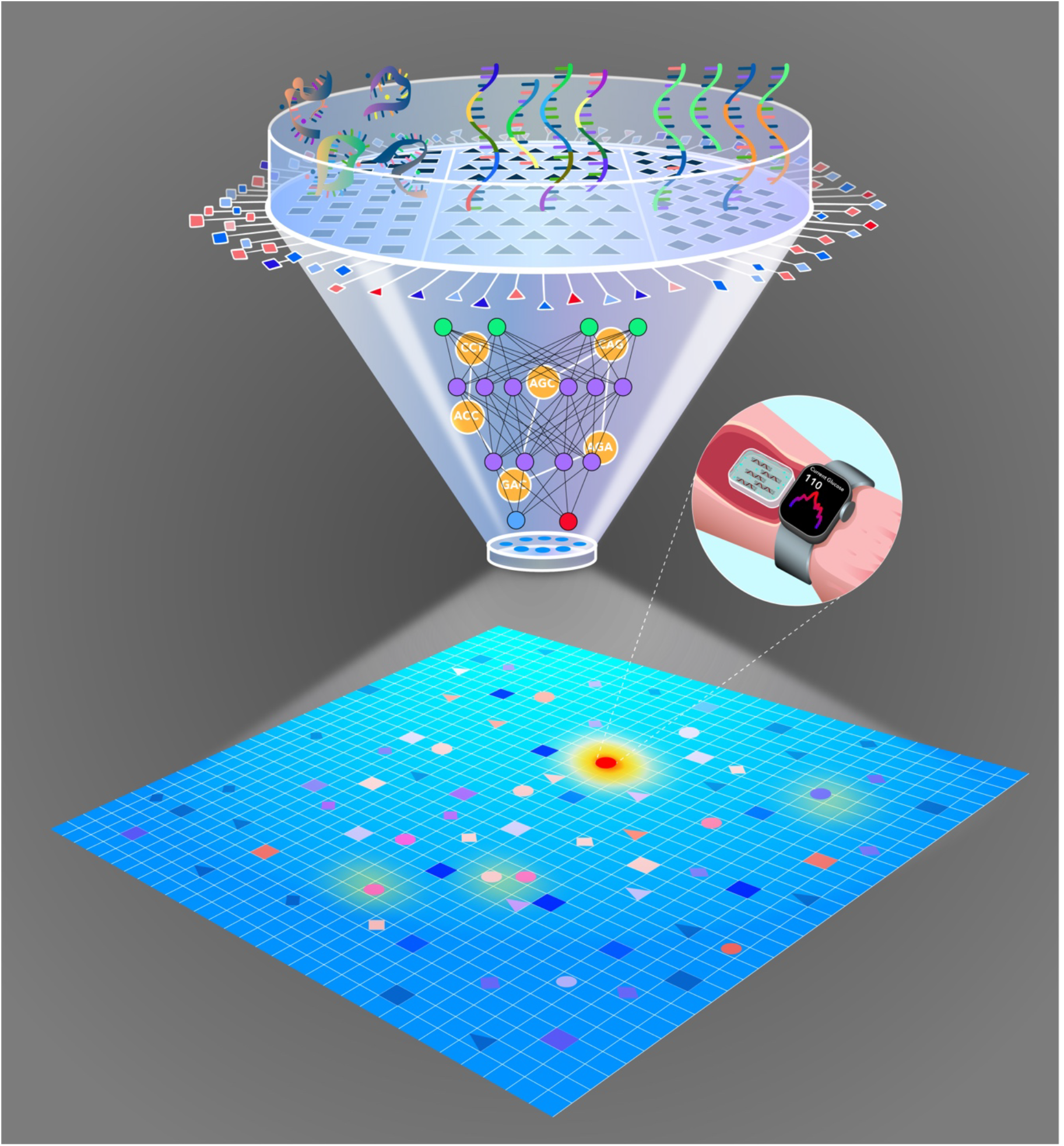

---

Diabetes mellitus is a chronic metabolic disease that will affect 642 million individuals by 2040 (*1*). It results in elevated blood sugar levels that can lead to severe health complications such as blindness and amputation (*1-4*). Without a cure, patients must manage their sugar levels through glucose monitoring. However, conventional methods based on electrochemical glucometers are inconvenient, painful, and rely on only on intermittent measurements taken by the patient. While continuous glucose monitoring (CGM) offers an automated approach for continuous measurements, the sensors are costly and require frequent calibration (*5-8*). Furthermore, they often rely on enzymes that generate hydrogen peroxide, an inflammatory and toxic byproduct. Consequently, the CGM sensors must be affixed externally with a penetrating needle and adhesive patch.

Optical sensors, on the other hand, enable contactless measurements that keep the skin intact after their insertion. These sensors especially benefit from probes that operate with near-infrared (NIR-II) emissions that can penetrate deep into biological tissue. Single-walled carbon nanotubes (SWCNTs) are among the few optical probes that fluoresce at these wavelengths (*9-14*). Their indefinite photostability and fluorescence response to even the slightest perturbations in their environment make SWCNTs highly suitable as CGM sensors. However, SWCNTs require surface functionalization for solubilization and control over the specificity of their fluorescence response (*13,14, 15*). Reversible SWCNT glucose sensors to date have thus largely relied on the surface immobilization of glucose-specific enzymes like glucose oxidase (*16-21*). However, in addition to the cytotoxic generation of hydrogen peroxide, the enzyme results in the poor colloidal stability of the solubilized SWCNTs.

In contrast, single-stranded DNA (ssDNA) can readily solubilize SWCNTs through the pi-pi stacking of their bases on the SWCNT surface (*21,22*). Importantly, the specificity of these ssDNA-wrapped SWCNTs (ssDNA-SWCNTs) can be tuned by varying the ssDNA sequence. Comprising four natural nucleotides, ssDNA offers a diverse and extensive sequence space of 4^n possible combinations, where n represents the nucleotide length of the ssDNA, for tuning the fluorescence properties of the SWCNTs. However, the effect of the ssDNA sequence on the fluorescence is unpredictable. This lack of information limits sensor design to the empirical and random screening of several sequences against a library of analytes (*23*).

In our recent work (*24,25*), we developed an approach inspired by directed evolution to tune ssDNA-SWCNT sensor properties in a guided manner. In this approach, an initial library of ssDNA-SWCNTs is screened against a specific analyte. Sequences that show the desired fluorescence behavior are selected and randomly mutated. The library of new ssDNA-SWCNT mutants is screened to identify sequences with improved characteristics, and the rounds of mutation and screening are repeated to iteratively improve sensor performance. This approach has been applied to enhance different sensor properties including selectivity, responsivity, brightness, and chiral-specific wavelength response (*24,25*).

Despite the ability to control multiple sensor properties, the approach relies on random mutations for improving sensor properties. Though recent studies have used machine learning to identify patterns in the ssDNA features that control the fluorescence of ssDNA-SWCNTs (*26*), these studies have been largely applied to unrelated and diverse sequences, as opposed to related mutated sequences. Considering the vastness of the possible space and the limited experimental data from 100-200 random sequences, these models are relatively limited in the accuracy of their predictions for related sequences with single- or few-nucleotide mutations. Furthermore, while directed evolution is useful for improving an existing property, it cannot create properties that do not already exist in the original, unmodified sequence. Therefore, the approach fails if the initial screening of ssDNA-SWCNTs does not identify any sequences that yield a response to the analyte. The directed evolution approach is therefore limited by the diversity and size of the initial screening library, especially when the desired characteristic can only be observed among a select number of related sequences (e.g. “a needle in a haystack”).

To address these limitations, we develop a combinatorial approach based on unsupervised and supervised machine learning to create a ssDNA-SWCNT sensor for glucose (Figure 1S; Figure 1A, left). Glucose represents a relatively inactive analyte for which a ssDNA sequence has never been identified through random screening. The DNA sequences are screened based on the change in the peak intensity and/or shifting of the (9,4) and (7,6) SWCNT chirality fluorescence wavelengths in response to glucose (Figure 1B, right). To engineer this sequence, we apply clustering, an unsupervised machine learning strategy, to categorize the relatedness of ssDNA sequences into different clusters (Step 1). Two million random, computationally generated sequences are clustered according to different properties, such as individual nucleotide composition (MAFFT), presence and frequency of nucleotide codons of K-length (K-mer), and folding properties (Folding). Representative sequences from different clusters are then selected to diversify the initial ssDNA-SWCNT library. After screening the initial library (Step 2), the sequence(s) that show(s) a response is selected and mutated through clustering to ensure the diversity of the mutations (Step 3). The data are used to train supervised deep learning and pattern recognition models (Step 4). The trained models are used to predict the effect of randomly generated mutations, and the performances of the most promising mutants are experimentally validated (Step 5). The new data are used to further train the models (Step 4), and the validation (Step 5) and training (Step 4) steps are repeated until the sequence converges to a local or, ideally, global maximum response (Step 6). Beyond the discovery of a sensor for a conventionally inactive and important analyte, this approach opens the doors to the unsupervised automated design of ssDNA-SWCNTs that is generalizable to most analytes and sensor properties.

**Fig. 1.**
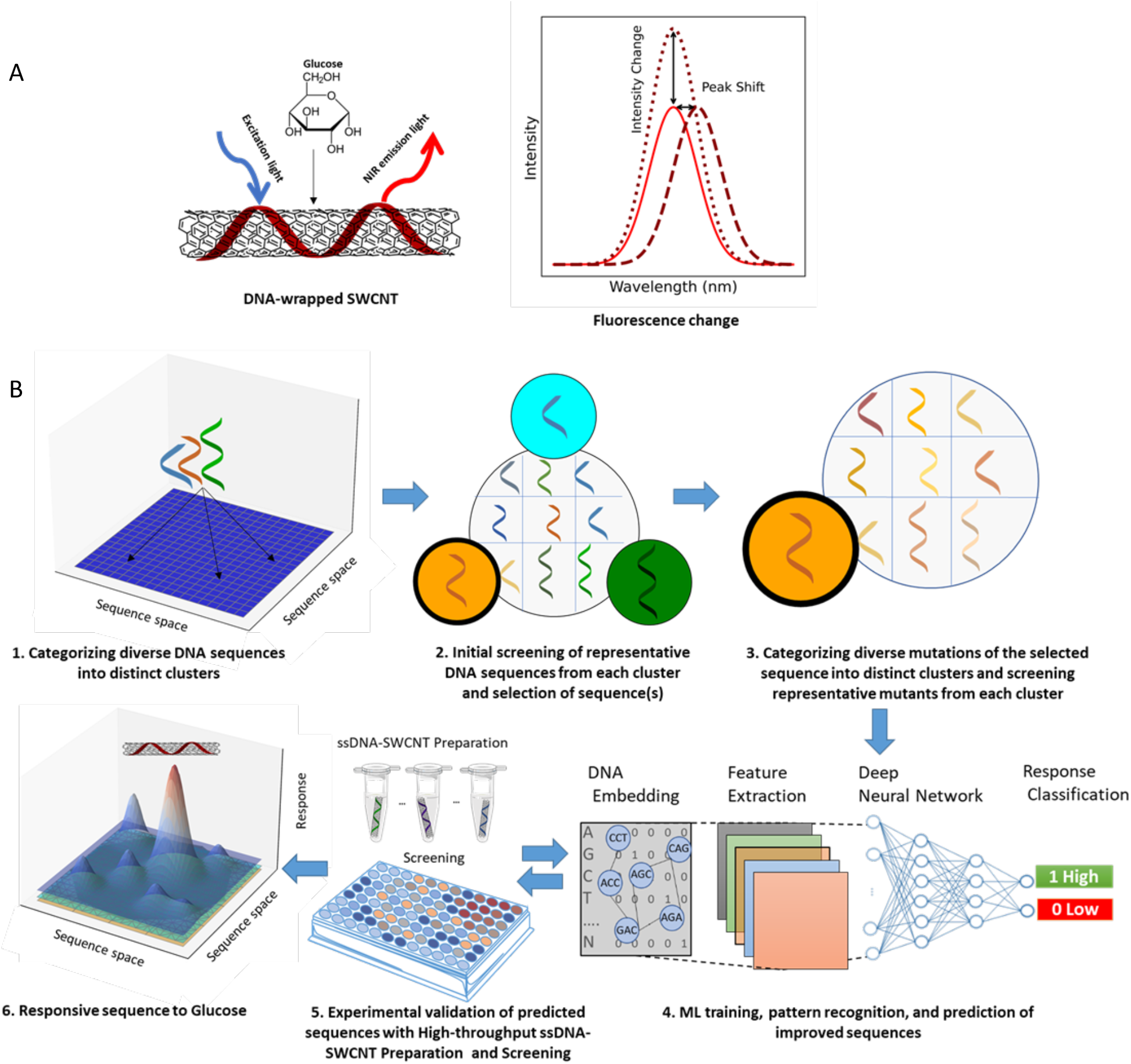
Overview of clustering- and deep neural network-assisted screening of DNA sequences for optical glucose detection with SWCNTs. **A)** (left) The ssDNA wrapping (red) is engineered to elicit a SWCNT fluorescence response to glucose. (right) The emitted fluorescence (solid line) may undergo a peak shift (dashed line) and/or intensity change (dotted line) in response to glucose. The schematic shows the wrapping of the ssDNA on a single SWCNT chirality and the corresponding fluorescence peak for that chirality. Actual measurements were done with a commercially available mixture containing several SWCNT chiralities. The same ssDNA sequence was used to suspend multiple chiralities in the same mixture, yielding multiple fluorescence peaks for each ssDNA sample that are chirality-specific **B)** Strategy for using unsupervised and supervised machine learning methods to navigate the different possible DNA sequences through clustering and learning. (1) Two million generated DNA sequences are categorized into different clusters based on relative similarities. (2) Representative sequences from each of the different clusters are experimentally screened for a glucose response. (3) The most responsive sequence(s) is selected, and mutations of the selected sequence(s) are then categorized into different clusters. Each mutant harbors 3 nucleotide and/or length mutations relative to the selected sequence. Representative sequences from each of the different mutant clusters are experimentally screened for a glucose response. (4) Pattern recognition analyzes the positions and length of motifs that interact with glucose. A deep neural network predicts the performances of new DNA mutants. The different deep neural networks include Graph Convolutional Networks (GCNs), Graph Neural Networks (GNNs), and Convolutional Neural Networks(CNNs). (5) The sequence(s) with the highest predicted performance(s) is experimentally validated in a high-throughput setup, and the results are used to further train the model in (4). Steps (5) and (4) are repeated until (6) the sensor benchmark is achieved or the performance cannot be further optimized.

## DNA Design by Clustering and Mutations

Initial attempts based on the random screening of 90 sequences (Figure S2) and of previously engineered ssDNA sequences, including those that have been previously reported to react with other analytes (Figure S3) (*27-34*), were unable to identify sequences that elicit any response to glucose. The search was therefore expanded to include two million computationally generated DNA sequences with varying lengths of 6 to 30 bases and different base compositions (Figure S4). Of the two million sequences, 1 million were generated based on randomized individual nucleotides and 1 million were generated based on randomized nucleotide motifs or “codons” of K-length.

Three different clustering methods were used to classify the two million sequences DNA based on differences in individual nucleotide composition (MAFFT), presence and frequency of nucleotide motifs of K-length (K-mer), and DNA folding (Folding) (Figure 2A). Each clustering method thus represents a distinct depiction and binning of relatedness of the DNA sequence space (Figure S7 and S8). For the MAFFT method, the relatedness of the sequences was based on the alignment of their bases, and a similarity index was calculated for each sequence. The sequences were then clustered by similarity using the k-means method (Figure S5). In the K-mers method, the frequency of different 3-mer patterns, such as GTC, ATC, etc., was analyzed, and sequences were clustered based on the types and frequencies of the 3-mers (Figure S6). For the Folding method, information on DNA folding information was categorized according to ViennaRNA predictions (*35,36*) (Table S1). The different bins of related sequences, or clusters, in each method are thus intended to capture the diversity of sequences from the entire ssDNA sequence space; the selection of representative sequences from each cluster allows efficient screening of the entire space without screening every possible sequence. The elbow method was used to determine the optimal number of clusters for each method (Figure 2B), and one representative sequence from each cluster was selected for further analysis.

**Fig. 2.**
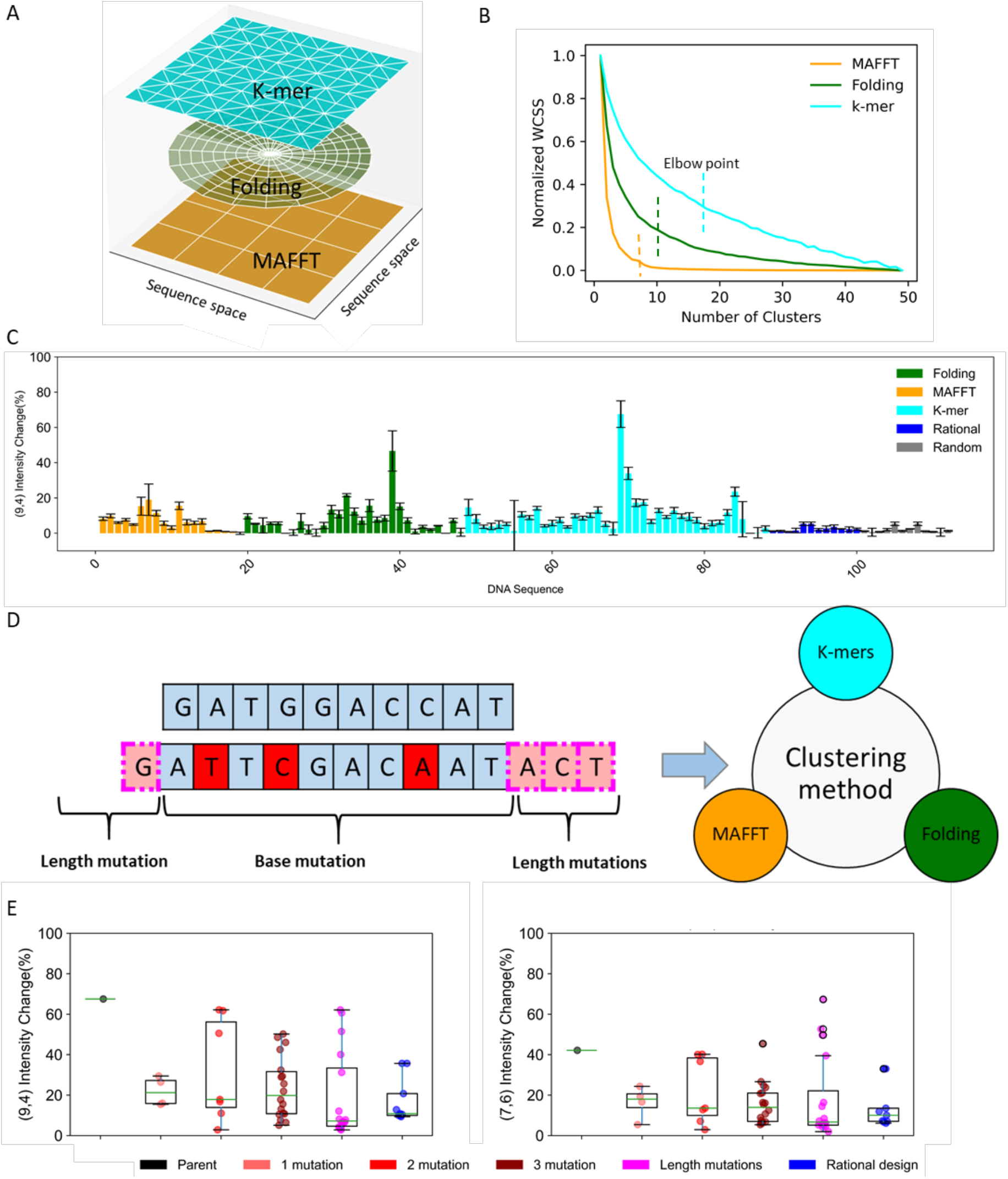
Different clustering methods for diversifying the initial library and DNA mutations. A) All possible DNA sequences depicted as an x-y plane. The white markings separate the different clusters of DNA, whereby sequences in the same cluster share similar metrics. The different metrics used to assess similarity and cluster the sequences include MAFFT (orange), Folding (green), and K-mers (cyan). The corresponding depiction of the sequence space for each metric is represented as a separate plane. B) Diversity of sequences within a cluster, represented as the within cluster sum of square (WCSS), as a function of the number of clusters for the MAFFT (orange), Folding (green), and K-mers (where K = 3) (cyan) clustering methods. The WCSS is normalized by the sum of square of all the tested DNA sequences (i.e. at maximum WCSS where the number of clusters is 0). The vertical dotted lines represent the elbow point for determining an optimum number of clusters for each method. C) Experimental (9,4) intensity changes of different ssDNA-SWCNT sequences in response to 7.5 mM glucose. Representative sequences are shown from each cluster from the MAFFT (orange), folding properties (green), and K-mers (where K = 3) (cyan) clustering methods as well as from rational design (blue) and from a random sequence library (grey). D) (left) Schematic representation of base and length mutations for a representative GATGGACCAT sequence (parent). Red boxes represent base mutations performed on 1, 2, or 3 randomly selected nucleotides. Salmon boxes represent length mutations through the random addition of 1, 2 or 3 nucleotides added on one or independently on both sides of the sequence. (right) Mutations are grouped into different clusters using the MAFFT, Folding, and K-mers (where K = 3) methods. E) Experimental (9,4) (left) and (7,6) (right) intensity changes of mutants harboring 1 (salmon), 2 (red), and 3 (maroon) base mutations as well as assorted length mutations (pink) of the parent sequence. The DNA sequences are listed in Table S2 and Table S3.

The selected ssDNA sequences were wrapped around SWCNTs, as depicted in Figure 1A, using a previously reported exchange method with SC-SWCNTs (*15*). The fluorescence spectra of each ssDNA solution were recorded at two different excitation wavelengths to capture the response of six different SWCNT chiralities contained within each solution. Though the majority of chiralities showed no glucose response for any of the ssDNA sequences, the (9,4) and the (7,6) chiralities showed a peak shifting and intensity response on the addition of 7.5 mM glucose when wrapped with specific ssDNA sequences (Figure S9). The (9,4) chirality also showed a pronounced fluorescence increase compared to the negative control (Figure 2C). As shown in the figure, all clustered sequences showed greater diversity in their responses compared the random and rational approaches based on previous approaches (Table S2) (*27-34*). Notably, sequences clustered with the K-mer method showed the greatest diversity, with the Clustered_Seq_69 sequence (GATGGACCAT) achieving an approximately 70% increase in intensity and a ~2.5 nm blue shift in the fluorescence peak.

The Clusterd_Seq_69 sequence was thus selected as the parent sequence for further improving the response. The sequence was modified by introducing base mutations at 1, 2, or 3 randomly selected nucleotides and length mutations of 1, 2, or 3 nucleotide additions at either one or both ends independently (Figure 2D, left). The 8000 computationally generated mutants were clustered using the MAFFT, K-mers, and Folding methods (Figure 2D, right), and representative mutants from each cluster were selected for screening (Figure S10, Figure 2E). As shown in Figure 2E, increasing the number of base mutations from one to two increases the diversity of the responses, which achieves a larger spread of both larger and smaller responses. The length mutations show the greatest diversity of responses, though they include a larger range 0-6 possible nucleotide differences. Although none of the DNA mutants showed any increase in the (9,4) intensity response compared to the parent sequence, several of the DNA mutants showed an increase in the (7,6) intensity response particularly for the length mutations. Several sequences, however, showed a (9,4) peak position response. Notably the Mutated-Clustered_Seq_42 sequence (AGGGATGGACCATAAT) showed a blue shift of up to 4 nm at the (9,4) peak and an increase of up to 70% in the (7,6) fluorescence intensity change (Figure 2E, Figure S11, and Table S3).

## Deep Neural Network Training, Pattern Recognition, and Prediction

The experimental results from the initial library (Table S2) and the mutations (Table S3) (Steps 2 and 3 of Figure 1B) were used to train deep neural network model (Step 4 of Figure 1B). The fluorescence intensity changes of the (7,6) and (9,4) chiralities upon glucose addition were designated as high-intensity (Class 1) and low-intensity (Class 0) responses based on a threshold (Figure S12). The deep neural network methods CNN, GNN, and GCN were then used to independently classify the intensity response, as described in the Methods (Figure 3A, Table S4). Test sequences were then ultimately classified based on majority vote among the three different models. Using this approach, most of the data were correctly classified as true low- and high-intensity responses, achieving a true positive to false positive ratio of 0.9 and 0.87 (Figure 3B, 3C, and S13) and high accuracy scores of 0.81 and 0.90 for the (7,6) and (9,4) chiralities, respectively (Table S5).

**Fig. 3:**
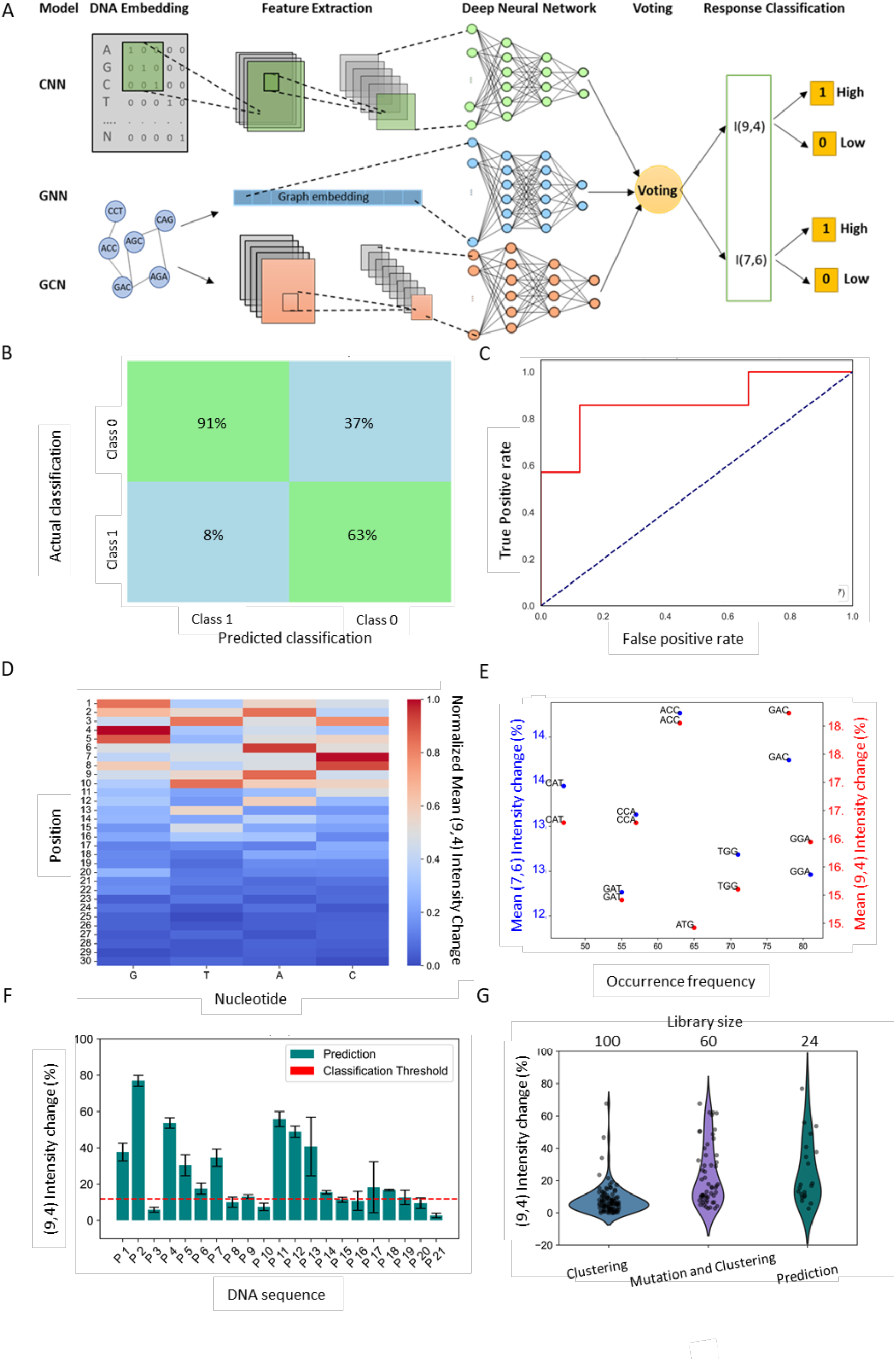
Deep neural network training, sequence pattern recognition, and prediction. **A**) Schematic of using CNN, GNN, and GCN neural network models for training and test data to predict the (9,4) and (7,6) chirality fluorescence intensity responses (I(9,4) and I(7,6), respectively) for different DNA sequences. The results of the models are weighted in the voting step to obtain individual predictions on the performances of different DNA sequences. The performances are classified based on high (Class 1) or low (Class 0) response. **B)** Confusion matrix comparing the predicted classification from the model and the actual performance based on experimental measurements. The percentages represent the fraction of predicted Class 0 (left quadrants) or Class 1 (right quadrants) (9,4) intensity responses that were experimentally validated as either Class 0 and Class 1. Green regions represent quadrants that designate consistency (true positive and true negative) between the predicted and experimental classifications and blue regions represent quadrants that designate inconsistency (false positive and false negative) between the predicted and the experimental classifications. **C)** Retention of Curve (ROC) (red) comparing the true and false positive classification rates. The blue dotted line represents equivalent true positive and false positive prediction rates for a random classifier. **D)** Investigation of sequence patterns to identify meaningful base positions affecting the (9,4) intensity change response for glucose detection. The bases yielding the most significant effects on fluorescence response are highlighted in red, while less significant effects are shown in blue, across different positions of DNA sequences ranging from 1 to 30 bases. **E)** K-mers (3 bases) of high-frequency and with high mean response effects on fluorescence intensity changes, with red dots representing the (9,4) chirality and the (7,6) chirality shown separately. **F)** Experimental validation of the response of DNA sequences predicted by deep neural network to have a high probability of strong response to glucose. **G)** Comparison of the experimental (9,4) chirality glucose sensing results across the main computational steps used in this study.

Pattern recognition was also applied to identify key sequence elements that enhance the sensor response to glucose (Figure 3D and 3E). These analysis identified sequence length (Figure S14), base composition (Figure S15 and S16), base position (Figure 3D and Figure S17 and S18), and K-mers (Figure 3E and Figure S19 and S20) as important features that govern the glucose response. According to the results, shorter sequences (less than 16 bases) that have specific bases at particular sequence positions have a dominating effect on increasing the glucose response. For example, G at positions 4 and 5, A at position 6, C at position 7, and T at position 10 all showed enhanced glucose sensitivity. Additionally, specific K-mers, namely the GAC, ACC, and GGA 3-mers, were frequent in highly responsive sequences (Table S6).

Finally, the majority voting approach from the trained deep neural network models was used to predict the behavior of new DNA sequences with improved performances (Figure 3F and 3G). Pattern recognition was first used to computationally generate 160000 new DNA sequences based on beneficial base positions and K-mers. The deep neural network model was used to predict the responses of these new DNA sequences, in addition to the responses of 8000 sequences from the previous clustering and mutation library. The most promising sequences predicted by the deep neural network (Figure S21) were experimentally tested (Figure 3F and Figure S22). The results revealed new sequences with even higher glucose responses that show up to an 80% intensity increase in the (9,4) chirality, up to a 60% intensity increase in the (7,6) chirality, and up to a 4-nm blue shift in the (9,4) chirality. Finally, we compared the experimental results across the main computational steps used in this study (clustering, mutations and clustering, and deep neural network prediction) (Figure 3G and Figure S23). Already the application of clustering to better navigate the DNA sequence space, allowed us to identify a ssDNA sequence responsive to glucose in ssDNA-SWCNT sensors for the first time. Furthermore, the data shows that with every step, we are able to identify sequences with higher analyte sensitivity, while significantly decreasing the library size that has to be experimentally screened. Thus, the final deep neural network model, decreases experimental efforts with a much higher success rate for glucose sensing, using predicted DNA sequences (Table S7).

## Sensor Application and ssDNA Binding Conformation

While our general workflow for development of sensitive ssDNA based optical biosensors proved to be successful, another significant outcome of our results thus far is the first identification of a glucose specific ssDNA-SWCNT sensor, with implications in non-invasive and continuous optical glucose monitoring. The performance of our most responsive glucose sensor was therefore assessed by first evaluating its analytical capabilities within a range of biologically relevant concentrations, followed by testing in media that mimic in vivo conditions, such as serum (Figure 4 A-D). The calibration curve of the sensor indicated a detection limit as low as 2.5 mM for glucose, displaying a sigmoidal relationship between both peak shift and intensity change as a function of increasing glucose concentration in PBS (Figure 4A). This behavior demonstrates the sensor’s utility within physiological concentration ranges. Additionally, the sensor exhibited a rapid response to glucose, with detection achieved in less than 5 minutes (Figure 4B), enabling real-time glucose monitoring. To assess the sensor’s suitability for optical device integration, its reversibility was tested across three cycles. Comparable peak shifts and comparable fluorescence intensity responses (Figure 4C) were observed during binding, release, and re-binding of glucose, confirming the sensor’s capacity for reversible glucose detection. Furthermore, to better approximate in vivo sensing conditions, the sensor’s ability to detect glucose in the presence of 10 % serum was evaluated. The results showed a time-dependent response, with significant detection occurring at short incubation times (Figure 4D). For in-vivo application, it is crucial that the sensor is biocompatible. Samples of the sensor where assayed for cytotoxicity using injection in Mice (performed at GLOW Biosciences). Optical monitoring revealed no significant production of ROS (Figure S25), and serum samples for biochemical measurements confirmed biocompatibility, as there was no increase in toxicity markers 24 h post-injection (Table S8).

**Fig. 4.**
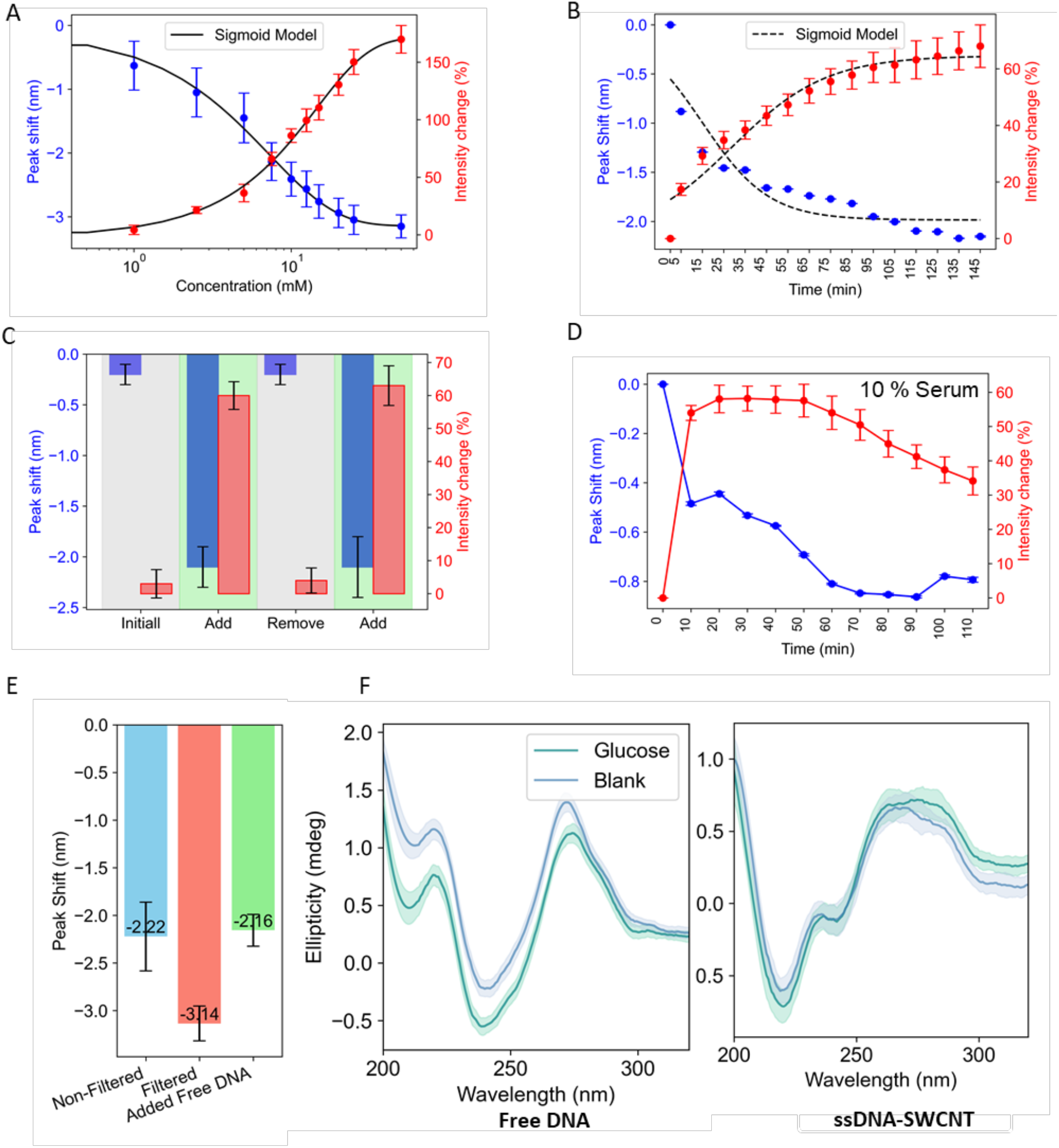
Sensor characteristics and DNA binding conformation. A) Concentration calibration curve showing the sensor’s response, including peak shift (blue dots) and intensity change (red dots), across a range of glucose concentrations. A sigmoid fit is applied to the log-transformed glucose concentration data. B) Sensor peak shift and intensity change response over time, measured from 5 minutes to 2 hours, with a sigmoid correlation demonstrating sensor functionality over time. C) Reversibility of the sensor over multiple cycles, showing the peak shift response during the addition (green) and removal of glucose. D) Sensor performance in 10% serum media after 10 minutes, measuring the peak shift (blue) and intensity change (red) responses. E) Effect of free ssDNA on the ssDNA-SWCNT sensor’s ability to detect glucose, measured by the peak shift response after removing free DNA from the sensor and adding free DNA. F) (left) CD measurements showing changes in DNA conformation in a free DNA sample with the addition of glucose. (right) CD measurements showing changes in DNA conformation on the surface of SWCNT in the ssDNA-SWCNT sample with the addition of glucose.

In the final stage of our study, we investigated the interaction mechanism between glucose and the ssDNA-SWCNT sensor to elucidate the observed peak shifts in SWCNT fluorescence (Figure 4 E and 4F). These fluorescence shifts can be attributed to either direct interaction of a given analyte with the SWCNT surface (*37,38*), or instead to conformational changes in the DNA wrapping around SWCNTs (*39,40*). Our initial findings indicated that the sensor’s response, evidenced by peak shift, improved when free DNA was removed from the ssDNA-SWCNT system (Figure 4E and S22). During the preparation of the sensor, free DNA remains present in the solution, representing an additional factor involved in the sensing process (*41,42*). This suggests that free DNA may directly bind to glucose, reducing the amount of glucose available to interact with the ssDNA on the SWCNT surface, thereby attenuating the sensor’s response. Moreover, this finding hints that the interaction between glucose and the sensor likely involves changes in the conformation and wrapping of the ssDNA around the SWCNTs. To confirm this hypothesis, we employed Circular dichroism (CD), a technique sensitive to structural changes in DNA both in solution (*43*) and when wrapped around SWCNTs (*39,44*) (Figure 4F). The binding of glucose to free DNA was confirmed by a peak shift towards higher wavelengths at 270 nm in the CD spectra (Figure 4F, left), indicating a conformational change in the DNA, a phenomenon not observed in glucose-unresponsive DNA sequences. The peak shift around 270 nm could also be attributed to the stacking of nucleotide bases (*45,47*) or the formation of a compact and ordered structure (*45*) (Figure S23). The CD spectrum of the glucose-responsive ssDNA-SWCNT sensor (Figure 4F, right) also showed a wavelength shift upon glucose addition, consistent with a conformational change in the ssDNA on the SWCNT surface. This conformational change, observed only in the responsive sensor (Figure S24), explains the fluorescence peak shift upon glucose binding.

## Acknowledgements

The authors thank Dr. Adrien Eric P Knoops for his valuable feedback and suggestions during the review of this work. They also thank Shokoufeh Hajatpour for designing the graphical abstract.

## Author contributions

Y.R., S.H.S., and A.A.B designed research. Computation was performed by Y.R. Experiments were performed by Y.R., J.B., B.R., L.B., S.B., and S.D. All authors contributed to the writing and reviewing of the paper.

## Data and materials availability

All the data related to clustering, pattern recognition, and deep learning methods, including descriptions of each part and Python scripts for different deep neural networks, are available on GitHub. Details on the convolutional neural networks (CNNs), graph neural networks (GNNs), and graph convolutional neural networks (GCNs) are also available on GitHub (https://github.com/YahyaRabbani20/AI-BioSense).

## Supplementary Materials

Materials and experimental methods

Computational methodology for clustering

Computational methodology for pattern recognition

Computational methodology for deep learning

Figures. S1 to S25

Tables S1 to S8

References (1-47)

