## Supplementary information for "Discovery of a DNA-based Optical Nanotube Sensor for Glucose Using Clustering and Deep Learning Algorithms"

**Supplementary Materials**

Materials and experimental methods

Computational methodology for clustering

Computational methodology for pattern recognition

Computational methodology for deep learning

Figures. S1 to S25

Tables S1 to S8

References (1–47)

### Materials and Methods of the Experimental Part

#### Materials

Purified SWCNTs, containing a mixture of chiralities, were purchased from NanoIntegris (HiPco). The ssDNA sequences were purchased from Microsynth. All other chemicals were purchased from Sigma-Aldrich.

#### ssDNA-SWCNT Preparation

To functionalize the ssDNA strands on the SWCNT surface, an existing exchange protocol was used (*13*). This method enables the replacement of the surfactant sodium cholate (SC), initially used to suspend the SWCNTs, with ssDNA strands. This approach allows for the high-throughput preparation of ssDNA-SWCNT complexes and facilitates the preparation of low-volume samples, thus reducing the cost of various screenings. The steps involved in preparing these ssDNA-SWCNT complexes are described in more detail below.

**SC-SWCNT and ssDNA Preparations**

A suspension of 30 mg of super-purified HiPco SWCNTs was prepared in 30 mL of a 2% SC solution in deionized water. This suspension was homogenized for 20 minutes at 5,000 rpm and sonicated for 1 hour using a probe-tip ultrasonicator at 10% amplitude in an ice bath. The suspension was then centrifuged at 30,000 rpm for 4 hours at 25°C, and the supernatant was collected. The concentration of SC-SWCNTs was adjusted to 110 mg/L by measuring the absorbance at 632 nm using a UV-Vis spectroscopic device (UV-3600 Plus, Shimadzu). The ssDNA oligomers were dissolved in DI water and their concentrations were adjusted to 50 µM by measuring absorbance at 260 nm (Nanodrop 2000 Thermo Scientific) using the OligoCalc website. The ssDNA samples were then stored at -20°C until further use.

**SWCNT Wrapping Exchange Method**

To perform the exchange, 60 µL of ssDNA solution was mixed with 60 µL of SC-SWCNT in 2 mL PCR-performant Eppendorf tubes. Then, 180 µL of methanol was added to increase the critical micelle concentration of the mixture. This caused SC molecules to leave the SWCNT surface and form micelles, allowing the ssDNA to wrap around the SWCNTs. The samples were then incubated overnight at room temperature (RT) to ensure complete wrapping and equilibration.

**ssDNA-SWCNT Purification**

Two purification steps were performed to remove SC and methanol and obtain pure ssDNA-SWCNT. In the first step, ssDNA-SWCNTs were pelleted by adding 46.2 µL of 1.5 M NaCl and 866 µL of cold 100% EtOH (stored at -20°C). The samples were stored at -20°C for 1 hour and then centrifuged at 4°C for 30 minutes at 15,000 rpm. The supernatant was discarded, and 1,000 µL of 70% EtOH was added for the second purification step, which also removed traces of salts. The ssDNA-SWCNT pellet was resuspended by vigorous mixing (pipetting up and down and vortexing). The samples were then centrifuged again for 1 hour at 21°C and 15,000 rpm. The supernatant was carefully removed, and after waiting 15 minutes to allow ethanol traces to evaporate without over-drying the pellet, 300 µL of DI water was added to resuspend the ssDNA-SWCNT. After 10 minutes, 10X PBS was added to adjust the sample concentration to 1X PBS. Samples were homogenized for over 1 hour before centrifugation (20 minutes, 21°C, 15,000 rpm). The supernatant was collected to remove potential aggregates. Finally, the concentration of ssDNA-SWCNTs in 1X PBS was adjusted to 5 mg/L by measuring absorbance at 632 nm for a 50 µL sample placed in a 384-well plate using a plate reader (Varioscan LUX, Thermo Scientific). The samples were stored overnight at RT in the dark before fluorescence measurements.

#### Fluorescence Spectroscopy Measurements

For the various screenings, the fluorescence of ssDNA-SWCNT samples was measured with and without the addition of glucose to determine fluorescence variations induced by glucose. To do this, 1 µL of either glucose (final concentration 7.5 mM in 1X PBS) or 1X PBS (as a blank) was added and mixed with 49 µL of ssDNA-SWCNT samples (5 mg/L), which were previously placed in a 384-well plate for screening. The glucose concentration of 7.5 mM was chosen as it represents the biological concentration in the blood of diabetic patients. Glucose or 1X PBS was added at 45-second intervals to mimic the operational timeline of the microscope, ensuring an equal incubation time of 2 hours for all samples. Each sample was replicated three times, and the fluorescence responses to glucose were averaged. The SWCNT fluorescence was monitored using a custom-built near-infrared microscope. This device was equipped with a supercontinuum laser, a tunable band-pass filter, and a short-pass filter. The laser was focused on the well plate using a 20X objective with a dichroic beam splitter, with an exposure time of 10 seconds at maximum relative power. Two excitations at 655 and 735 nm (with a bandwidth of 10 nm) were used to excite the samples, and the environment was maintained at RT. The fluorescence of the samples was detected by an InGaAs NIR detector and recorded via LightField software and a LabView program. A custom Python code using a Lorentzian fitting approach was used to construct the spectra relating intensity to the wavelength of the fluorescent emission. The positions and intensities of the peaks corresponding to the chiralities (7,5) and (7,6) obtained after excitation at 655 nm, and the chiralities (10,2), (9,4), (8,6), and (8,7) obtained after excitation at 735 nm, were extracted by deconvolving the emission peaks. This allowed for the calculation of fluorescence intensity change and peak shift due to glucose addition.

### Free DNA Filtration

The filtration steps were performed using Amicon Ultra-0.5 100 kDa devices. The membranes were first rinsed with DI water (500 µL). The ssDNA-SWCNT solutions were then added to the top compartment, totaling 500 µL, and filtered using centrifugal force (5000 x g for 5.30 minutes). The rotation speed and filtration time were adjusted so that 100 µL of solution remained in the top compartment of the filter. The ssDNA-SWCNTs were then washed four times by filling the upper filter compartment with DI water and ensuring complete resuspension of the ssDNA-SWCNTs by pipetting up and down. Finally, after the last round of filtration, DI water was added to obtain 200 µL of resuspended ssDNA-SWCNTs. The concentration of PBS was adjusted to 1X using 10X PBS, and the concentration of ssDNA-SWCNTs was adjusted for different experimental needs by measuring absorbance at 632 nm using a plate reader (Varioscan LUX, Thermo Scientific).

### Influence of Free DNA on the Response to Glucose

For this experiment, 49 µL of non-filtered, filtered (according to **Free DNA Filtration** section), or filtered with post-addition of DNA samples (5 µg/L in 1X PBS) were placed in different wells of a 384-well plate. Subsequently, 1 µL of either glucose (final concentration 7.5 mM) or 1X PBS (as a blank) was added and incubated for 2 hours. Each sample was replicated three times, and the fluorescence responses to glucose were averaged. Fluorescence measurements were acquired and analyzed similarly to the **Fluorescence Spectroscopy Measurements** section.

### Circular Dichroism (CD) measurement

The experiment was designed to measure the degree of ellipticity of ssDNA-SWCNTs (13 mg/L in 1X PBS) or free DNA (20 µM) with or without the addition of glucose (40 mM, 1-hour incubation) over the wavelength range of 200-340 nm. For each sample, three CD spectra were acquired (Chirascan, V100) and the ellipticity measurements were averaged. The CD spectra were then smoothed by applying a Savitzky-Golay filter with an order 4 polynomial fitting. The ssDNA-SWCNTs were previously filtered (according to **Free DNA Filtration** section) to prevent overlapping signals from free DNA in the solution.

### Influence of Glucose Concentration

49 µL of ssDNA-SWCNTs (5 mg/L in 1X PBS) was placed in different wells of a 384-well plate. Subsequently, 1 µL of either glucose or 1X PBS (as a blank) was added and incubated for 2 hours. The final glucose concentrations studied were: 5 mM, 7.5 mM, 10 mM, 12.5 mM, 15 mM, 20 mM, and 25 mM. Each condition was replicated three times, and the fluorescence measurements were averaged. Fluorescence measurements were acquired and analyzed similarly to those described in the **Fluorescence Spectroscopy Measurements** section.

### Influence of Incubation Time

49 µL of ssDNA-SWCNTs (5 mg/L in 1X PBS) was placed in different wells of a 384-well plate. Subsequently, 1 µL of either glucose (final concentration 7.5 mM) or 1X PBS (as a blank) was added and incubated for 5 minutes. Each sample was replicated three times, and the fluorescence measurements were averaged. Fluorescence measurements were acquired every 10 minutes for 2.5 hours, similarly to the **Fluorescence Spectroscopy Measurements** section.

**Response in Serum**

48 µL of ssDNA-SWCNTs (5 mg/L in 1X PBS) was placed in different wells of a 384-well plate. Subsequently, 2 µL of either glucose, dissolved in 1X PBS and mixed with various proportions of serum (final concentration 20 mM), or 1X PBS with the same proportions of serum (reference situation) was added and vigorously mixed. The different proportions of serum tested were 0%, 10%, 50%, and 75%. Each condition was replicated three times, and the fluorescence measurements were averaged. Fluorescence measurements were acquired after 2 and 4 hours of incubation, similarly to the **Fluorescence Spectroscopy Measurements** section.

**Reversibility test of ssDNA-SWCNT sensor**

In order to test the reversibility of the sensor, 800 µL of ssDNA-SWCNT were prepared and placed in different wells of a 384-well. Fluorescence measurements were acquired, similarly to the **Fluorescence Spectroscopy Measurements section** as initial timepoints. Subsequently, 1 µL of either glucose (to a final concentration of 20 mM) or 1X PBS was added and mixed vigorously and incubated for 1 hour. The response to glucose was then tested with fluorescent measurements. The samples were then filtered 5 times (similarly to the Free DNA filtration) in order to remove glucose using Amicon Ultra-0.5 3 kDa devices. Fluorescence measurements were acquired, similarly to the Fluorescence Spectroscopy Measurements, to evaluate if the initial intensity of the sensor prior to glucose addition was recovered. Either 1 µL glucose or 1X PBS was then added a second time and incubated 1 hour so that the recovery of the glucose response was evaluated. This procedure was then repeated 2 times to test the reversibility of the ssDNA-SWCNT sensor.

**Computational Methodology of the Clustering Section**

**Generating a Large-Sized DNA Sequence Library**

The methodology for generating a large-sized DNA sequence library involves two methods: DNA motif creation and DNA random creation. In the DNA motif creation step, possible DNA motifs of 3 nucleotides (e.g., AGC, GTA) are generated, resulting in 4^3 (instead of 4^30) possible DNA sequences due to the 3-nucleotide length:

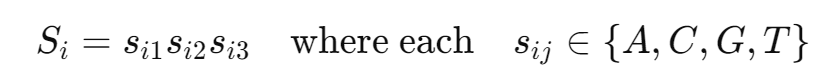

Then, 1 million different DNA sequences with lengths of 6 to 30 nucleotides are generated using these K-mers combinations:

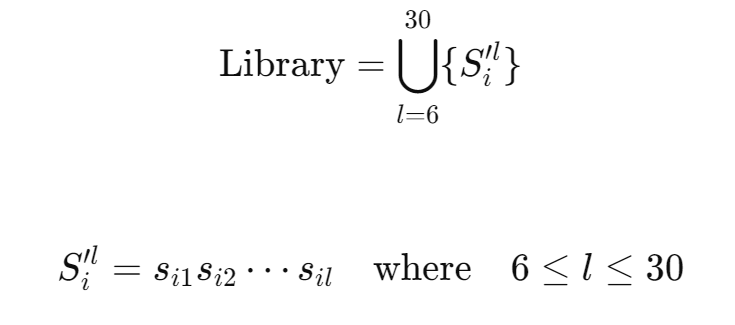

In the random DNA strands method, sequences are randomly created with lengths varying between 6 and 30 nucleotides, resulting in 1 million sequences:

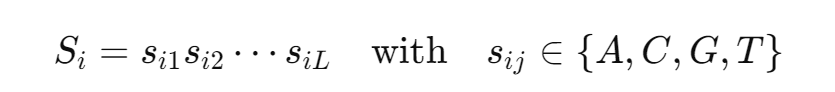

Finally, both sets of generated DNA sequences undergo clustering to select diverse sequences for further analysis, ensuring no reverse complements are duplicated:

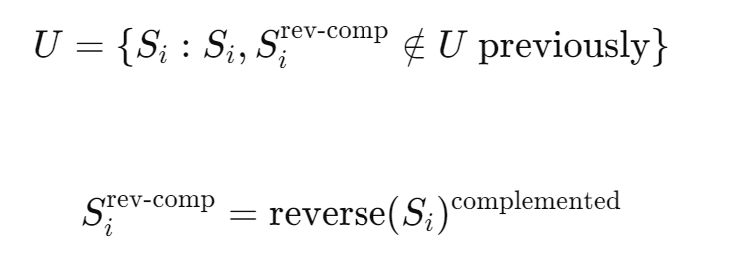

**MAFFT Clustering**

The next step in the methodology involves aligning the DNA sequences and performing clustering based on the alignment results. Pairwise similarity alignment is performed using the MAFFT algorithm. Given a set of DNA sequences S={s_1_,s_2_,…,s_n_}, where each s_i_ is a DNA sequence, the alignment A of sequences S is obtained as:

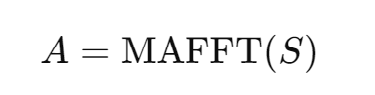

For two sequences s_i_ and s_j_ in the aligned set A, the similarity index sim_ij_ is computed as:

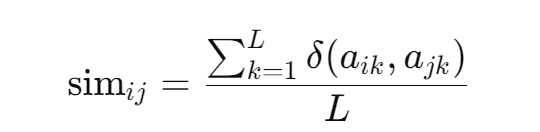

where L is the length of the sequence after alignment, δ(x,y)=1 if x=y; otherwise, δ(x,y)=0, and a_ik_ and a_jk_ are the k-th nucleotides of sequences s_i_ and s_j_ after alignment. Once the similarity indices are calculated, k-means clustering is applied to group the DNA sequences based on their similarities. k-means clustering is an iterative algorithm that partitions the data into k clusters, where each data point belongs to the cluster with the nearest mean. The optimal number of clusters k is determined using the elbow method, which involves plotting the within-cluster sum of squares (WCSS) against the number of clusters and identifying the point where the rate of decrease sharply slows, forming an "elbow. The within-cluster sum of squares (WCSS) for k clusters is calculated as:

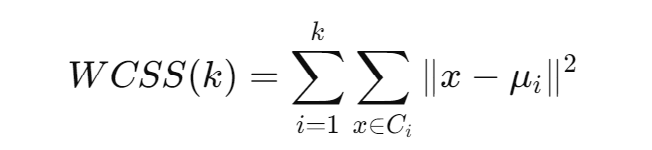

where C_i_ is the set of points (similarity indices) in cluster i, and μ_i_ is the centroid of cluster i.

**K-mers Clustering**

In the K-mers method, each DNA sequence is divided into motifs of length k. For example, for k=3, a sequence such as AGCTGGTTC is divided into overlapping K-mers: AGC, GCT, and CTG. Each sequence S_i_ is then represented as a vector of counts of each K-mers . The motif encoding for a sequence S_i_ is given by:

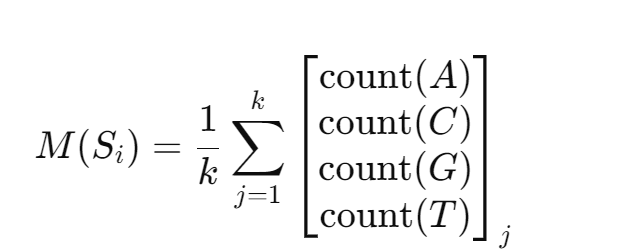

where each sequence S_i_ is divided into motifs of length k, and the counts of each nucleotide in these motifs are used to create a feature vector. After encoding the sequences into K-mers feature vectors, k-means clustering is applied to group the sequences based on these vectors. k-means clustering aims to minimize the within-cluster sum of squares (WCSS), which measures the variance within each cluster as explained in the previous k-means clustering method. This method further refines the clustering of DNA sequences by leveraging the K-mers representation and ensuring that sequences with similar K-mers compositions are grouped together, optimizing the cluster quality.

**RNA Folding Properties Clustering**

This step involves calculating DNA properties and RNA folding properties to refine the clustering process. The aim of incorporating RNA folding properties is to gain insights into the structural characteristics of RNA molecules derived from DNA sequences, which can provide additional dimensions for clustering based on both sequence and structural information. For each DNA sequence, several properties are calculated, including atomic values, electron-ion interaction potential (EIIP), molecular weights, melting temperature, and entropy. The average atomic value for a sequence is calculated as:

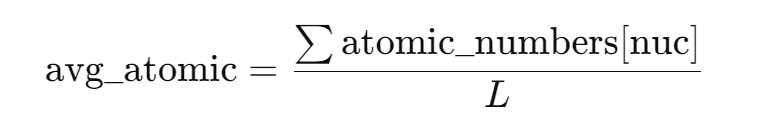

The average EIIP is calculated as:

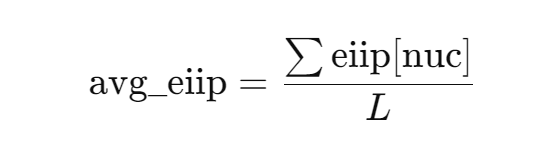

The average molecular weight is calculated as:

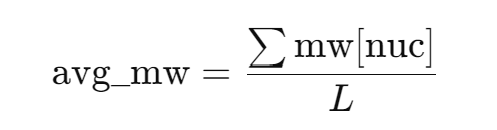

The melting temperature is calculated using the formula:

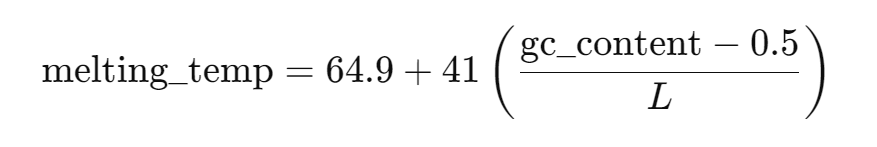

The entropy is calculated as:

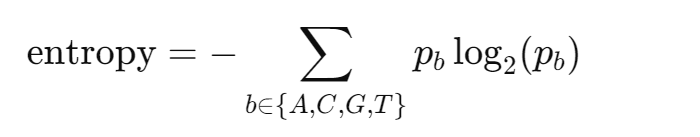

where p_b_ is the probability (or frequency) of each base b (adenine A, cytosine C, guanine G, uracil U in RNA).

**RNA Folding**

Properties such as minimum free energy, ensemble free energy, and paired probabilities are calculated via ViennaRNA folding techniques to provide insights into the potential secondary structures of RNA molecules. The minimum free energy (MFE) calculation is:

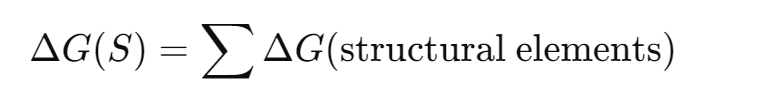

This equation computes the total Gibbs free energy (ΔG) of a secondary structure S, where the sum is over all the structural elements such as base pairs, loops, and unpaired regions. The partition function Z sums over all possible secondary structures S, considering the exponential negative of each structure’s free energy divided by the product of the gas constant R and temperature T:

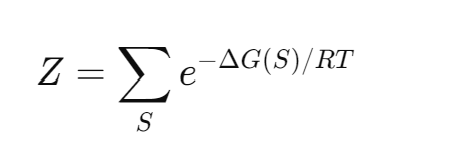

This function captures the statistical mechanics of RNA folding. The ensemble free energy is:

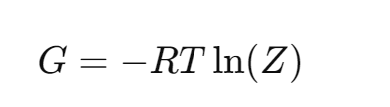

The base pairing probabilities P(I,j) that nucleotides I and j are paired, based on their specific pairing free energy ΔG(pair(I,j)) and the partition function, are calculated as:

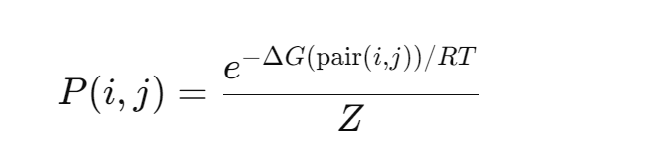

The centroid structure S_centroid_ is the most representative structure that minimizes the expected base pair distance to all other structures in the ensemble, weighted by the probability of each structure:

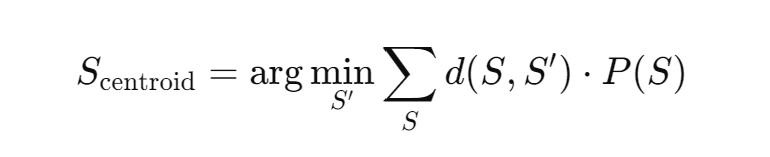

By calculating these properties, DNA sequences can be clustered based on both their inherent properties and RNA folding characteristics. Finally, k-means clustering is applied to group the sequences based on these combined properties. The within-cluster sum of squares (WCSS) for k clusters is calculated as mentioned before. This clustering method ensures that sequences with similar structural and compositional properties are grouped together, leading to more refined and biologically relevant clusters.

**Computational Methodology of the Pattern Recognition Analysis**

The next step in the methodology involves recognizing patterns in our DNA library by analyzing nucleotide proportions, positions, and K-mers frequencies. This analysis aims to identify significant patterns and motifs that may be crucial for understanding the functional and structural properties of the DNA sequences.

**Nucleotide Proportions**

For each DNA sequence S, the proportion of each nucleotide N (where N is one of {A, C, G, T} is calculated to understand the composition of the sequence. This proportion is calculated as follows:

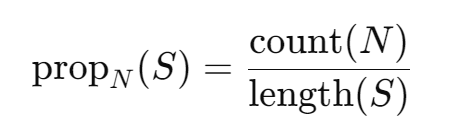

where count(N) is the number of occurrences of nucleotide N in sequence S, and length(S) is the total length of the sequence. The mean intensity for each sequence is calculated and grouped by nucleotide proportions. This helps in understanding the average signal intensity associated with different nucleotide compositions, providing insights into the overall characteristics of the sequences.

**AT and GC Content**

To further characterize the sequences, the AT and GC content are calculated. The AT content is the sum of the proportions of adenine (A) and thymine (T), while the GC content is the sum of the proportions of guanine (G) and cytosine (C):

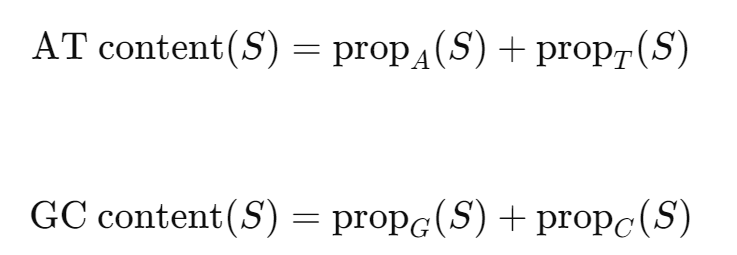

**Position Analysis**

The next phase of the methodology involves analyzing the positional effects of nucleotides within the DNA sequences. This step aims to understand how the position of each nucleotide within a sequence influences the intensity response, providing insights into positional dependencies and patterns. Given a set of DNA sequences S={DNA_1_, DNA_2_, ..., DNA_n_} and their corresponding intensity responses R={R_1_, R_2_, ..., R_n_}, the first step is normalization. The response for each nucleotide in a sequence is normalized by the length of that sequence:

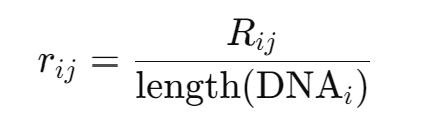

where r_ij_ is the normalized response for the j-th nucleotide in the i-th sequence. Next, the normalized responses for each nucleotide at each position across all sequences are aggregated and averaged. This helps in calculating the mean response for each nucleotide at specific positions:

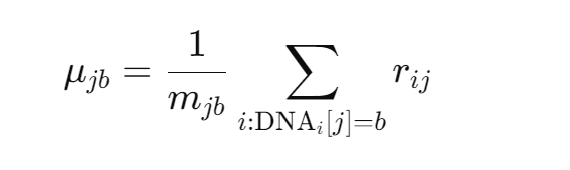

where μ_jb_ is the mean normalized response for nucleotide b at position j, and m_jb_ is the count of nucleotide b appearing at position j across all sequences.

To visualize the positional effects, a matrix M is constructed where each element M_jb_ is the mean response μ_jb_. This matrix is then used to create a heatmap, which visually represents the normalized and averaged responses of nucleotides at various positions in the DNA sequences. The heatmap helps in identifying positional patterns and hotspots that are critical for further analysis.

**K-mers analysis**

To recognize patterns in our DNA library, we utilize K-mers analysis. This methodology involves extracting K-mers from DNA sequences, calculating their frequency and mean intensity response. For each DNA sequence in the dataset, we extract K-mers of a specified length k. For instance, if k=3, a sequence such as "AGCTGGTTC" is divided into overlapping K-mers: AGC, GCT, CTG, etc. This is performed using:

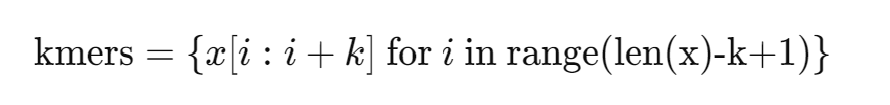

where x is a DNA sequence. The frequency of each K-mers is calculated by counting its occurrences across all sequences. This results in a frequency F:

F(K-mers )=count(K-mers )

For each K-mers , we calculate the mean intensity response. If R represents the intensity response for a sequence containing the K-mers , the mean intensity μ_kmer_ is:

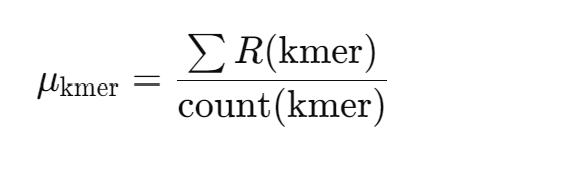

The frequency of each K-mers is calculated, and the K-mers with high frequency and high response are selected as the most effective K-mers in our dataset.

**Computational Methodology of Deep Learning**

In our research, we used a combination of deep learning methods, including convolutional neural network (CNN), graph convolutional network (GCN), and graph neural network (GNN) models, to predict the behavior of DNA sequences in our library. The goal of this study was to classify DNA sequences based on their photoluminescence (PL) intensity changes in the (7,6) and (9,4) nanotube chiralities in response to glucose. The workflow included data preprocessing, DNA sequence embedding, feature extraction, and neural network training to classify the fluorescence response of the two chiralities. After training, we implemented an optimized model using majority voting across the three methods to predict the response of new DNA sequences, which were then experimentally validated for glucose detection.

To initially process DNA sequences, embedding was performed either using 5-eye encoding to create a matrix or using DNA encoding as a graph. Working with these methods in parallel, enabled us to extract a maximum of information from the DNA and to preserve K-mer information. We also applied various feature extraction techniques, namely graph2vec, convolution, pooling, and flattering layers, to capture the most relevant information from the DNA sequences for subsequent training of neural networks using various combinations of layers and neurons (Figure 3A). Additionally, k-fold cross-validation was used to increase the reliability of models and prevent overfitting or underfitting. Finally, all hyperparameters (combinations of feature extraction and neural network properties, as well as k-fold) were optimized using the Optuna model (Table S4).

**Data Preprocessing**

Photoluminescence (PL) spectra were acquired using a custom-built near-infrared optical microscope system. Excitation wavelengths of 660 nanometers and 730 nanometers were employed to selectively excite specific chiralities, with (7,5) and (7,6) excited at 660 nanometers, and (10,2), (9,4), (8,6), and (8,7) excited at 730 nanometers. The emission spectra, ranging from 900 to 1400 nanometers, were collected using an IsoPlane SCT-320 spectrometer equipped with a NIRvana 640 ST InGaAs camera. The raw PL data were processed using Python scripts that implemented background noise correction, Savitzky-Golay filtering, and segmentation of the dataset according to the excitation wavelength. To resolve overlapping spectral features, Lorentzian peak deconvolution was performed on each spectrum. Nonlinear least-squares fitting was used to optimize parameters including full width at half maximum, peak position, and intensity. This approach enabled the extraction of precise peak characteristics for each chirality and facilitated more accurate subsequent analyses. For ssDNA-SWCNT experiments involving glucose detection, additional scripts were used to calculate statistical measures such as mean values, standard deviations, peak shifts, and intensity changes across multiple DNA sequences. Among the analyzed chiralities, (7,6) and (9,4) exhibited significant intensity changes in response to glucose and were therefore selected for further study and used as output variables in model training.

**CNN Model**

In the CNN approach, DNA sequences will be classified based on their photoluminescence (PL) intensity changes using a convolutional neural network. The process involves several steps: data preprocessing, feature extraction, model training, and evaluation. The dataset is including X represents the DNA sequences and y the PL intensity changes. For classification purposes, the PL intensity values are binarized into two classes based on a threshold of 12. The distribution of PL intensity values and their binarized classes are visualized to understand their distribution. To convert DNA sequences into a suitable form for the CNN, one-hot encoding is employed. Each sequence is encoded into a matrix of size fixed_length×5 where each row represents a nucleotide (A, G, C, T, N). Given a DNA sequence S, one-hot encoding maps each nucleotide to a unique binary vector:

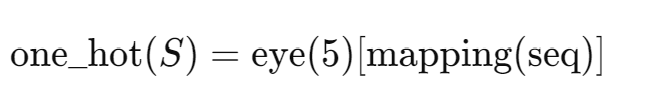

Where mapping converts nucleotides to indices and eye(5) creates a one-hot encoded matrix. The encoded sequences are reshaped to a 4D tensor suitable for CNN input. The CNN architecture is optimized using Optuna for hyperparameter tuning. The architecture consists of convolutional layers followed by pooling layers, flattening, and dense layers. The hyperparameters optimized include the number of filters in the convolutional layers, kernel sizes, dense units, dropout rates, learning rates, pooling types, and optimizers. The architecture of the CNN can be summarized as:

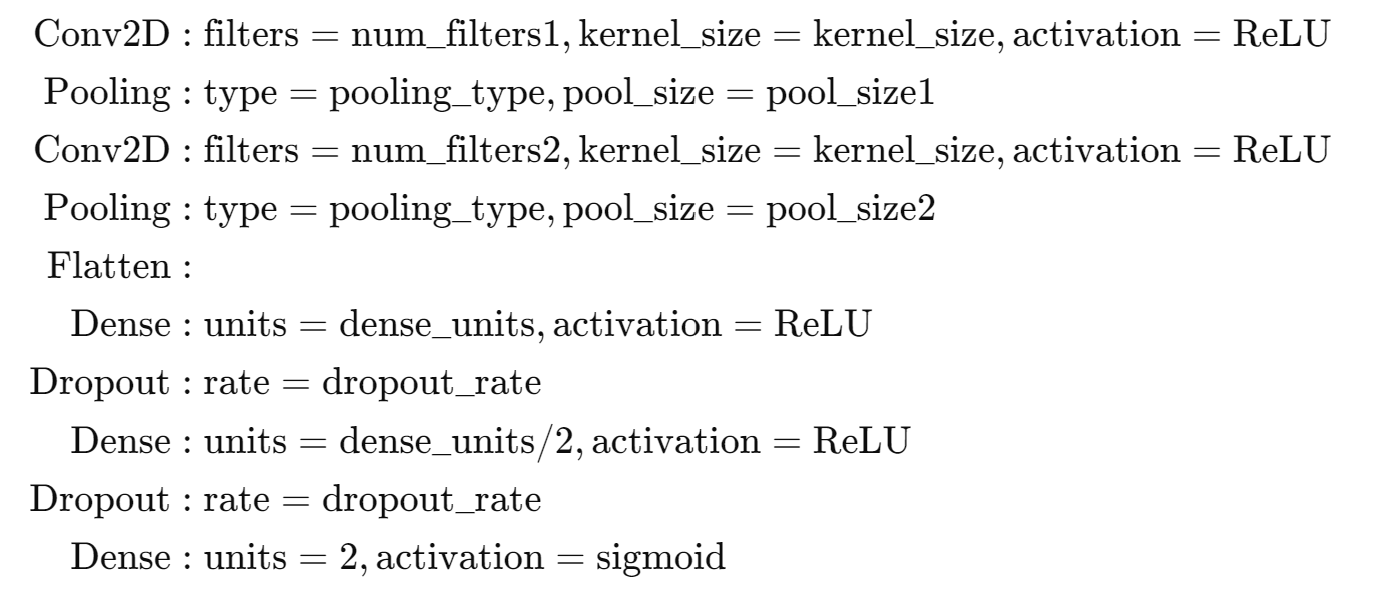

The model is compiled using the selected optimizer (Adam, SGD, or RMSprop) with the chosen learning rate and binary cross-entropy loss. Training is monitored with early stopping based on validation loss. The objective function for Optuna is defined to minimize the loss over k-fold cross-validation:

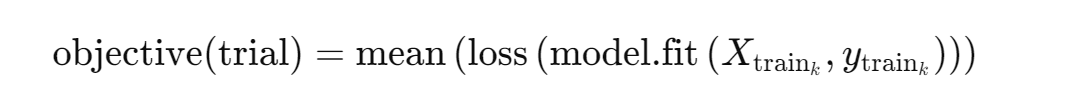

where the hyperparameters for each trial are sampled from predefined ranges and distributions. The best hyperparameters are identified and saved for future use. K-fold cross-validation is used to ensure that the model generalizes well to unseen data by splitting the dataset into multiple folds and training the model on different subsets of the data. K-fold cross-validation involves dividing the dataset into k equally sized folds. For each fold, the model is trained on k-1 folds and tested on the remaining fold. This process is repeated k times, with each fold being used as the test set once. The results from each iteration are averaged to produce a single performance metric. In this study, the number of folds best_fold is determined based on optimization. The model's performance on the test set is evaluated using various metrics such as accuracy, precision, recall, F1 score, and ROC AUC score.

**GNN Model**

By using a GNN model, DNA sequences are classified based on their photoluminescence (PL) intensity using graph-based features and neural network models. The process was done similarly to the CNN model in different steps: data preprocessing, feature extraction, model training, and evaluation. The dataset includes X, which represents the DNA sequences, and y, the PL intensity values. The data is classified into high and low response classes, ensuring a balanced dataset in both classes. To transform the DNA sequences into a suitable form for GNN models, Graph2Vec embeddings are used. Each sequence is converted into a graph based on K-mers (substrings of length k) and then embedded into a fixed-dimensional vector. Given a DNA sequence S={s_1_,s_2_,…,s_n_} and a K-mers size k:

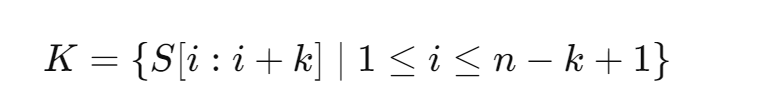

A directed graph G=(V,E) is constructed, where V is the set of K-mers and E is the set of directed edges between overlapping K-mers. Graph2Vec is then used to convert G into a fixed-dimensional vector v∈Rd. A neural network is trained to classify the DNA sequences based on their Graph2Vec embeddings. The dataset is split into training and test sets. The neural network model *f* is described as follows: the input layer x∈Rd, hidden layers

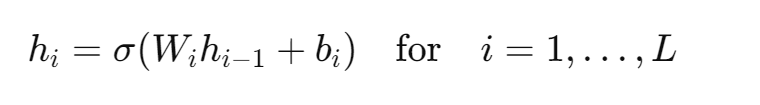

where σ is the activation function, W_i_ are the weight matrices, and b_i_ are the biases, and the output layer:

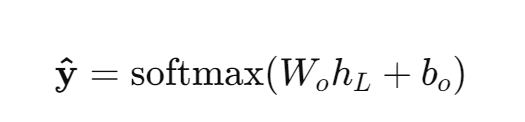

The model is trained using backpropagation to minimize the loss function.

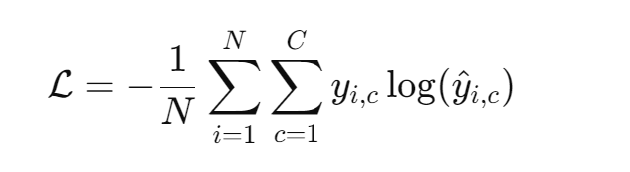

where N is the number of samples, *C* is the number of classes, y*_i_*_,c_ is the true label, and y^_i,c_ is the predicted probability. The optimization of hyperparameters and k-fold cross-validation was done as described in the CNN model procedure.

**GCN Model**

Similar to the previous method, this study aims to classify DNA sequences based on their photoluminescence (PL) intensity using a Graph Convolutional Network (GCN). The process involves multiple steps: data preprocessing, feature extraction, model training, and evaluation. The optimization of hyperparameters and k-fold cross-validation was done as described in the CNN model procedure. To convert DNA sequences into a suitable form for the GCN, K-mers are employed. Each sequence is divided into overlapping K-mers, which are then used to construct a directed graph where each unique K-mers is a node. The nodes are connected by directed edges if consecutive K-mers overlap. Given a DNA sequence *S* and a K-mers size *k*:

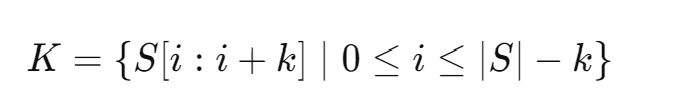

A directed graph G=(V, E) is constructed, where *V* is the set of K-mers and *E* is the set of directed edges between overlapping K-mers. The adjacency matrix *A* and the feature matrix *X* for the graph are then created. The feature matrix is derived by one-hot encoding each K-mers. The GCN architecture consists of graph convolutional layers followed by a dense layer for classification. The architecture can be summarized as:

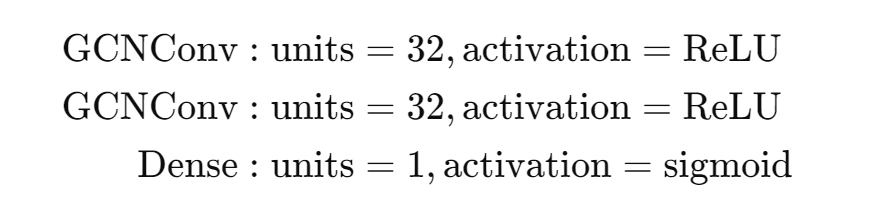

The model is compiled using the Adam optimizer with binary cross-entropy loss. The objective function for the GCN can be written as:

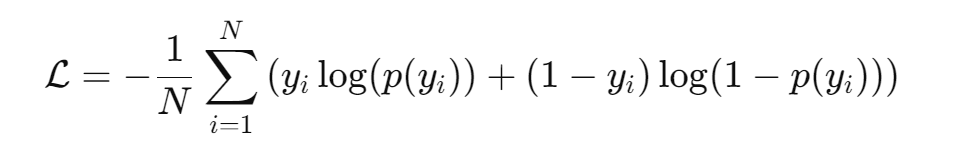

where N is the number of samples, y_i_ is the true label, and p(y_i_) is the predicted probability.

The GCN model is trained using mini-batches of the graph data. Each batch consists of a subset of graphs, and the corresponding labels are used for training. The training process involves iterating over the dataset for a specified number of epochs. The evaluation of the model involves calculating accuracy metrics on the test set. The GCN model is implemented using the Spektral library, which provides tools for handling graph data and building graph neural networks in TensorFlow. This methodology ensures a robust approach to classifying DNA sequences using graph-based features and a Graph Convolutional Network, providing insights into the relationship between DNA sequence structure and PL intensity.

**Model evaluation parameters**

A variable called the confusion matrix (COF) was defined to evaluate different machine learning classification models. This matrix compares the actual values (experimental data) with the predictions made by the model. An effective model has high true positive (TP) and true negative (TN) rates, and low false positive (FP) and false negative (FN) rates.

True Value

|  | Positive  Predicted Class | Negative |
| --- | --- | --- |
| Positive | True  Positive | False Negative |
| Negative | False  Positive | True Negative |

**Confusion matrix**

True Positive (TP) indicates that the model correctly predicted a 'Yes' when the actual value was also 'Yes.' True Negative (TN) means the model correctly predicted 'No' when the actual value was 'No.' False Positive (FP) occurs when the model predicted 'Yes,' but the actual value was 'No.' False Negative (FN) happens when the model predicted 'No,' but the actual value was 'Yes.'

The following are the definitions for true positive rate, true negative rate, and accuracy:

- **True Positive Rate (TPR)**: TP / (TP + FN)
- **True Negative Rate (TNR)**: TN / (TN + FP)
- **Accuracy (Acc)**: (TP + TN) / (TP + FP + TN + FN)

The Receiver Operator Characteristic (ROC) plot is a graphical representation of a binary classifier's diagnostic performance. It plots the true positive rate against the false positive rate to form the ROC curve. ROC curves are useful for comparing several classifiers by combining their performance into a single metric. The area under the ROC curve (AUC), ranging from 0 to 1, is a common metric for prediction precision. It represents the likelihood that a randomly chosen positive case is ranked higher than a randomly chosen negative case, or the Wilcoxon rank-sum statistic for two samples

**Model Prediction**

In the next step, we generated a new library of DNA sequences from our data analysis to use as input for the machine learning model prediction. Approximately 100,000 DNA sequences were generated and then evaluated using our optimal model. Sequences with a high probability of high-intensity response were selected for experimental validation to identify new sequences responsive to glucose.

**Generate DNA Sequence from position analysis**

To generate new DNA sequences based on intensity response matrices from two datasets, I76(j,b) and I94(j,b), where j denotes the position and b the nucleotide, the following steps are undertaken. The goal is to create a set of sequences S with specific properties defined by the response data. Given:

- I76(j,b) and I94(j,b): Intensity response matrices for two datasets.
- n: Total number of sequences to generate.
- L: Randomly chosen sequence length within predefined bounds.
- S: Set of all generated sequences.

Each sequence s is defined as s={b_1_,b_2_,...,b_L_}, where each nucleotide b_i_ is selected based on the following criteria:

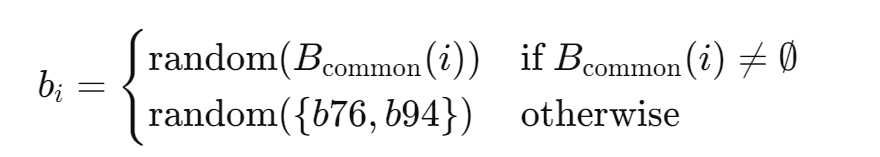

where B_common_(*i*) is the set of nucleotides with top responses in both I76 and I94 at position *i*, and b76 and b94 are the nucleotides with the highest response at position *i* in I76 and I94, respectively. The overall response for a generated sequence s is calculated by summing the intensity responses from both datasets:

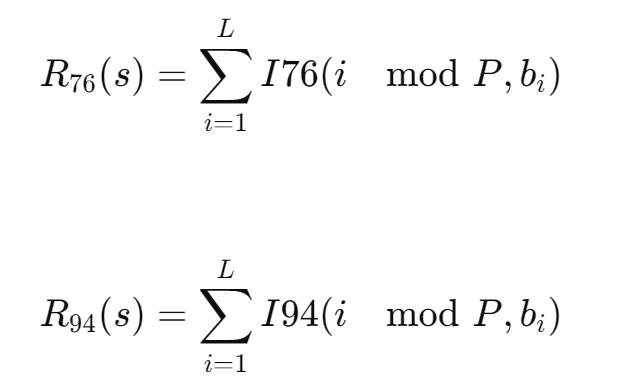

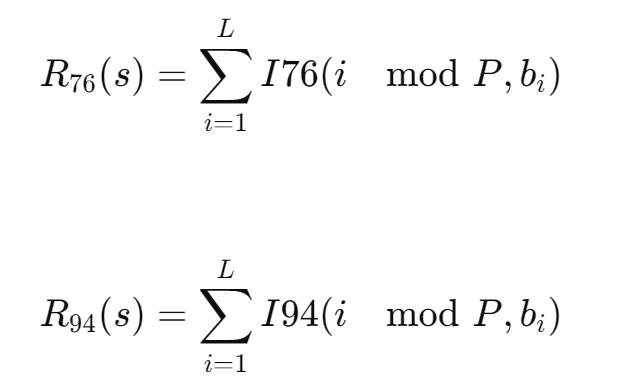

Where P is the number of positions covered by the datasets. This approach ensures that the generated sequences are informed by the top intensity responses from both datasets, providing a robust method for creating sequences with desirable properties. By selecting nucleotides based on high response values and commonality between datasets, the generated sequences are optimized for further experimental and analytical purposes.

**Generating New Sequences from K-mers Analysis**

Based on the identified high-response K-mers, new DNA sequences are generated by combining these K-mers in various ways. This generation process ensures that the new sequences retain the desirable properties observed in the high-response K-mers. Essentially, K-mers with a high response in our dataset are combined with high frequency to create DNA sequences of different lengths.

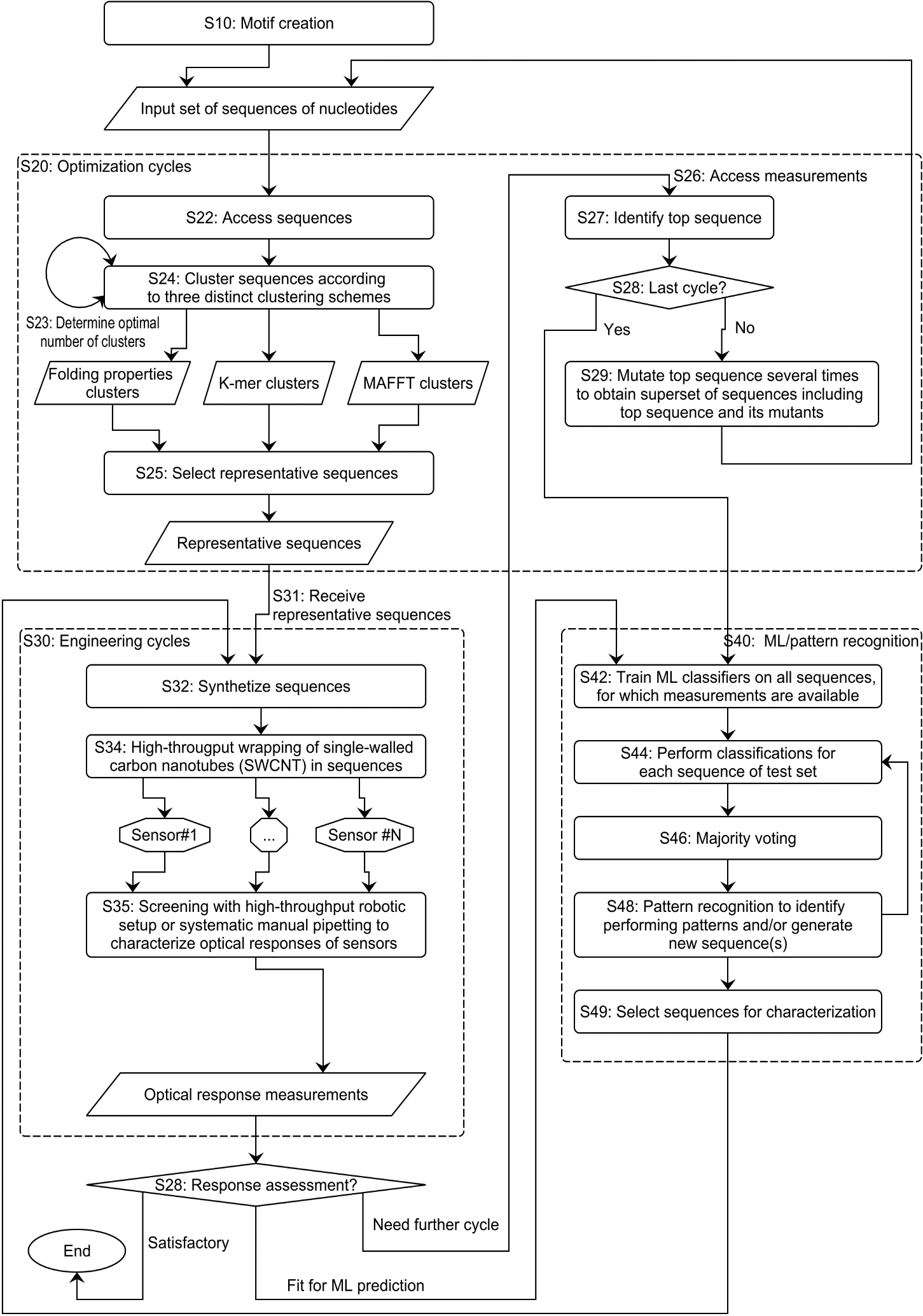

**Fig. S1 - Flowchart illustrating the strategies for applying computational clustering methods and deep learning to discover DNA sequences that interact with small molecules**. The diagram outlines the steps involved in identifying candidate DNA sequences, refining their performance using deep learning, and applying this methodology to cases where the interaction mechanisms between DNA and specific target molecules are unknown. This approach is particularly useful for discovering sequences that respond to challenging small molecules with unclear interaction pathways. Some of these iterative cycles are used to find DNA sequences that can interact with glucose, but as indicated in the flowchart, this process can be repeated as needed based on the specific response required for target detection.

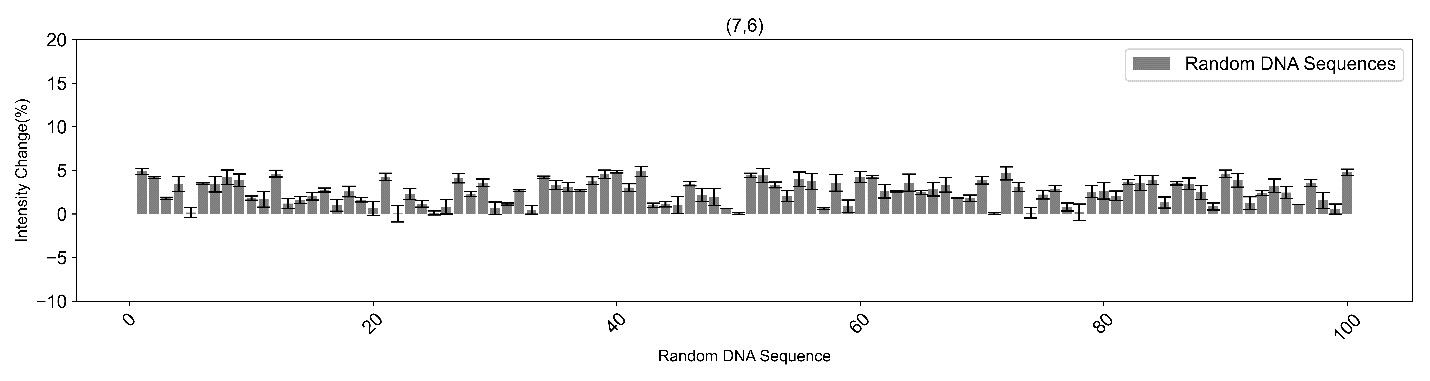

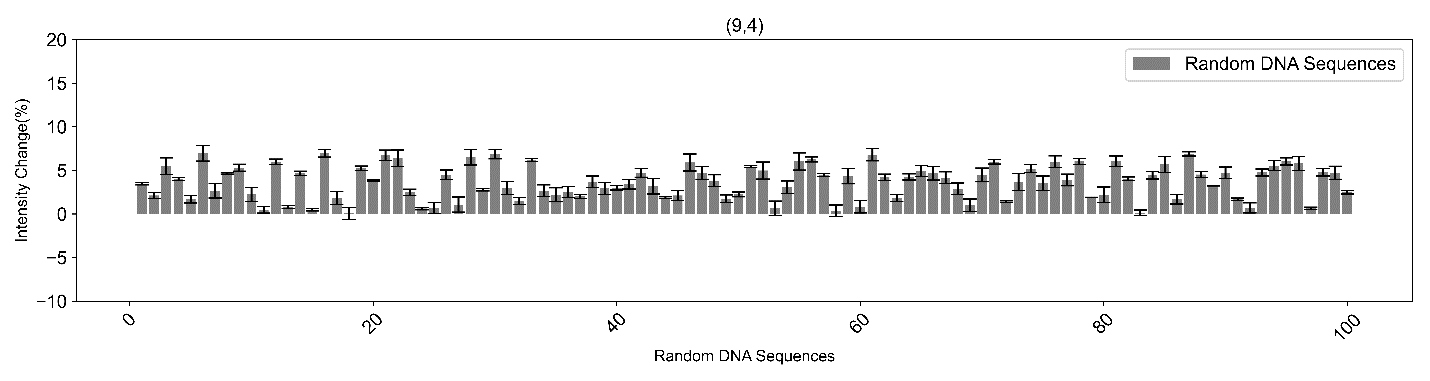

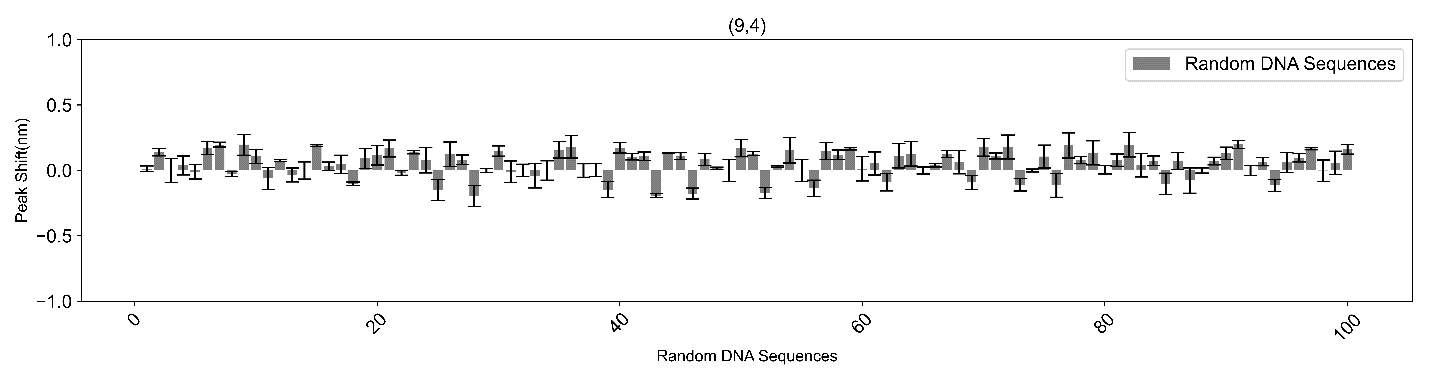

**Fig. S2 - Experimental results of random DNA screening for glucose detection.** Screening results of different random DNA sequence creations for making an ssDNA-SWCNT sensor for glucose detection at 7.5 mM. This data represents most of the random DNA sequences created and screened for glucose detection. After several attempts at generating different libraries of random DNA sequences, no sequence was found that resulted in an intensity response of more than 10% or a peak shift greater than 0.5 nm. Intensity changes of the (7,6) and (9,4) chiralities, as well as the peak shift of the 9,4 chirality of SWCNTs were selected are reported, as they showed the highest sensitivity to glucose.

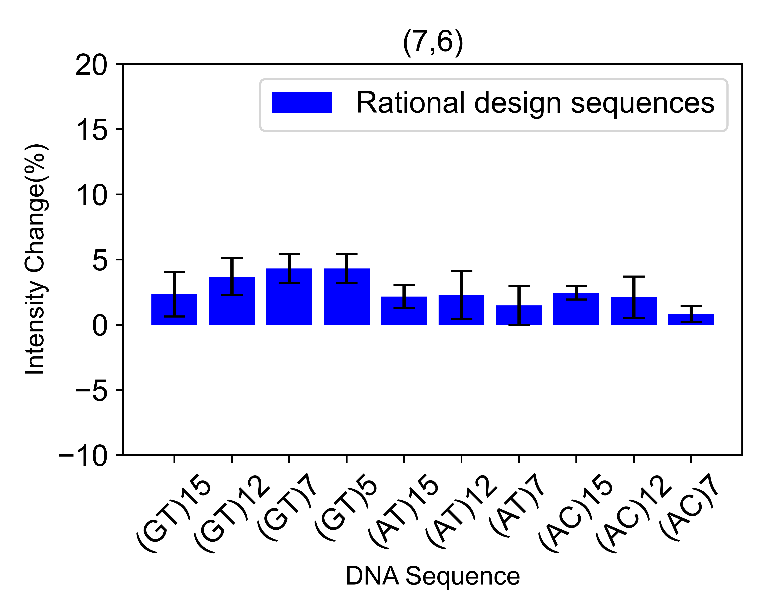

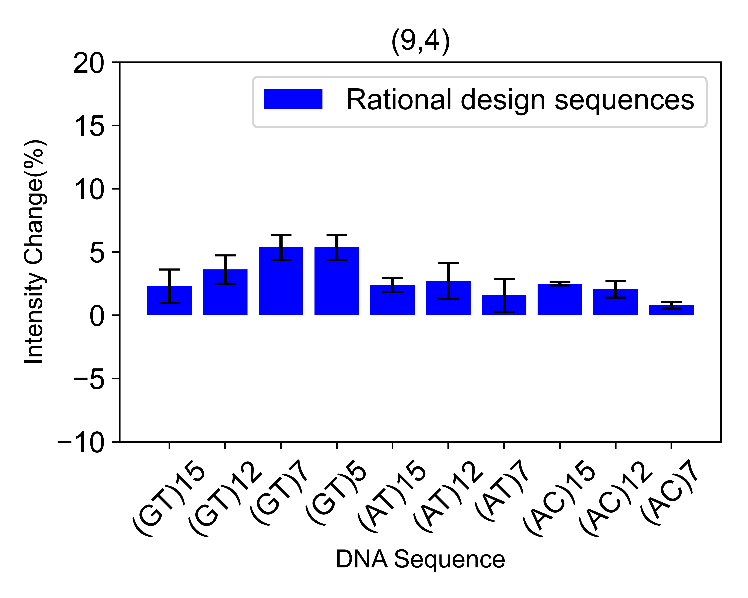

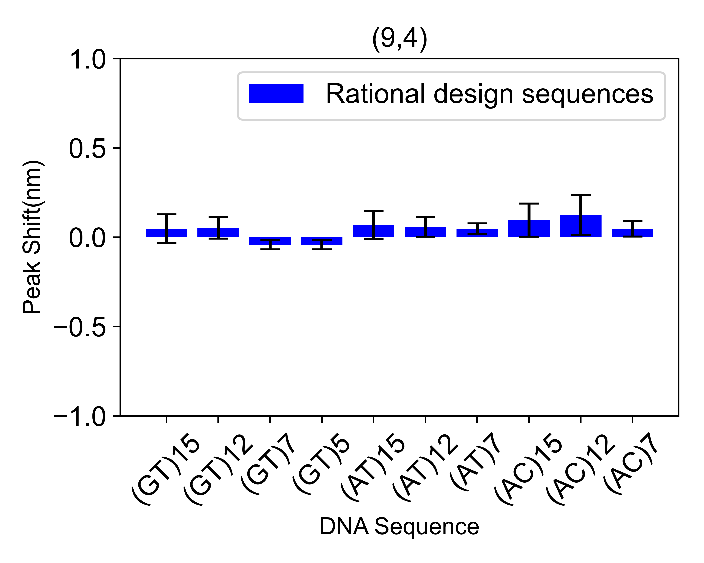

**Fig. S3 – Experimental results of rational design screening for glucose detection:** Screening results of different ssDNA-SWCNT for glucose detection at 7.5 mM: The effect of the lengths on classic and well-studied (GT)x, (AC)x, and (AT)x sequences by rational design shows that there is no response improvement by reducing the length for these sequences. Therefore, the sequence length, as well as the proportion and position of each base, plays a key role in engineering a sensitive sensor. Intensity changes of the (7,6) and (9,4) chiralities, as well as the peak shift of the 9,4 chirality of SWCNTs were selected are reported, as they showed the highest sensitivity to glucose.

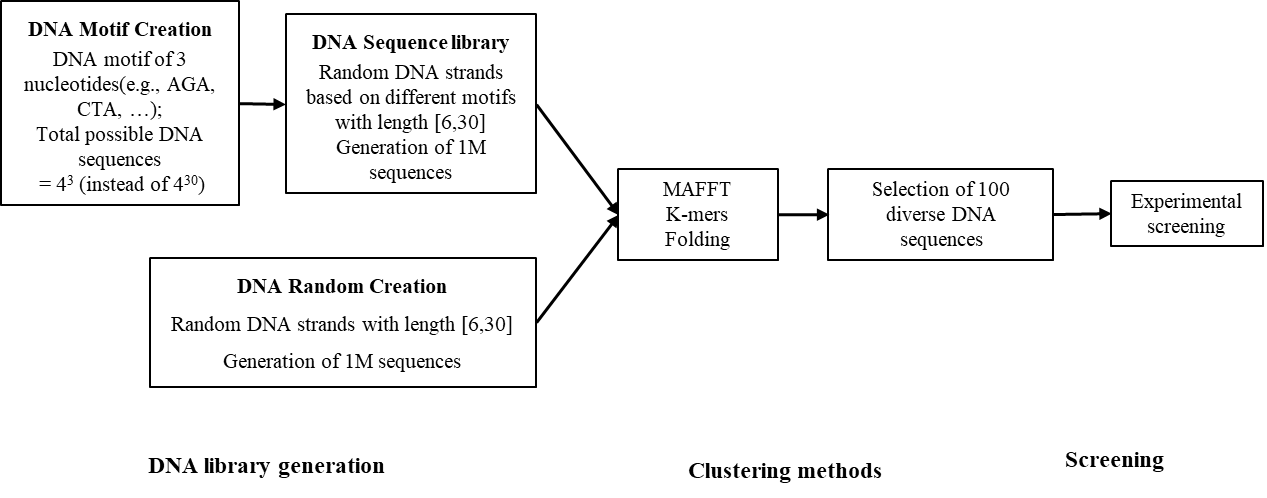

**Fig. S4- Workflow of DNA library generation, clustering and screening**: Generating DNA sequences through motifs and random combinations to encompass all possible sequence variations in the entire search space. A first library of 1 million ssDNA sequences was randomly generated for ssDNA strand lengths ranging from 6 to 30 nucleotides. A second library of 1 million ssDNA sequences was created by first generating all possible 3-nucleotide ssDNA motifs (eg. ACT, CTA, GGA, ...), and then randomly combining these motifs to form sequences ranging in length from 6 to 30 nucleotides. The whole library was clustered using different methods including MAFFT, K-mers , and Folding (see main text) to select diverse sequences. The selected sequences were then chosen for the screening step, which involves measuring the fluorescence peak shift or intensity change of different SWCNT chiralities wrapped with different DNA sequences, to identify the sequences that show a response to glucose.

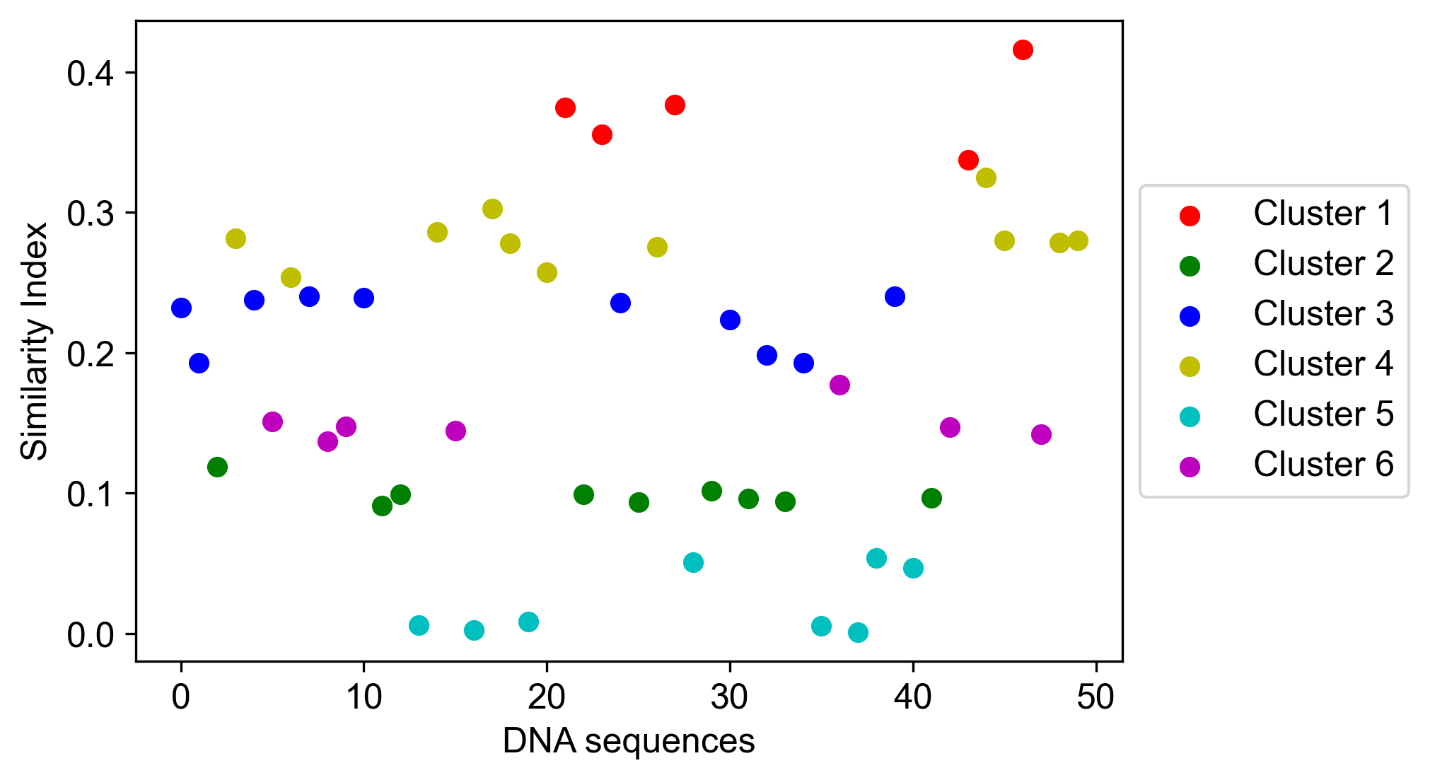

**Fig. S5- Similarity-based (MAFFT) DNA Clustering:** Clustering of DNA sequences illustrated with an example of 50 DNA sequences for the MAFFT method. In this method, different DNA sequences were compared based on their alignment, and a similarity index was calculated for each sequence. The DNA sequences were then clustered by k-means clustering based on the similarity index.

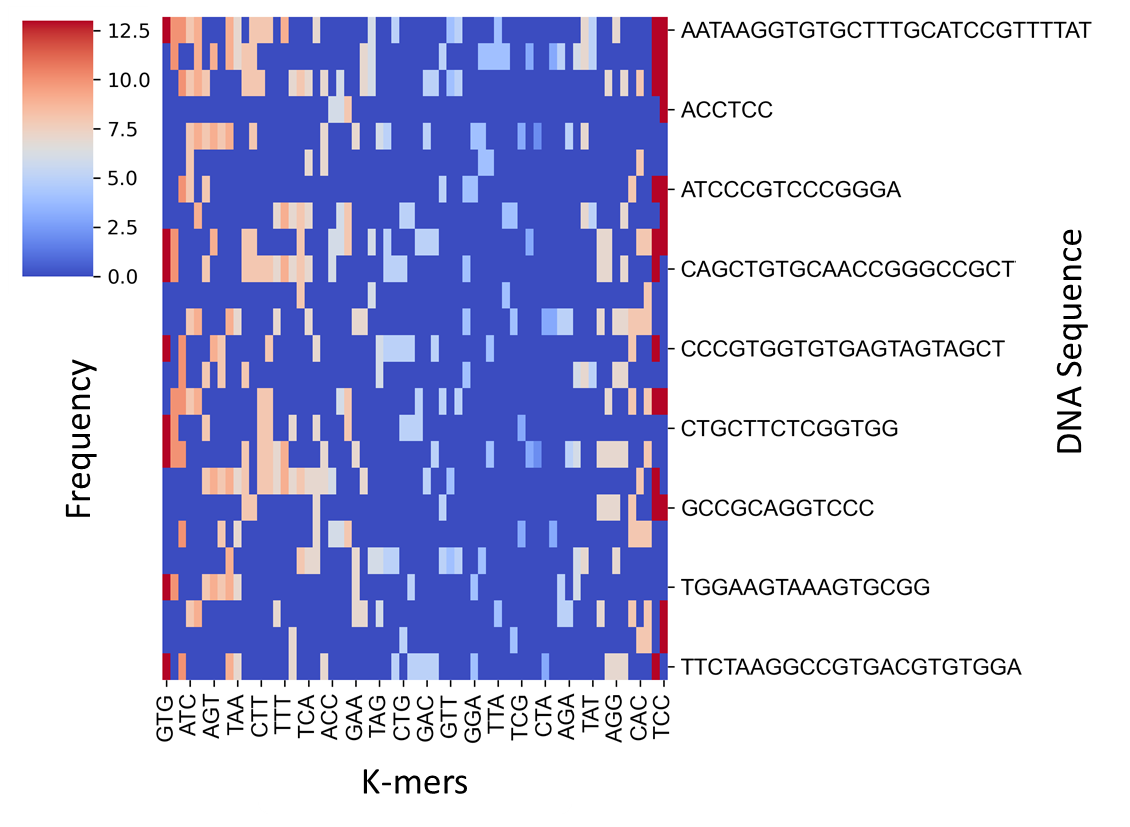

**Fig. S6 - Pattern-based (K-mers) DNA Clustering:** Clustering of DNA sequences illustrated with an example of 50 DNA sequences for the K-mers method. In this method, different DNA sequences were compared based on their 3-mer combinations and the frequency of each 3-mer within the sequences. k-means clustering was then applied based on the composition and frequency of the 3-mers within each sequence.

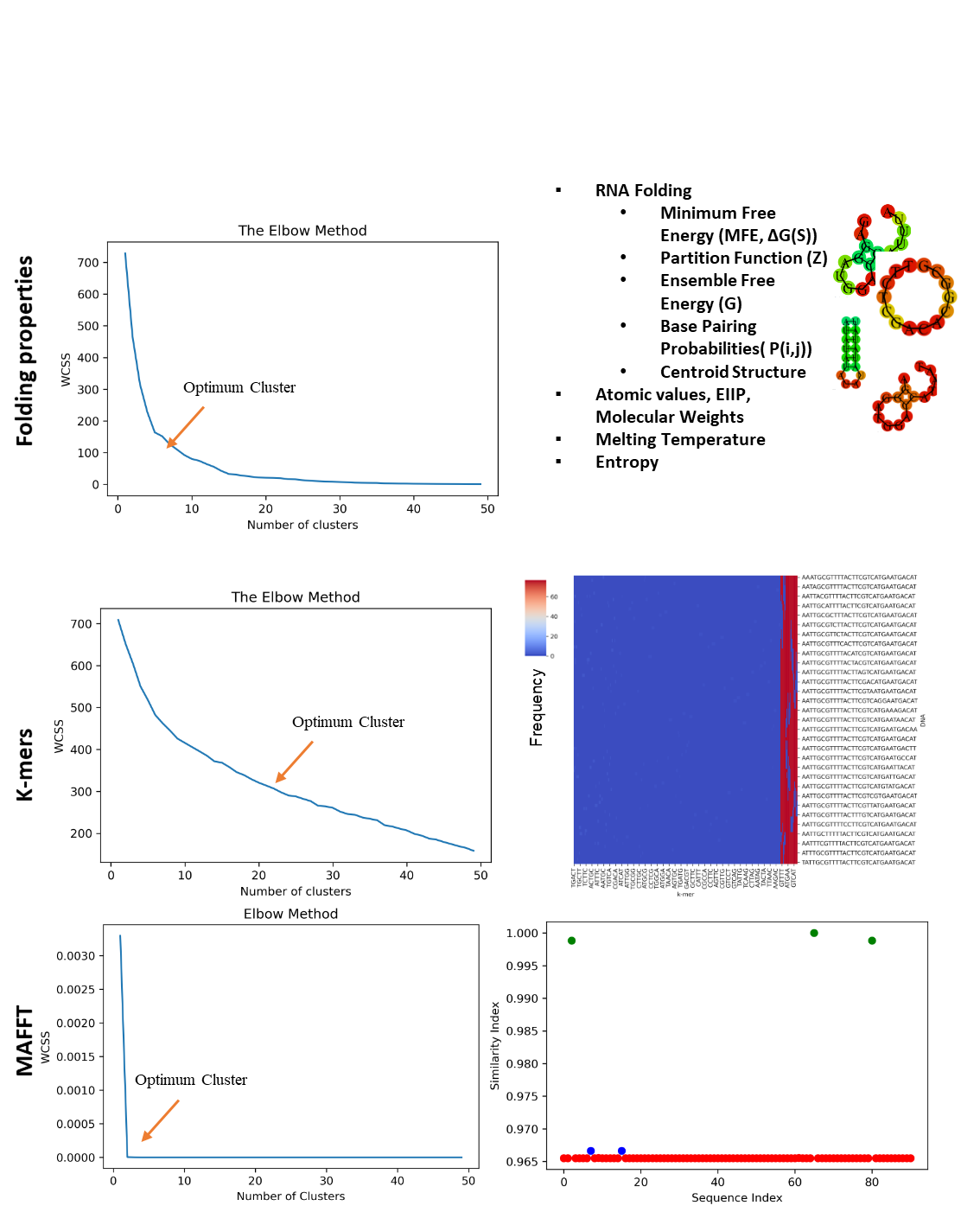

**Fig. S7- DNA Clustering methods applied to a small library of 90 DNA sequences derived by mutations from the same parent sequence.** The elbow method shows the optimum number of clusters for each clustering method (left plots). On the right side, the visualization distinguishes the different DNA sequences. The results show that the MAFFT method, based on DNA alignment, can not subcluster sequences qith high sequence similarity effectively. In contrast, other methods show more detailed distinctions, as the K-mers pattern and folding properties inside the sequence change significantly, even with only one base change.

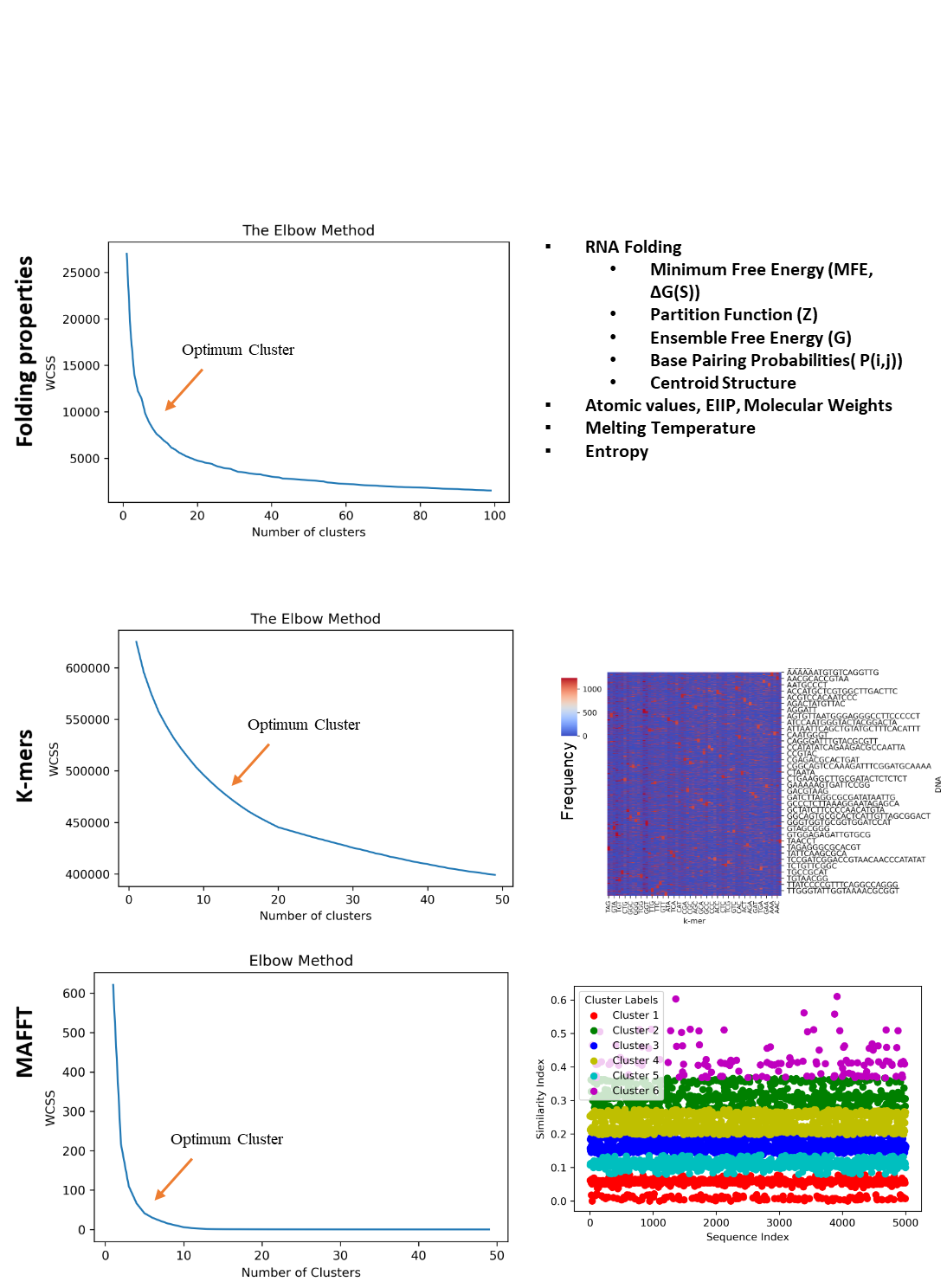

**Fig. S8- DNA Clustering methods applied to a diverse library of 1 million DNA sequences** The elbow method and visualization are shown for different clustering methods. Note that the figures on the right display clustering for 5,000 DNA sequences, as it was not feasible to display all 1 million sequences. Results reveal that the clustering methods are working independantly to seperate the DNA sequence space into various sections, making their parallel application highly complimentary.

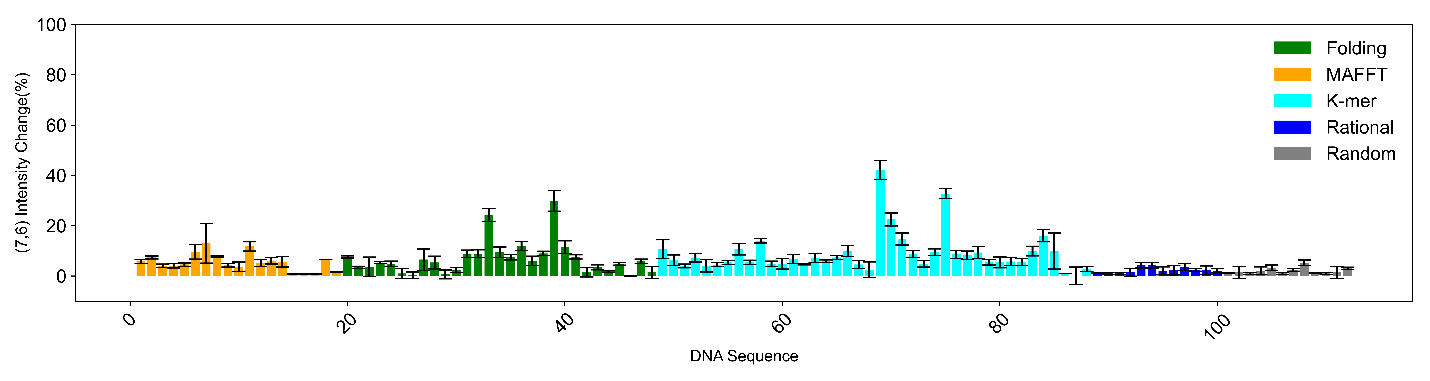

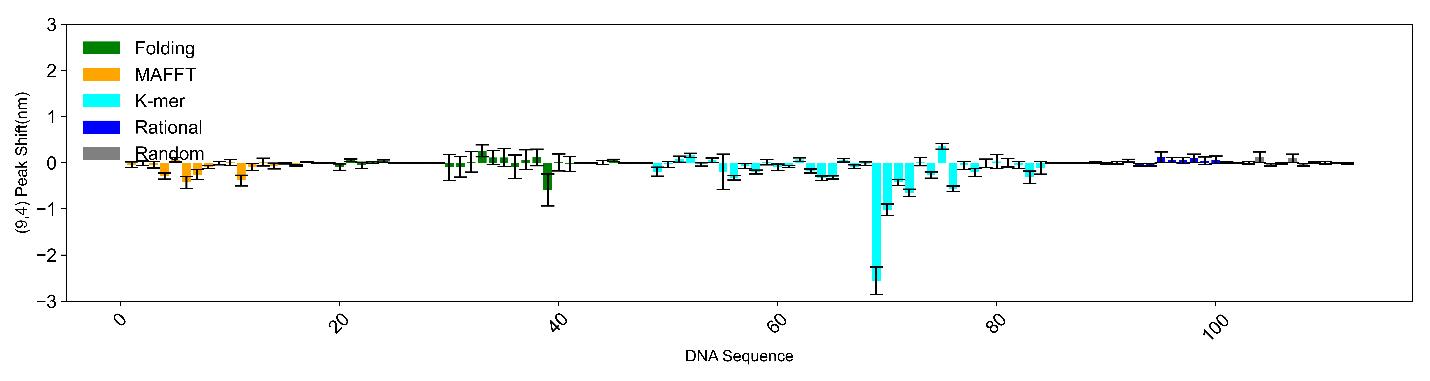

**Fig. S9 - Experimental response of DNA-SWCNT sensors to glucose using diverse ssDNA sequences selected through clustering.** Experimental (9,4) peak shift and (7,6) intensity change in response to glucose detection at 7.5 mM by ssDNA-SWCNTs resulted in finding one sequence with a -2.5 nm peak shift at (9,4) and a 40% intensity change at (7,6). The results, categorized based on different clustering methods, rational design and a random sequence library, demonstrate the ability of successful clustering to find a responsive sequence for glucose detection, which was not possible with rational and random design.

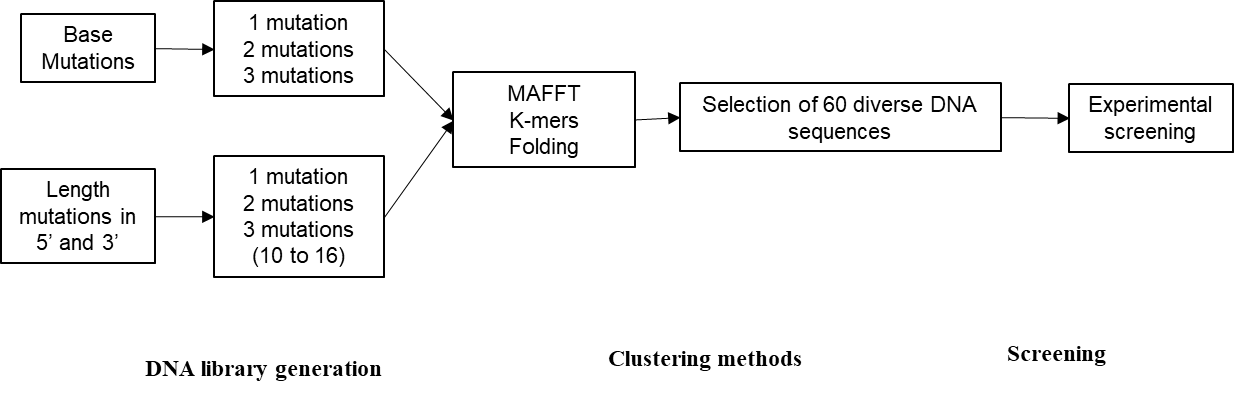

**Fig. S10- Workflow of DNA mutant library generation, clustering and screening:** Different mutations were introduced to generate a large library of mutants, potentially having a strong response to glucose. Mutations were made in the base sequence (1, 2, or 3 mutations) and by adding bases at each extremity of the ssDNA sequence (1, 2, or 3 bases at each end). These two libraries of mutant sequences were then clustered using the MAFTT, K-mers, and Folding methods to select sequences to be experimentally tested, maximizing diversity for the screening.

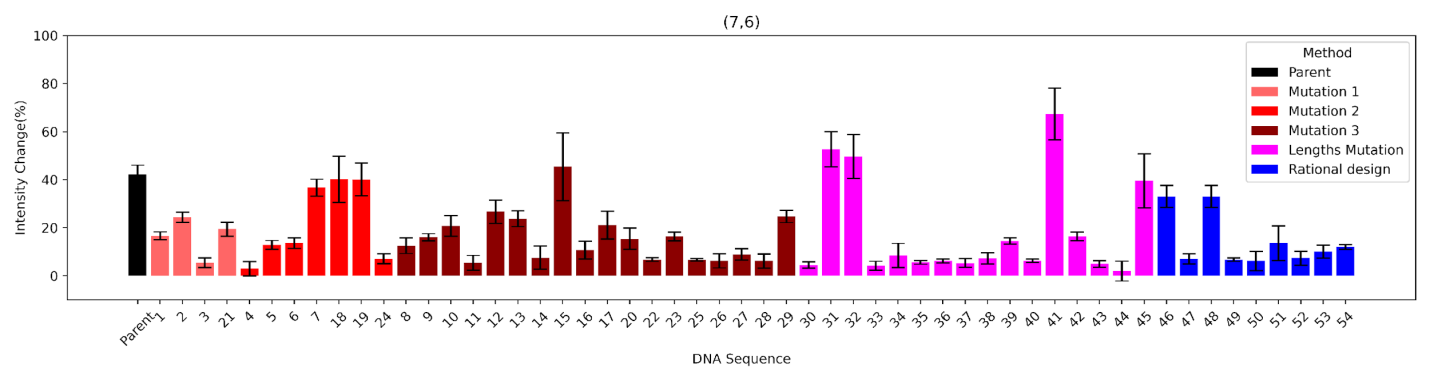

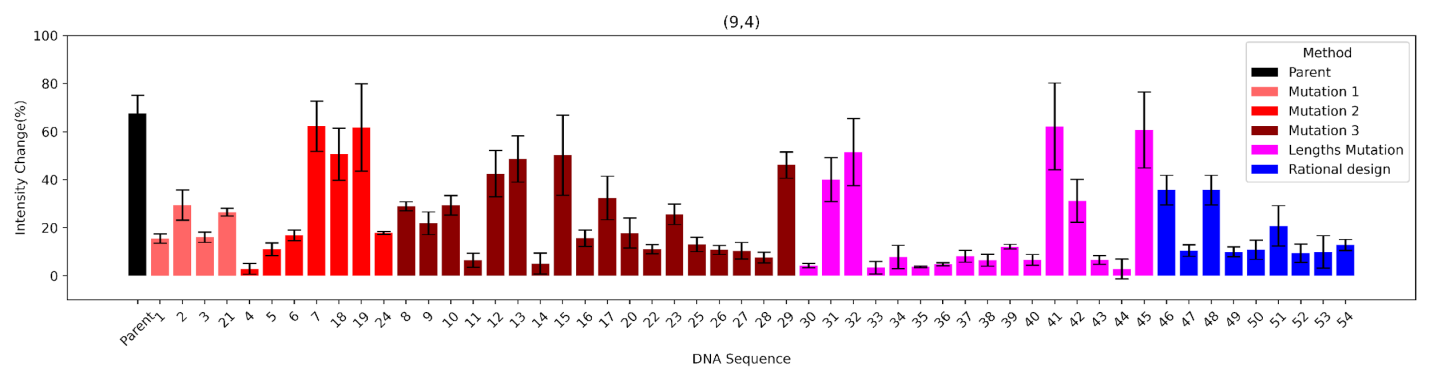

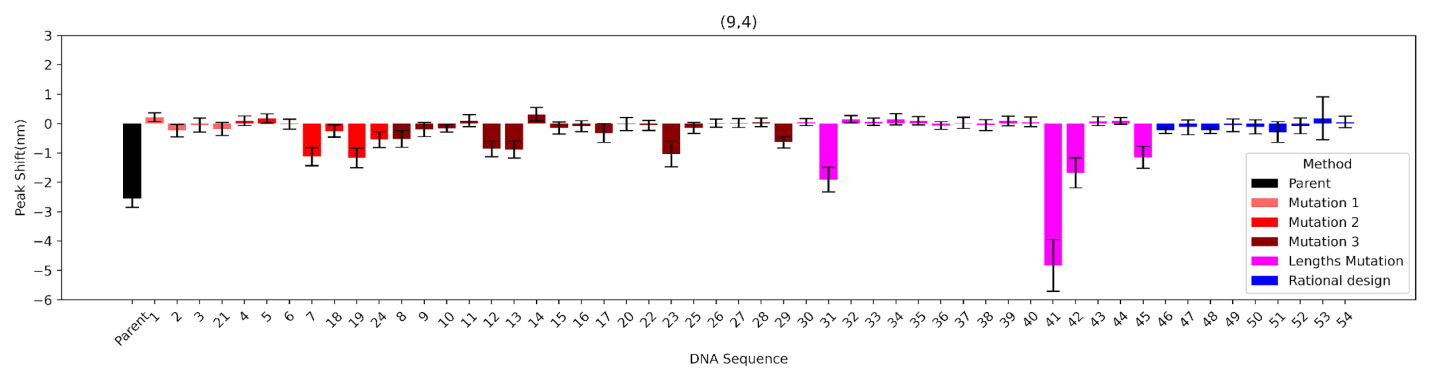

**Fig. S11 - Experimental response of DNA-SWCNT sensors to glucose using diverse ssDNA sequences selected through clustering from a mutant library.** Experimental (9,4) peak shift, (9,4) and (7,6) intensity change response of glucose detection at 7.5 mM by ssDNA-SWCNT for different diverse ssDNA sequences, which resulted in finding more sequences responsive to glucose and improving the response, enhancing the peak shift from -2.5 to -4 nm for (9,4) and the intensity change from 40% to 70% at (7,6).

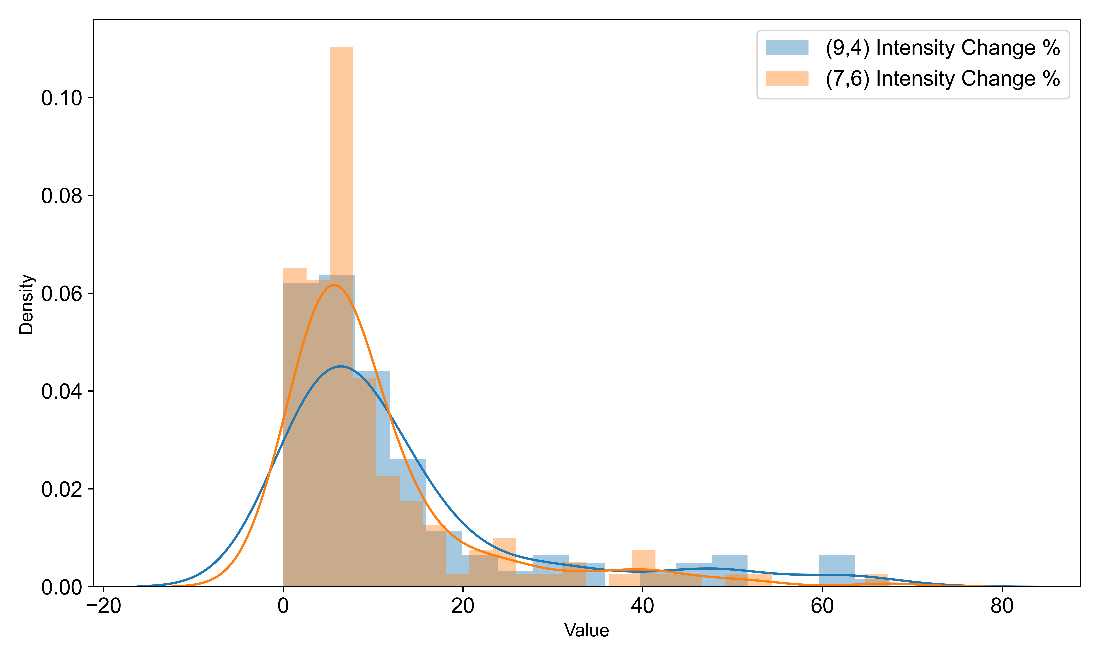

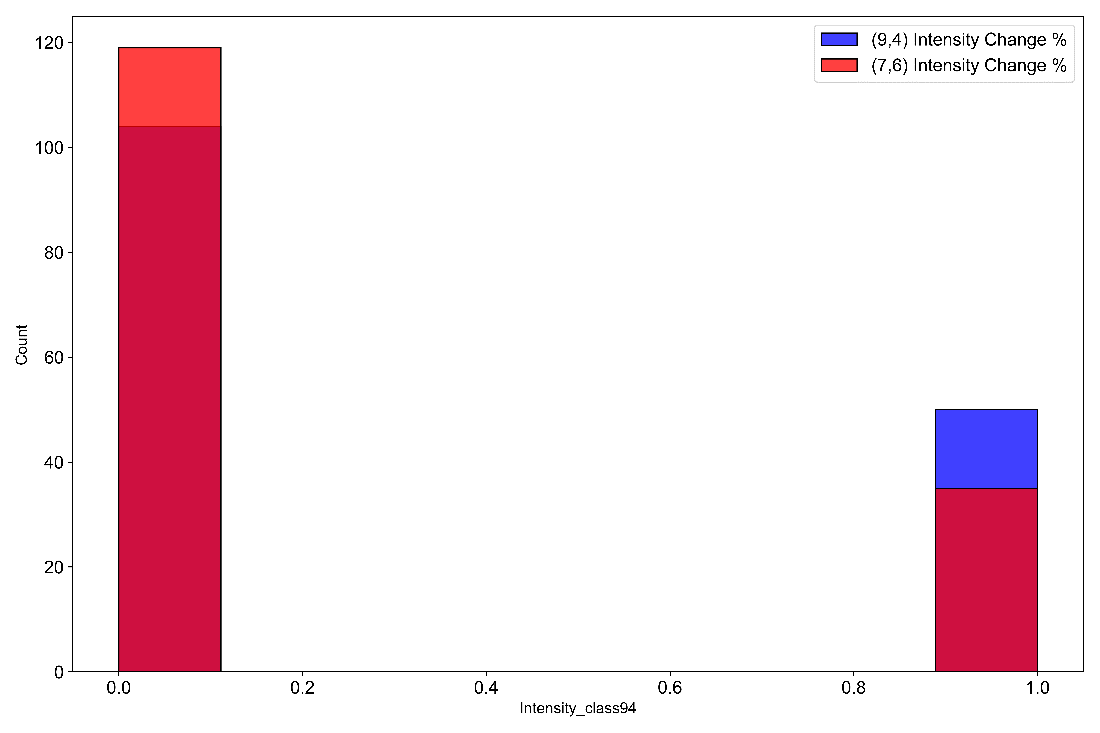

**Fig. S12: Sensor response distribution and classification**: The distribution plot for the sensor library is shown as a function of the response (top). The sensors are to be classified into two categories: class 1 (high-intensity response) and class 0 (low-intensity response) for the (9,4) and (7,6) chiralities' intensity change upon glucose addition. The classification threshold determination aims to balance the data between both classes to improve ML model training performance. However, since the majority of the data shows no response to glucose, we set a 12% threshold for intensity change as the optimal point to achieve a more balanced dataset between the two classes, ensuring better training performance (bottom).

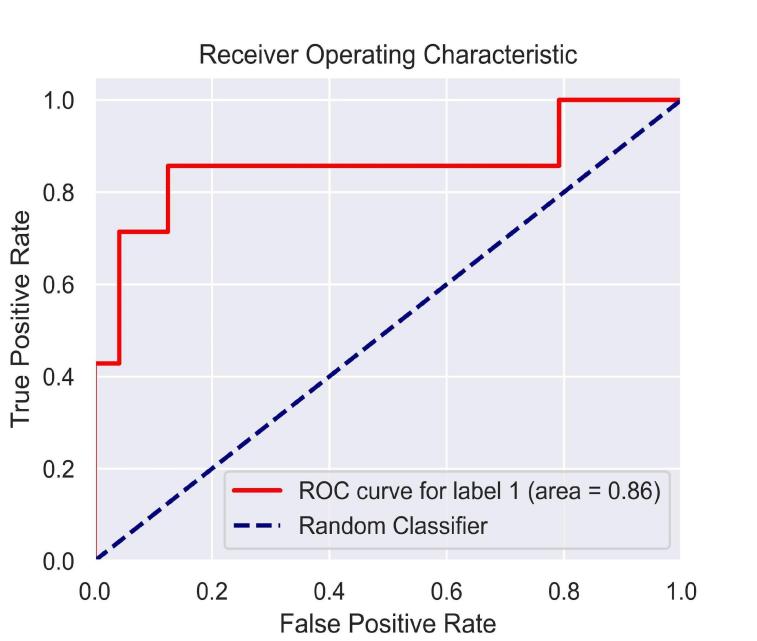

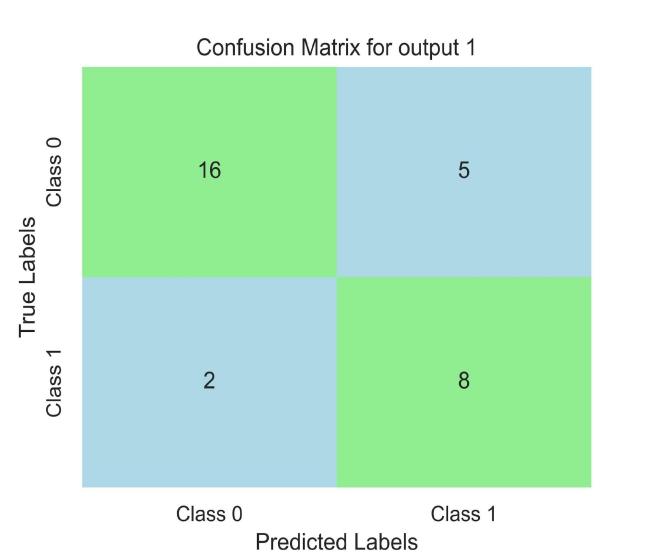

**Fig. S13- Confusion matrix and ROC curve for output 1 (Intensity change of (7,6)).** (left) Confusion matrix comparing the predicted classification from the model and the actual performance based on experimental measurements. The percentages represent the fraction of predicted Class 0 (left quadrants) or Class 1 (right quadrants) (7,6) intensity responses that were experimentally validated as either Class 0 and Class 1. Green regions represent quadrants that designate consistency (true positive and true negative) between the predicted and experimental classifications and blue regions represent quadrants that designate inconsistency (false positive and false negative) between the predicted and the experimental classifications. (right) Retention of Curve (ROC) (red) comparing the true and false positive classification rates. The blue dotted line represents equivalent true positive and false positive prediction rates for a random classifier. Results show that most of the data are predicted in the correct class, including high and low responses. The ROC curve indicates a high ratio of true positives to false positives, with an accuracy of 0.81 for output 1, as shown in Table S5.

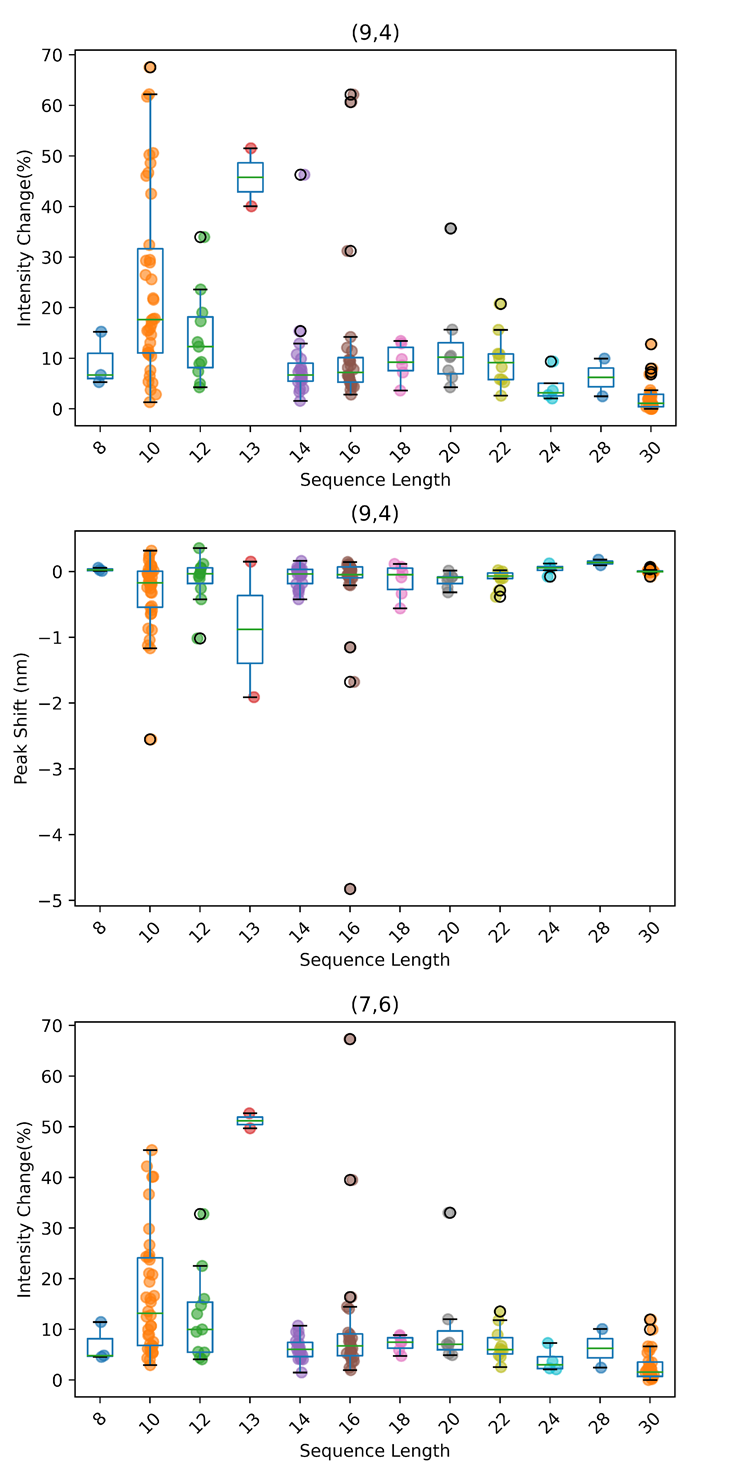

**Fig. S14** **- The effect of lengths on (9,4) intensity change, (9,4) peak shift, and (7,6) intensity change response across various DNA sequences.** The results showed that there is an optimum length range between 10 and 16 for glucose sensitivity for different DNA sequences. Additionally, the response within this range of lengths completely depends on the DNA base combination.

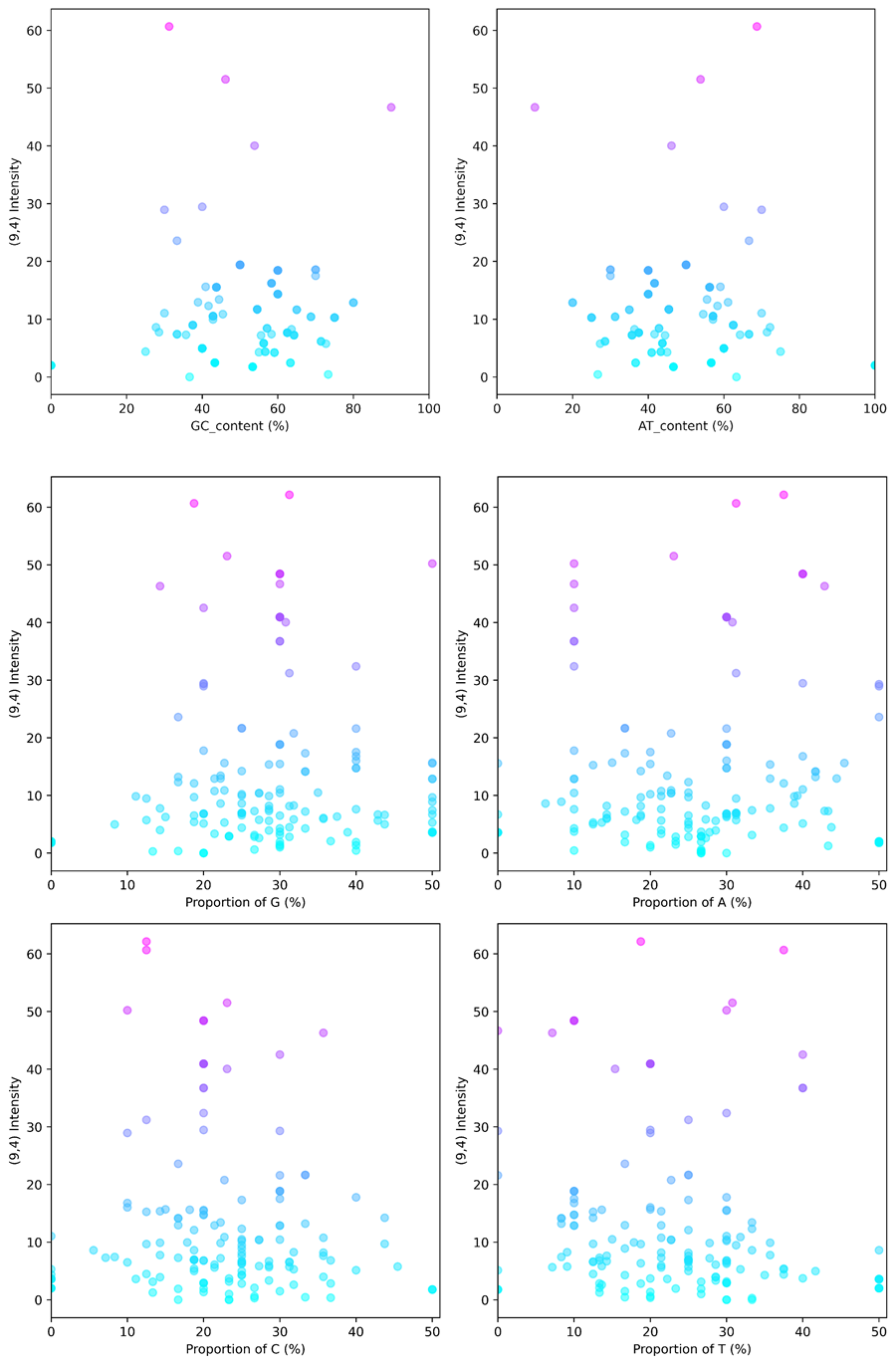

**Fig. S15: Base proportion analysis for the (9,4) intensity response.** This analysis was conducted to find the effect of each single base, such as A, G, C, and T, as well as the proportion of AT and GC content in our sequence library. The analysis showed that lower C content increases the sensitivity of the sensor for (9,4) intensity change. However, it is generally difficult to determine the effect of other bases on the response.

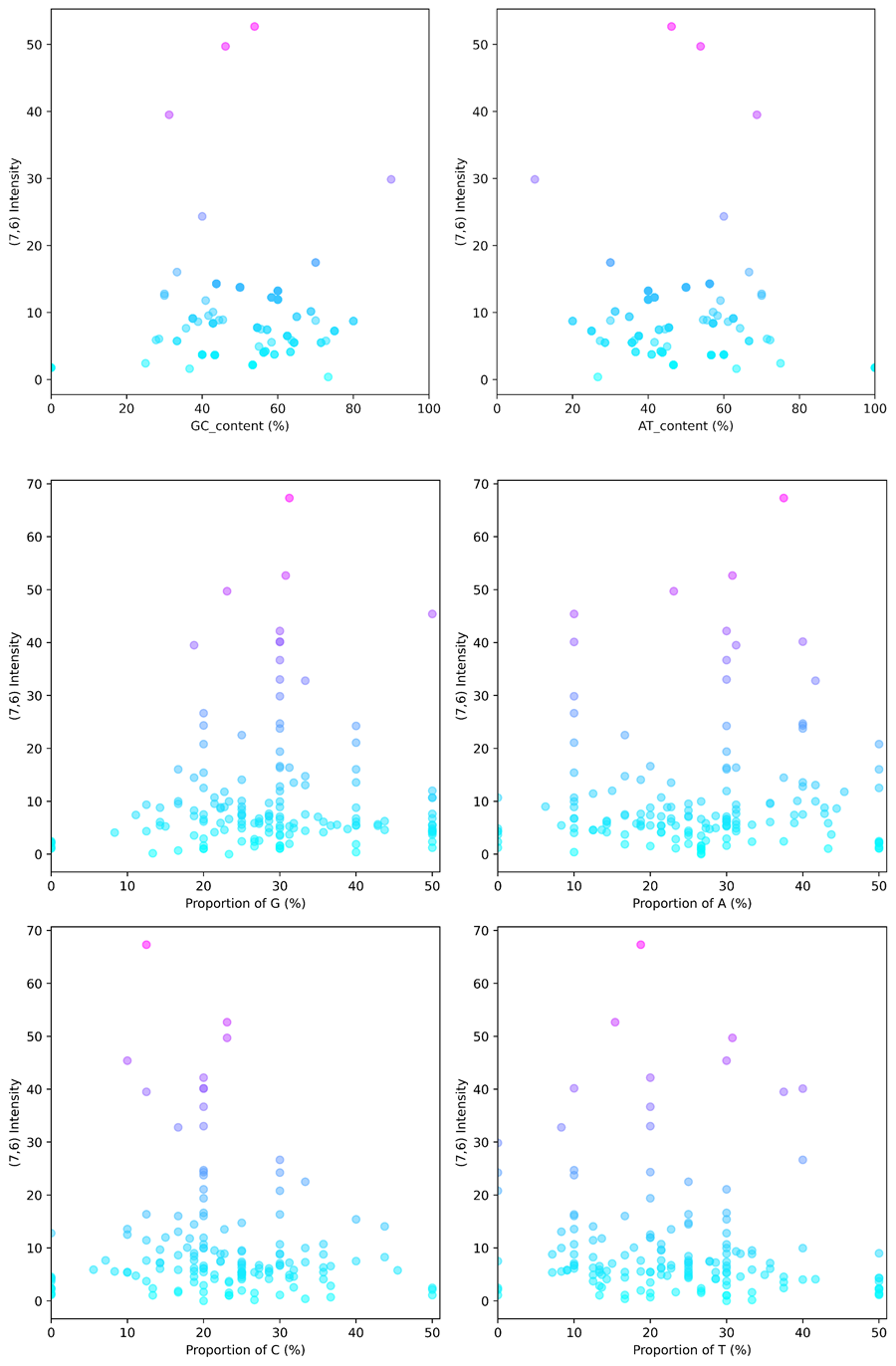

**Fig. S16: Base proportion analysis for the (7,6) intensity response.** This analysis was conducted to find the effect of each single base, such as A, G, C, and T, as well as the proportion of AT and GC content in our sequence library. The analysis showed that lower C content also increases the sensitivity of the sensor for (7,6) intensity changes. However, it is generally difficult to determine the effect of other bases on the response.

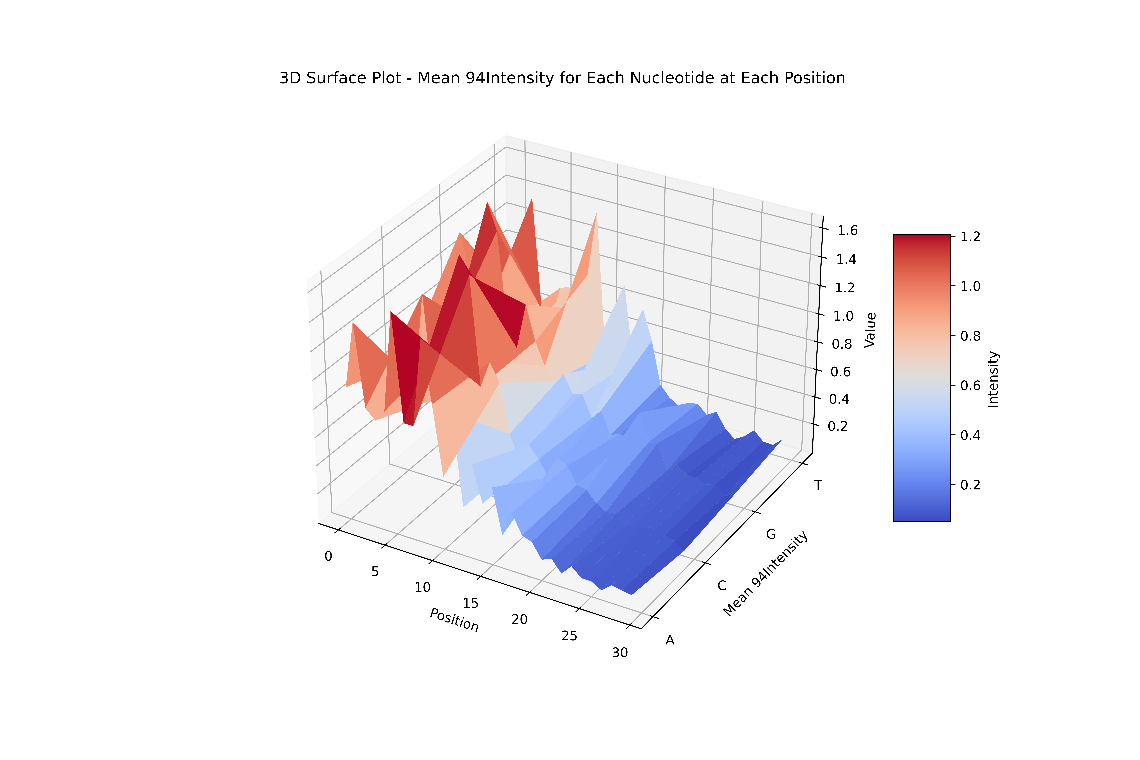

**Fig. S17: Position analysis of each nucleotide for (9,4) intensity change response.** Details of each base position's effect at different lengths of the DNA on the (9,4) chirality intensity change response are plotted. This shows that some specific bases have more effect on the response at certain specific length positions of the DNA sequence. (**An interactive plot is available on GitHub**).

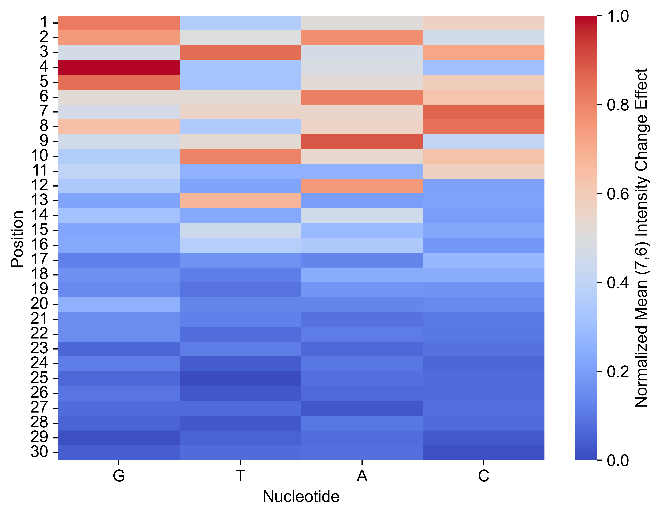

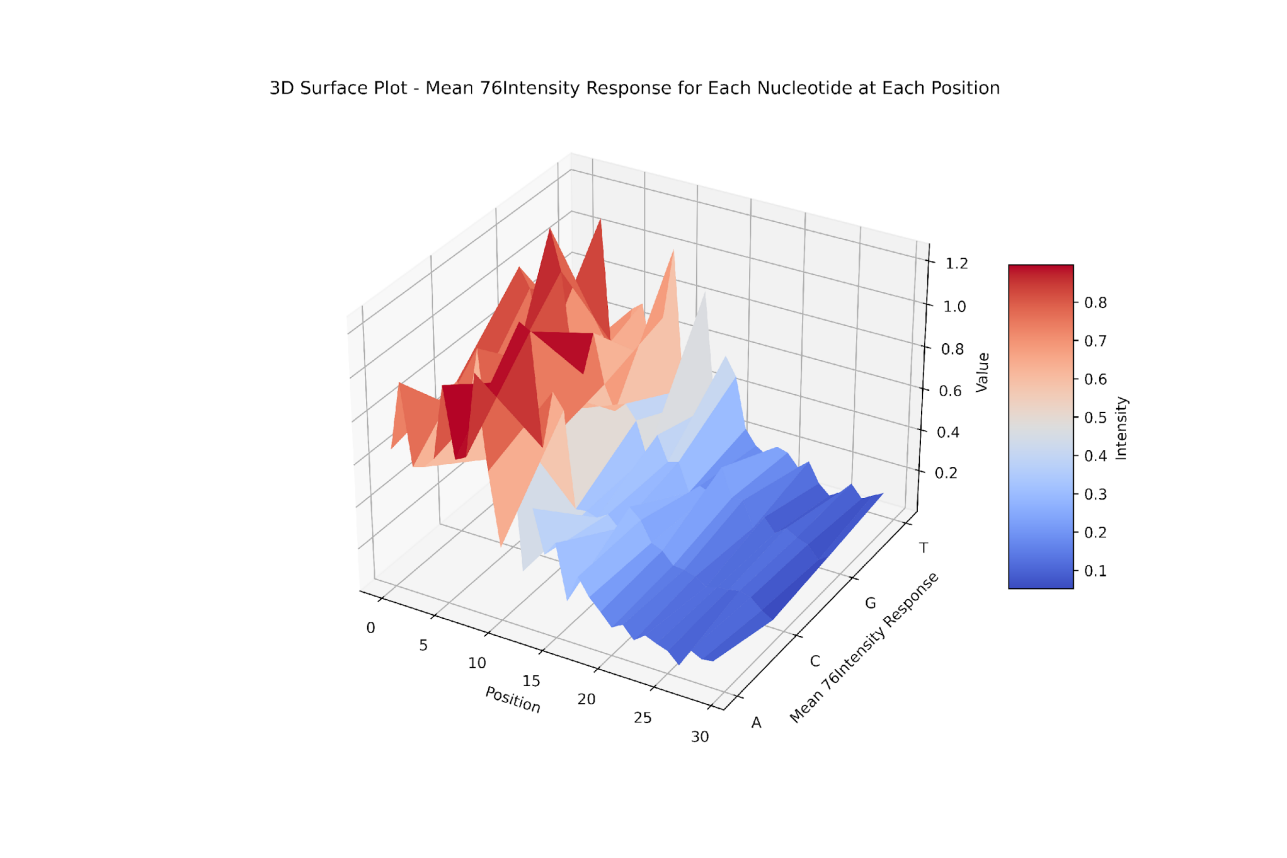

**Fig. S18: Position analysis of each nucleotide for (7,6) intensity change response.** Details of each base position's effect at different lengths of the DNA on the (7,6) chirality intensity change response are plotted. This shows that some specific bases have more effect on the response at certain specific length positions of the DNA sequence. For example, G at position 4 has a maximum effect on the (7,6) intensity change response, and C at position 30 has a minimum effect, as shown on the heatmap. (**An interactive plot is available on GitHub**).

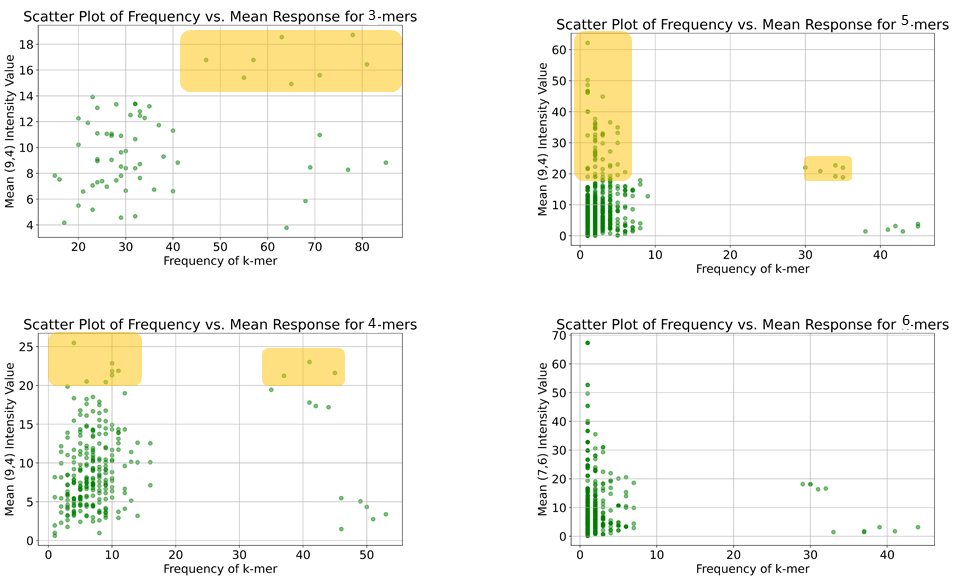

**Fig. S19: Analysis of different K-mers (K=3 to 6) to define a suitable K-mers length.** This analysis shows that a 3-mers have a good distribution and frequency, also among sequences with high fluorescence responses, making it a suitable choice for selection.

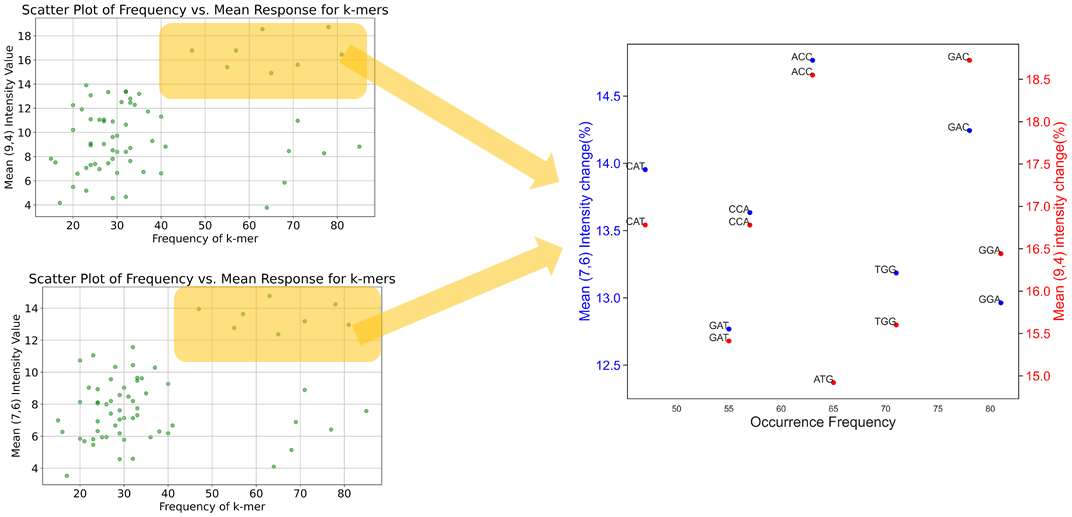

**Fig. S20: K-mers analysis for the (9,4) and (7,6) intensity change response.** First, the entire combination of K-mers in our sequence library was extracted, and the mean effect of each K-mers was calculated for the whole sequence. Based on the frequency of the K-mers with the response (left plot), the K-mers with a high response effect were selected. This selection included frequent K-mers that showed a high response at both the (7,6) and (9,4) chirality intensity change responses, as shown on the right side of the figure with several 3-mer bases.

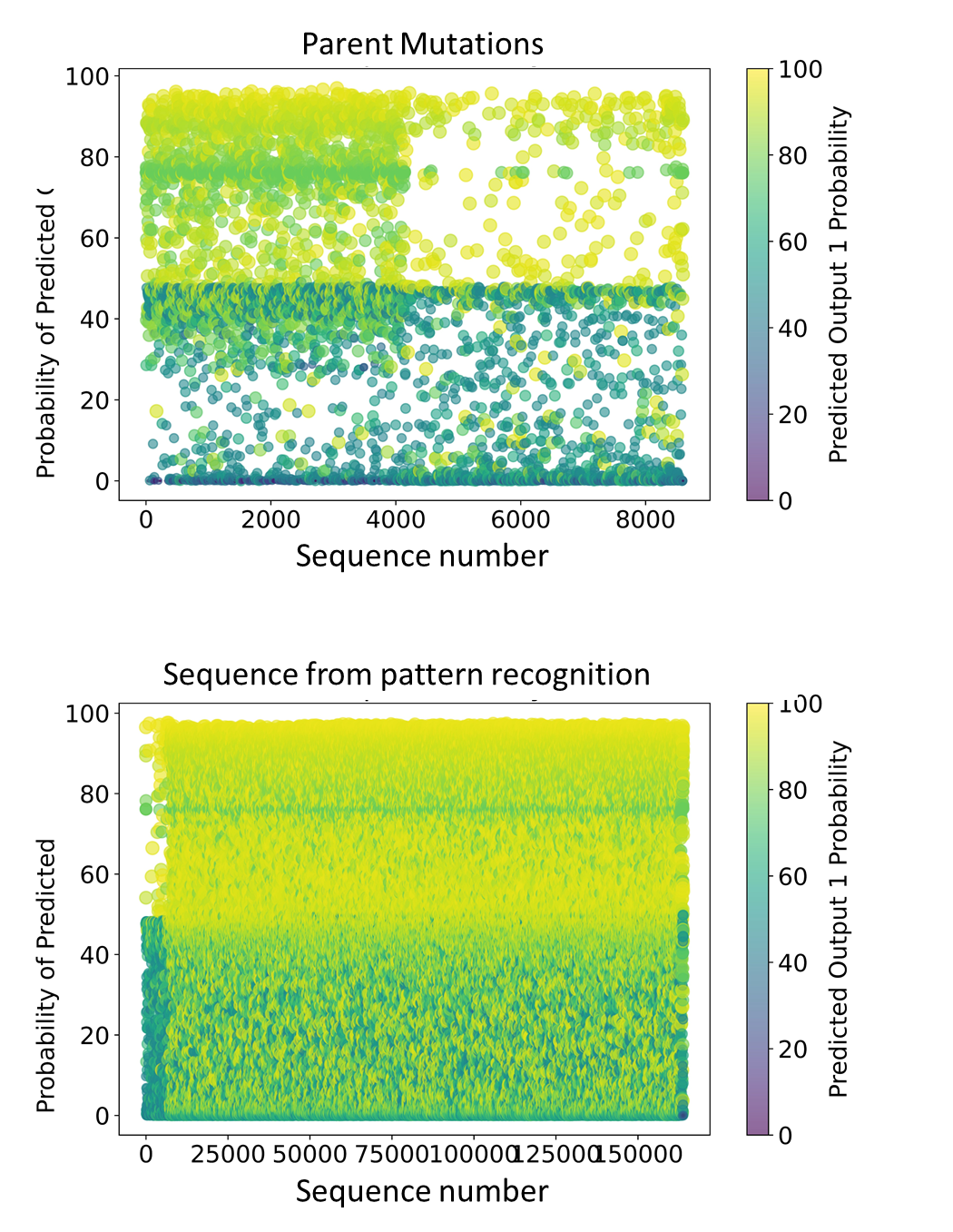

**Fig. S21: Deep neural network Prediction for the Library of Parent Mutations and Pattern Recognition Analysis.** The entire library of DNA sequences generated from pattern recognition, which is around 160k sequences, was predicted by our optimal deep neural network (bottom). We also applied the network to a previous library (size = 8000) from the clustering and mutation steps (top). The probability of each sequence being in class 1 (high intensity change response) is shown in this figure. The DNA sequences with the highest probabilities were selected for the next step of experimental screening.

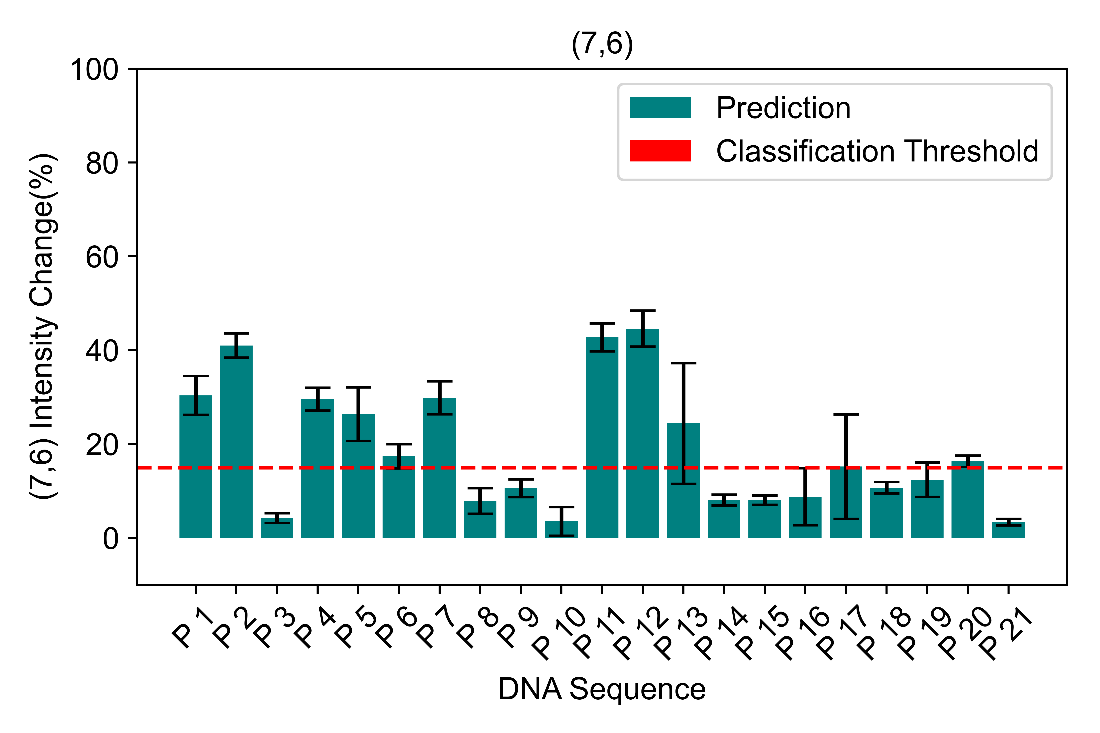

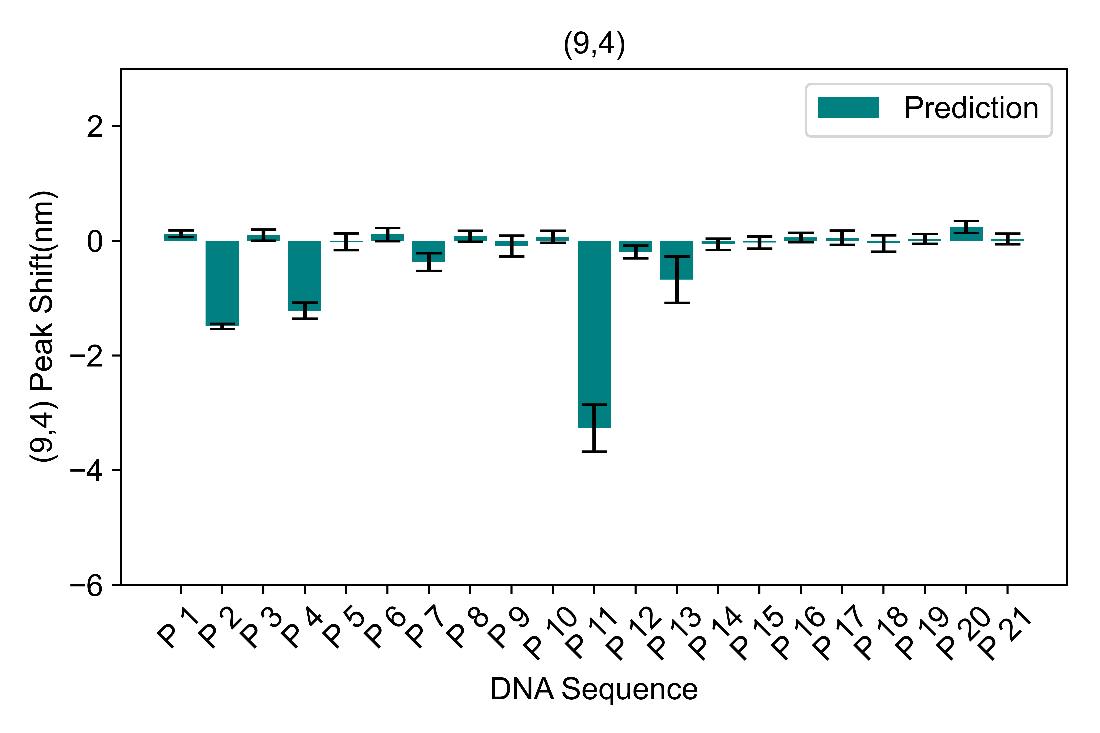

**Fig. S22 - Prediction by Deep Neural Network.** Experimental validation of the (7,6) intensity and (9,4) peak shift response of DNA sequences predicted by ML with a high probability of having a strong response. This led to the discovery of more responsive sequences for detecting glucose.

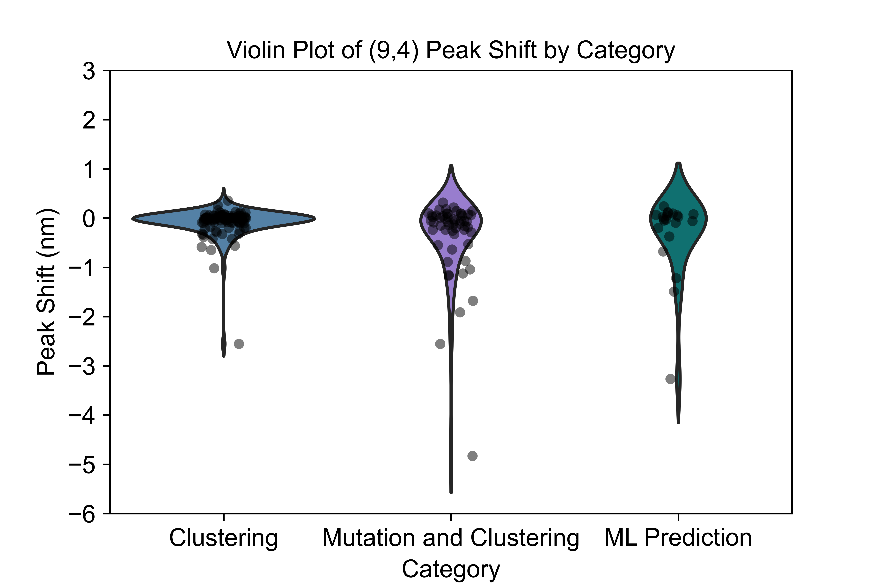

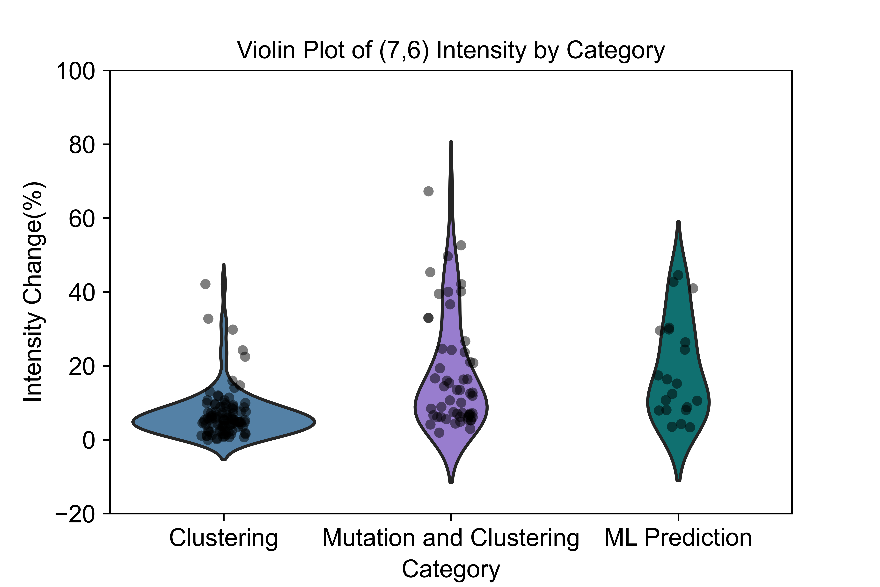

**Fig. S23 – Comparison of the experimental glucose sensing results across the main computational steps used in this study.** Comparison of the different steps, including clustering, mutation and clustering, and ML training, of glucose distribution response at the peak shift of (9,4) chirality and intensity change of (7,6) for different DNA sequences. This result revealed that ML training has a strong ability to discover new sequences using a small library size.

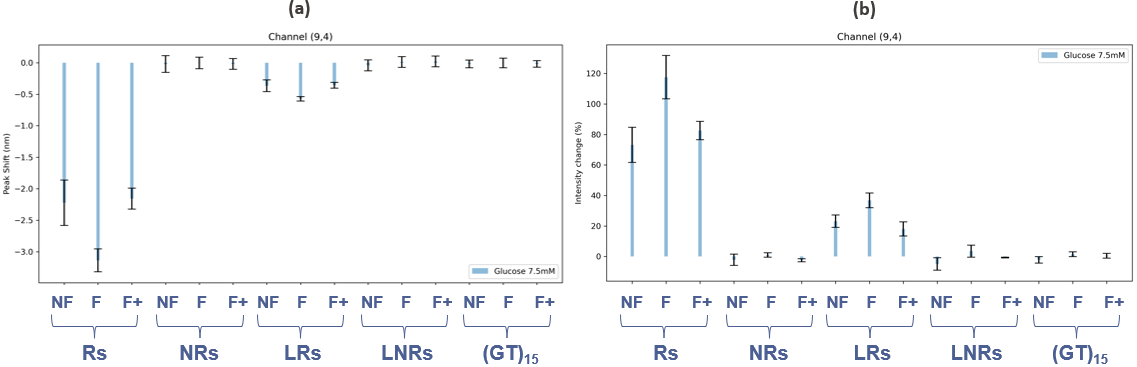

**Fig. S22- Influence of free DNA on the sensitivity of ssDNA-SWCNTs to glucose** (7.5 mM) for the chirality (9,4). Peak shifts (a) and intensity changes (b) induced by five different sequences are presented: Rs = responsive sequences, NRs = non-responsive sequences of the same length, LRs = longer responsive sequences, LNRs = non-responsive sequence of the same longer length, and (GT)15 as negative control. For each sensor, three different conditions are presented: NF = non-filtered ssDNA-SWCNTs, F = filtered ssDNA-SWCNTs, and F+ = filtered ssDNA-SWCNTs with post-addition of free DNA.

| Responsive sequence | Non-Responsive sequence |
| --- | --- |
| 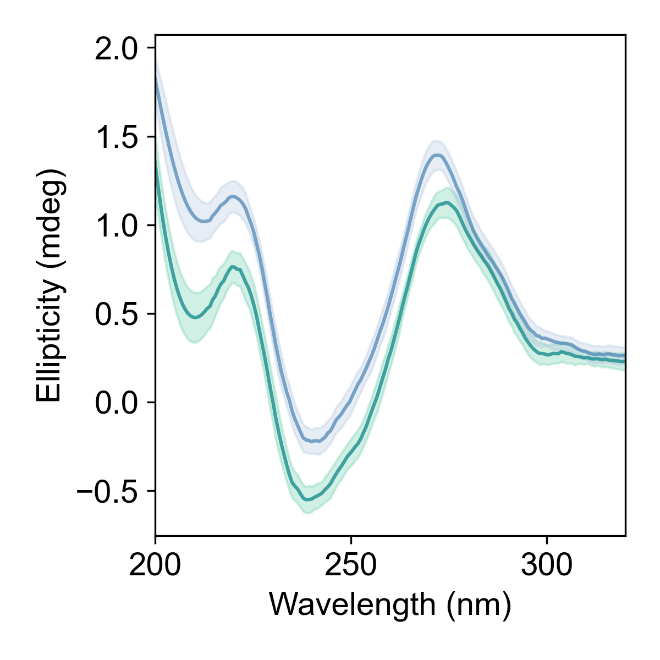 | 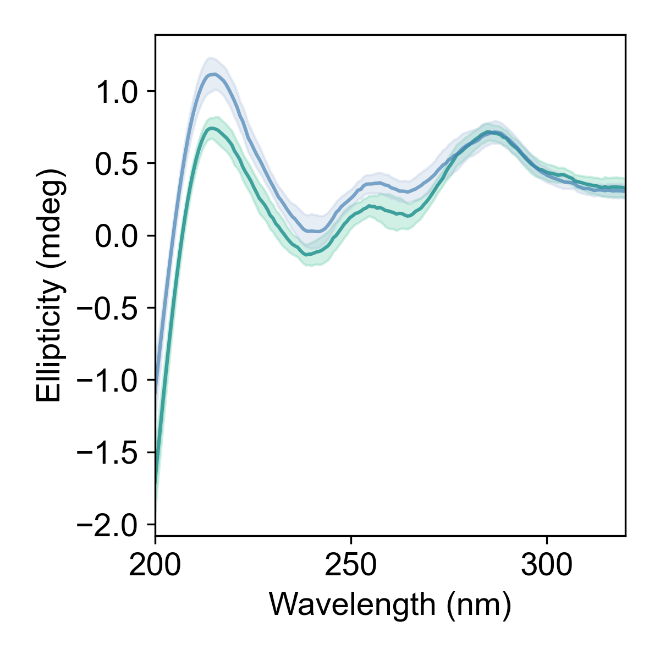 |
| GT15 | |
| 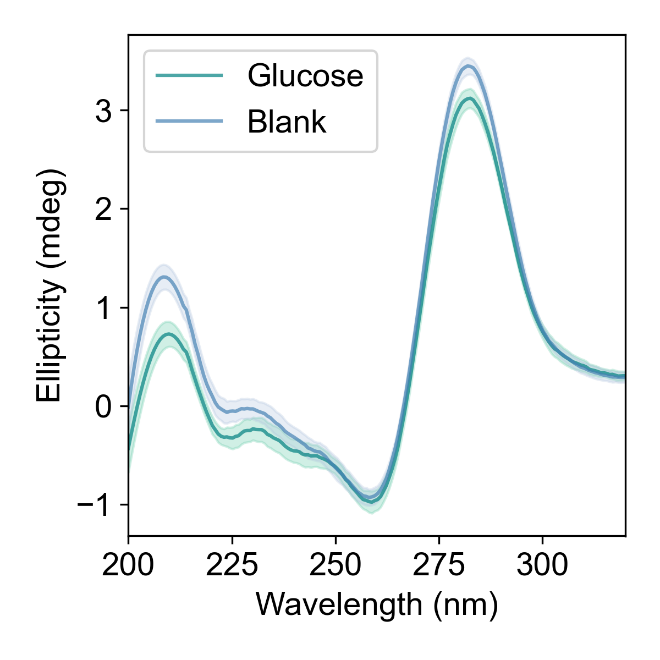 | |

**Fig. S23 - CD measurement results for DNA samples. S**tructural changes were studied for responsive and non-responsive sequences, with GT15 as a negative control, with and without the addition of 40 mM glucose. This shows peak shift changes due to the conformation change of our responsive sequence.

| Responsive sequence | Non-Responsive sequence |
| --- | --- |
| 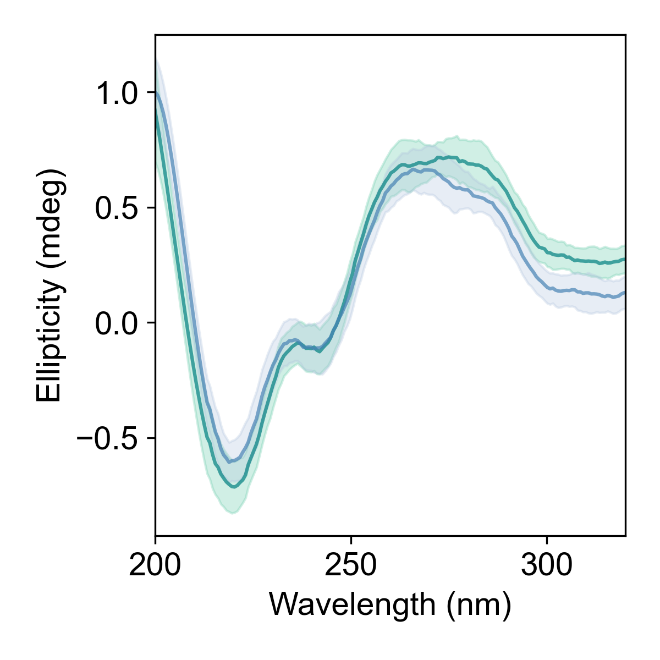 | 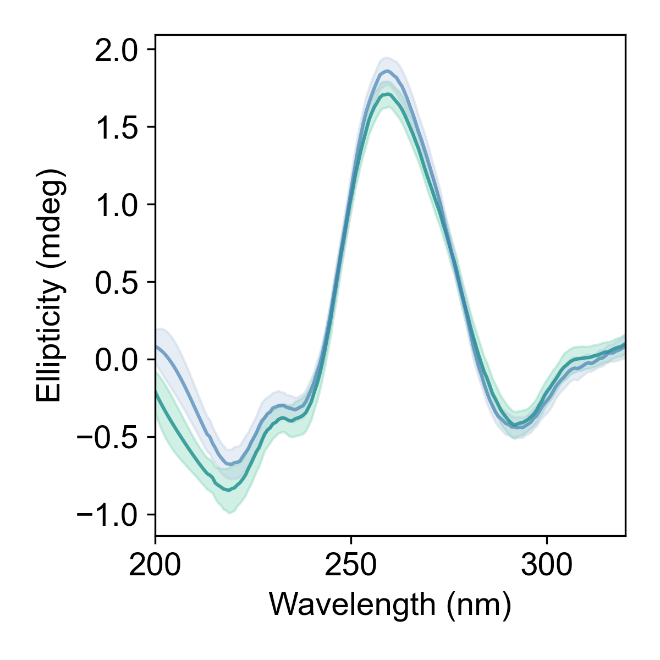 |
| GT15 | |
| 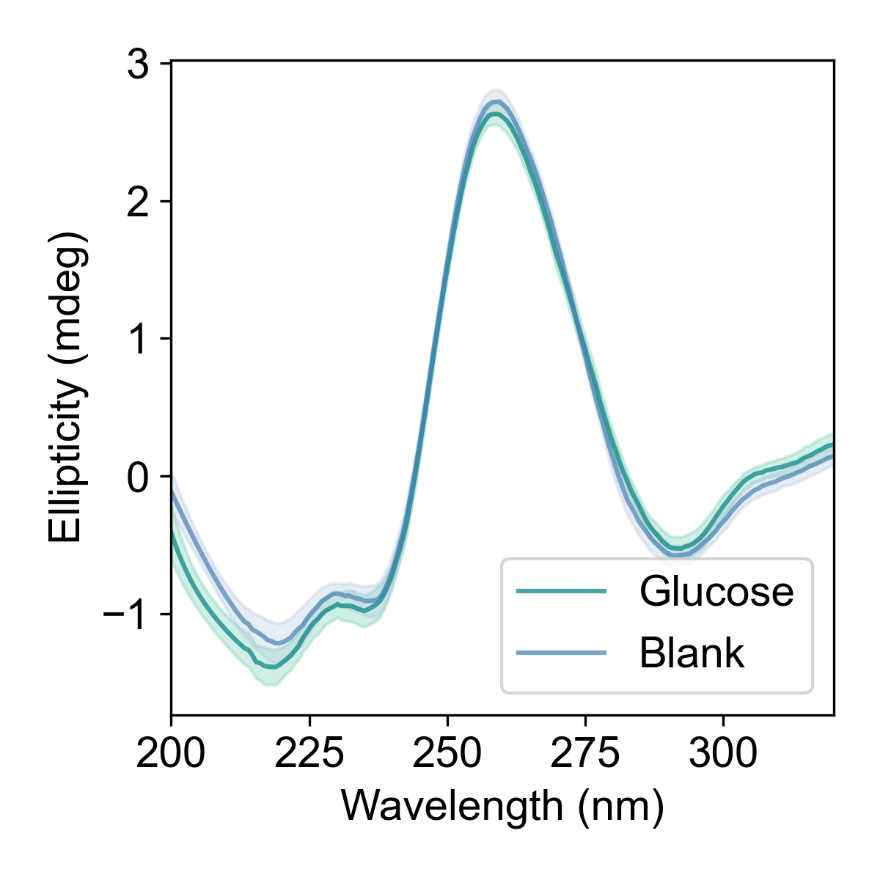 | |

**Fig. S24 - CD measurement results for DNA-SWCNT samples. S**tructural changes were studied for responsive and non-responsive sequences, with GT as a negative control, with and without the addition of 40 mM glucose. This shows the conformation change of DNA on the surface of the CNT upon the addition of glucose.

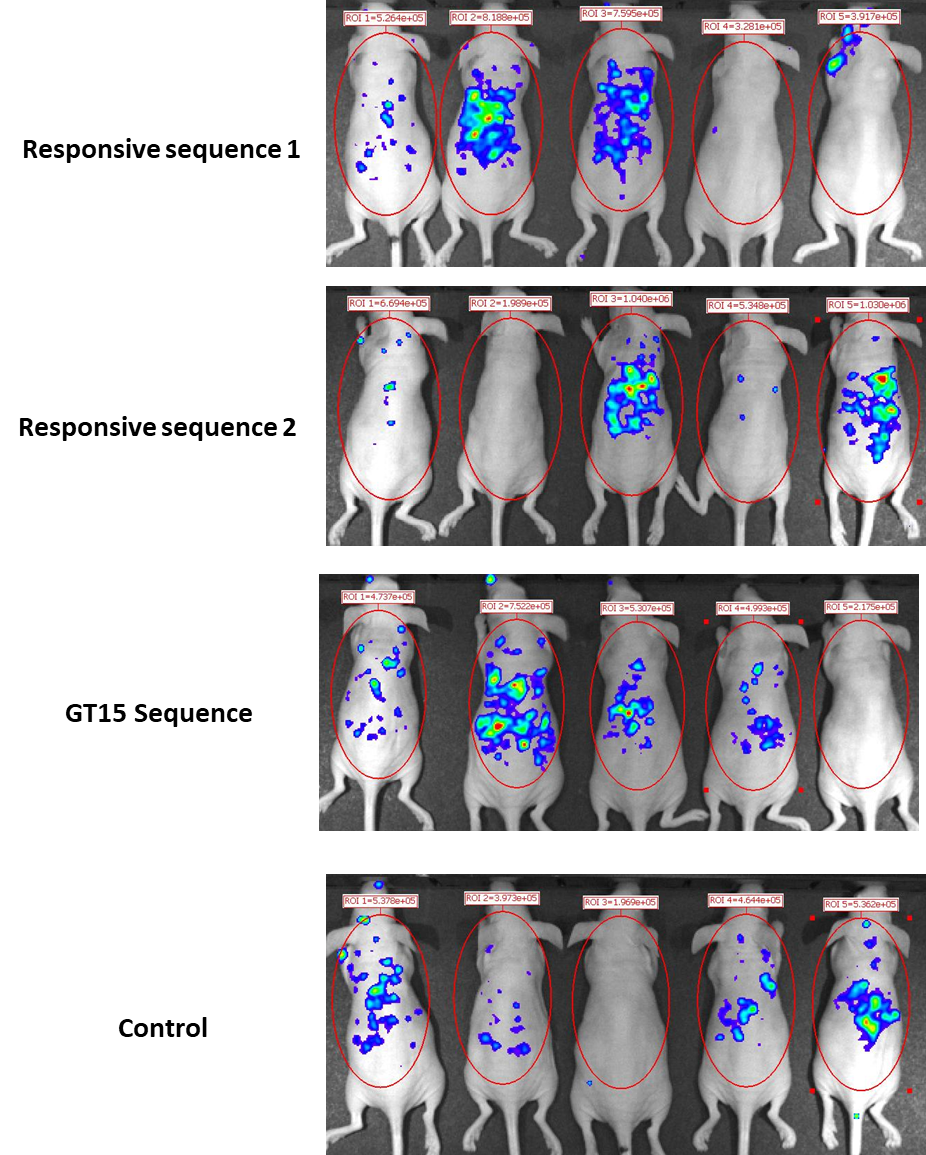

**Figure S25-** **Evaluation of the ROS production in mice injected ssDNA-SWCNT samples** for sequence responsive 1, sequence responsive 2, GT15 and in a control group for 5 animals per group. Mice were injected intraperitoneally with 200mg of luminol and imaged 15 minutes post-injection. Luminol was previously used in multiple publications to detect ROS production with high sensitivity.

**Table S1. DNA folding-based (Folding) DNA clustering:** Clustering results based on folding properties for random DNA sequences as an example. In this method, the folding prediction of each single strand was performed using ViennaRNA folding prediction. The folding properties were then extracted using a Python script, and combined with data on the physicochemical properties of the DNA for k-means clustering based on the different properties.

| **DNA** | **Avg_Atomic_Number** | **Avg_EIIP** | **Avg_Molecular_Weight** | **Melting_Temp** | **Seq_Length** | **Seq_Entropy** | **Seq_MFE** | **Ensemble_Energy** | **Centroid_Structure** | **Paired_Probs** | **Cluster** |
| --- | --- | --- | --- | --- | --- | --- | --- | --- | --- | --- | --- |
| GTAAAATTTGAGATTG | 70.5 | 0.117463 | 331.825 | 64.25938 | 16 | 1.561278 | 0 | -0.04447 | ................ | 0.218205 | 1 |
| TATAGGAGCCTTTACTGC | 67.77777778 | 0.120189 | 326.4222 | 64.77346 | 18 | 1.974938 | 0 | -0.49682 | .................. | 1.296238 | 2 |
| TAGTGGTAAGCCTATTACCGTC | 67.81818182 | 0.119886 | 326.5182 | 64.81529 | 22 | 1.98305 | -4.7 | -5.26318 | ...((((((.....)))))).. | 5.584762 | 2 |
| CCTTTTGTGGAGACGCAG | 68.88888889 | 0.114728 | 328.7 | 65.02654 | 18 | 1.954686 | -3.4 | -3.81254 | .....((((....)))). | 3.866314 | 3 |
| GCGATGACTGGGTAGGTGGC | 71.4 | 0.106 | 333.8 | 65.2075 | 20 | 1.785475 | -0.8 | -1.56957 | ((.((..........)).)) | 3.276788 | 4 |
| TTATATCGCAATAG | 68 | 0.123336 | 326.8429 | 64.27245 | 14 | 1.863121 | 0 | -0.05786 | .............. | 0.223704 | 5 |
| TCTGTATTTCTTCGGG | 67.75 | 0.1199 | 326.2 | 64.73984 | 16 | 1.70282 | -1.1 | -1.43587 | .(((........))). | 2.724948 | 1 |
| GACGCCCGCC | 65.2 | 0.11718 | 321.6 | 66.54 | 10 | 1.295462 | 0 | -0.02486 | .......... | 0.039986 | 4 |
| GATCGTCAAAAACCGCGGTTTGACGGGTAT | 69.06666667 | 0.11573 | 329.1 | 64.9 | 30 | 1.983871 | -7.3 | -8.13996 | ..((((((((........)))))))).... | 7.935374 | 3 |
| GGTAAAGTCCAAAGATGCATGAGGGACAAA | 70.26666667 | 0.114447 | 331.6 | 64.80889 | 30 | 1.819056 | -4 | -4.32696 | ......((((.............))))... | 4.369561 | 3 |
| CTGGGTCACACCCTGAGTGAAGGCCAATCG | 68.13333333 | 0.11603 | 327.3 | 65.03667 | 30 | 1.962898 | -5.8 | -6.35783 | ........(((.....)))........... | 6.746134 | 2 |
| GTGTGTGTGTGTGTGTGTGTGTGTGTGTGT | 72 | 0.10705 | 334.7 | 64.9 | 30 | 1 | 0 | -0.01759 | .............................. | 0.040188 | 1 |
| ATATATATATATATATATATATATATATAT | 68 | 0.12975 | 326.7 | 64.21667 | 30 | 1 | -8.4 | -9.12417 | .(((((((((((......))))))))))). | 11.48879 | 3 |
| GATGGAACCT | 69.2 | 0.11548 | 329.4 | 64.9 | 10 | 1.970951 | 0 | -0.01363 | .......... | 0.021362 | 2 |
| AATGGACCGC | 68.4 | 0.11553 | 327.9 | 65.31 | 10 | 1.895462 | 0 | -0.00505 | .......... | 0.005496 | 1 |
| GGATGGACCATC | 69 | 0.114117 | 329.0333 | 65.18472 | 12 | 1.959148 | -0.4 | -0.92899 | ............ | 1.863047 | 6 |
| GATGGACCATT | 68.90909 | 0.117118 | 328.7455 | 64.73058 | 11 | 1.980826 | 0 | -0.10618 | ........... | 0.405316 | 3 |
| TGATGGACCATT | 68.66667 | 0.118483 | 328.2 | 64.61528 | 12 | 1.959148 | 0 | -0.15567 | ............ | 0.563051 | 2 |
| GGATGGACCATT | 69.66667 | 0.114075 | 330.2833 | 64.9 | 12 | 1.959148 | -0.4 | -0.72338 | ((.....))... | 1.432835 | 1 |
| TGATGGACCATA | 69 | 0.117858 | 328.95 | 64.61528 | 12 | 1.959148 | 0 | -0.10062 | ............ | 0.339013 | 1 |

**Table S2. Experimental results of the sensor from diverse sequences selected through clustering method.** Experimental res(9,4) peak shift, (9,4) and (7,6) intensity changes in response to glucose detection at 7.5 mM using ssDNA-SWCNT for different diverse ssDNA sequences, including clustering information, along with the DNA sequence.

| Num. | Method | **DNA Sequence** | **(7,6) Intensity** | **(7,6) Intensity_Std** | **(9,4) Shift** | **(9,4) Shift_Std** | **(9,4) Intensity** | **(9,4) Intensity_Std** |
| --- | --- | --- | --- | --- | --- | --- | --- | --- |
| 1 | MAFFT | AGTAAGCTCCACGCGCCGGAAG | 5.84 | 0.87 | -0.04 | 0.06 | 8.25 | 0.98 |
| 2 | MAFFT | ACCAGACCTATTATAATG | 7.41 | 0.59 | -0.01 | 0.05 | 9.84 | 1.06 |
| 3 | MAFFT | GTTGACGCGGTTAC | 4.15 | 0.63 | -0.04 | 0.07 | 6.09 | 0.47 |
| 4 | MAFFT | GAGGCTCCCAGCAT | 4.10 | 0.83 | -0.29 | 0.07 | 7.63 | 0.73 |
| 5 | MAFFT | AGGTCCTGTGTCGGGA | 4.59 | 0.64 | 0.07 | 0.05 | 4.99 | 0.46 |
| 6 | MAFFT | TGTACTAGGAAGCA | 9.67 | 2.92 | -0.42 | 0.13 | 15.36 | 5.11 |
| 7 | MAFFT | AAAACCGGGGAT | 13.06 | 7.89 | -0.26 | 0.11 | 19.02 | 8.91 |
| 8 | MAFFT | TACGATAACTCGTGGA | 7.66 | 0.35 | -0.09 | 0.03 | 11.41 | 1.55 |
| 9 | MAFFT | TCACGTTAGCCAAACT | 4.34 | 0.75 | 0.00 | 0.04 | 5.72 | 1.17 |
| 10 | MAFFT | TCTCGACTGGAGTTCTTGGGTCGCCATAAG | 3.72 | 1.89 | 0.01 | 0.07 | 3.17 | 1.00 |
| 11 | MAFFT | CGGAACTAATAAAGAGCTCAAG | 11.79 | 1.86 | -0.38 | 0.12 | 15.61 | 2.12 |
| 12 | MAFFT | TCTCAGCGACAATTATAGCT | 5.25 | 1.43 | -0.09 | 0.08 | 6.25 | 1.53 |
| 13 | MAFFT | GAGAGCCCTGCTGT | 6.12 | 1.32 | 0.02 | 0.08 | 5.93 | 0.91 |
| 14 | MAFFT | GAGAGCCCTGCTGG | 5.65 | 2.07 | -0.07 | 0.06 | 6.68 | 1.55 |
| 15 | MAFFT | GATCGTCAAAAACCGCGGTTTGACGGGTAT | 0.83 | 0.13 | -0.01 | 0.01 | 1.22 | 0.09 |
| 16 | MAFFT | GGTAAAGTCCAAAGATGCATGAGGGACAAA | 0.82 | 0.17 | -0.06 | 0.01 | 1.53 | 0.15 |
| 17 | MAFFT | GGGTATCGTGTCGGGGGTACAGACGTTATC | 0.92 | 0.13 | 0.01 | 0.01 | 1.00 | 0.09 |
| 18 | MAFFT | GAGTTAGGTTCTCTGCTTCTACACAGGAAA | 6.63 | 0.00 | 0.00 | 0.00 | 0.78 | 0.00 |
| 19 | MAFFT | TAGTTAGGTTCTATGCTTCTACACAGGAAA | 1.61 | 0.00 | 0.00 | 0.00 | 0.00 | 0.62 |
| 20 | Folding | ACGAGGAGCTTGGTGG | 7.62 | 0.41 | -0.09 | 0.07 | 9.66 | 1.47 |
| 21 | Folding | AGACTTGTCGATTTAC | 3.54 | 0.40 | 0.05 | 0.03 | 5.39 | 0.29 |
| 22 | Folding | GGGATAGACAAGACAT | 3.72 | 3.84 | -0.05 | 0.07 | 4.47 | 4.27 |
| 23 | Folding | CTGGCGCGCGAAGA | 5.36 | 0.44 | 0.01 | 0.02 | 5.64 | 0.67 |
| 24 | Folding | TCGAGTCACTGGAA | 4.94 | 1.01 | 0.04 | 0.03 | 5.59 | 0.82 |
| 25 | Folding | TAGTGAGGTTCTCTGCTTCTACACAAGCAA | 1.03 | 2.00 | 0.00 | 0.00 | 0.00 | 0.00 |
| 26 | Folding | CGTGGCGGTGCATCAGACCCCGTGTGCCGG | 0.38 | 1.26 | 0.00 | 0.00 | 0.44 | 1.62 |
| 27 | Folding | AGACCACCCAGTTGACGATACGCCCCTGAA | 6.56 | 4.24 | 0.00 | 0.00 | 6.83 | 4.33 |
| 28 | Folding | GCGCGGGCCGAAGTAATGTCAGGGCACCTA | 5.40 | 2.51 | 0.00 | 0.00 | 2.07 | 2.60 |
| 29 | Folding | TAAACTCAACCCCCCTGGAACTCAGTGGTC | 0.69 | 1.75 | 0.00 | 0.00 | 0.35 | 2.03 |
| 30 | Folding | GTAAAATTTGAGATTG | 2.42 | 1.06 | -0.10 | 0.28 | 4.39 | 1.79 |
| 31 | Folding | TATAGGAGCCTTTACTGC | 8.86 | 1.45 | -0.09 | 0.22 | 13.41 | 2.42 |
| 32 | Folding | TAGTGGTAAGCCTATTACCGTC | 8.91 | 1.54 | 0.02 | 0.22 | 10.87 | 2.13 |
| 33 | Folding | AGGACCAGGC | 24.24 | 2.54 | 0.26 | 0.13 | 21.59 | 0.90 |
| 34 | Folding | TCCCTTAGATAG | 9.53 | 2.06 | 0.12 | 0.15 | 12.30 | 2.04 |
| 35 | Folding | CCTTTTGTGGAGACGCAG | 7.52 | 1.07 | 0.12 | 0.20 | 7.20 | 1.99 |
| 36 | Folding | GCGATGACTGGGTAGGTGGC | 11.99 | 1.69 | -0.08 | 0.25 | 15.68 | 3.43 |
| 37 | Folding | TTATATCGCAATAG | 6.07 | 1.86 | 0.07 | 0.22 | 7.75 | 1.46 |
| 38 | Folding | TCTGTATTTCTTCGGG | 8.99 | 0.91 | 0.12 | 0.18 | 8.59 | 1.59 |
| 39 | Folding | GACGCCCGCC | 29.85 | 4.05 | -0.59 | 0.35 | 46.67 | 11.45 |
| 40 | Folding | GTGAGGCG | 11.45 | 2.56 | 0.01 | 0.18 | 15.25 | 1.86 |
| 41 | Folding | CAGTATGAAATGGA | 7.65 | 0.81 | -0.03 | 0.16 | 7.30 | 1.52 |
| 42 | Folding | TGCTGTCTGCGAAGGTCCGATTCCTAAGAG | 1.53 | 1.85 | 0.00 | 0.00 | 0.99 | 1.37 |
| 43 | Folding | GGGAATATCCAGACACCGCTGTGAGACTTG | 3.49 | 0.99 | 0.00 | 0.00 | 3.69 | 0.83 |
| 44 | Folding | AACTAGTTCGAGATGAGGTACGTACTTGAA | 1.78 | 0.31 | 0.00 | 0.04 | 2.17 | 0.43 |
| 45 | Folding | CTGGGTCACACCCTGAGTGAAGGCCAATCG | 4.85 | 0.51 | 0.05 | 0.03 | 4.32 | 0.18 |
| 46 | Folding | TAGTGAGGTTCTCTGCTTCTACACAGGAAA | 0.00 | 0.00 | 0.00 | 0.00 | 0.00 | 0.00 |
| 47 | Folding | GCGAAACAACTGCGAACGGTTGGAAATAAA | 5.86 | 0.90 | 0.00 | 0.00 | 7.28 | 1.18 |
| 48 | Folding | CCCGTAGGAATTGGAGAGTTCACGCACATC | 1.48 | 2.42 | 0.00 | 0.00 | 0.61 | 2.27 |
| 49 | K-mers | CGGGGCAGTC | 10.69 | 3.74 | -0.19 | 0.10 | 14.65 | 4.58 |
| 50 | K-mers | TTCTCGACACGGCG | 6.31 | 2.11 | -0.04 | 0.07 | 8.18 | 2.48 |
| 51 | K-mers | TGGTATGGTG | 4.03 | 0.74 | 0.08 | 0.07 | 3.73 | 0.97 |
| 52 | K-mers | GATACCTATAGGTG | 7.24 | 1.64 | 0.16 | 0.05 | 3.92 | 1.78 |
| 53 | K-mers | ATACGTCTTTAC | 4.08 | 2.43 | -0.03 | 0.04 | 4.97 | 2.18 |
| 54 | K-mers | GGAGTTTG | 4.55 | 0.80 | 0.06 | 0.05 | 5.29 | 0.91 |
| 55 | K-mers | ATGCAGCGGT | 5.44 | 0.85 | -0.20 | 0.38 | 1.34 | 17.28 |
| 56 | K-mers | GCTGCTACACCTAG | 10.72 | 2.30 | -0.32 | 0.05 | 10.76 | 1.49 |
| 57 | K-mers | ACGCTGGCGTGG | 5.44 | 0.87 | -0.07 | 0.06 | 8.90 | 1.12 |
| 58 | K-mers | GCTCCGAAGTACCCGC | 14.07 | 0.90 | -0.20 | 0.04 | 14.22 | 1.06 |
| 59 | K-mers | CTAGTTGCTCGTATGGCGCT | 4.92 | 1.02 | 0.01 | 0.05 | 4.28 | 0.66 |
| 60 | K-mers | ACGGTAATCCGGAGCAACGCTG | 4.93 | 2.12 | -0.10 | 0.07 | 5.84 | 1.55 |
| 61 | K-mers | GTACTGGCGGGCTATTGGCC | 6.75 | 1.78 | -0.08 | 0.03 | 7.60 | 1.50 |
| 62 | K-mers | GTCGGGGATTTGGAAACA | 4.75 | 0.27 | 0.07 | 0.04 | 3.60 | 0.65 |
| 63 | K-mers | GTTACTAGTCAGTG | 7.14 | 1.82 | -0.18 | 0.04 | 9.89 | 1.11 |
| 64 | K-mers | GACAAAGGATAGATTTTT | 5.90 | 0.57 | -0.33 | 0.05 | 8.59 | 1.06 |
| 65 | K-mers | TTAATGGGCTCCGTACTGCA | 7.38 | 0.78 | -0.32 | 0.03 | 10.21 | 1.24 |
| 66 | K-mers | CCGCTAAACGAA | 9.99 | 2.26 | 0.06 | 0.04 | 13.20 | 1.79 |
| 67 | K-mers | CCCCTTGTGAGTGTCCAGCTAT | 4.59 | 1.59 | -0.08 | 0.04 | 5.29 | 1.46 |
| 68 | K-mers | GCACGACCCATACAGGTGCAGT | 2.55 | 3.10 | -0.02 | 0.04 | 2.61 | 3.63 |
| 69 | K-mers | GATGGACCAT | 42.18 | 3.84 | -2.55 | 0.30 | 67.54 | 7.56 |
| 70 | K-mers | CTTACTAGGCCG | 22.51 | 2.56 | -1.02 | 0.13 | 33.95 | 3.47 |
| 71 | K-mers | GAACCTTTCGGG | 14.74 | 2.40 | -0.42 | 0.07 | 17.31 | 2.15 |
| 72 | K-mers | CGCGAGACTG | 8.80 | 1.31 | -0.65 | 0.08 | 17.51 | 1.64 |
| 73 | K-mers | GTGCGTCG | 4.85 | 1.26 | 0.02 | 0.09 | 6.70 | 1.11 |
| 74 | K-mers | TACGGGTAACCTAA | 9.53 | 1.29 | -0.26 | 0.07 | 12.91 | 1.18 |
| 75 | K-mers | CGTGGCAAAAGA | 32.78 | 1.96 | 0.36 | 0.06 | 9.25 | 0.88 |
| 76 | K-mers | ATGTTAAAGCCCAAAGGA | 8.63 | 1.67 | -0.56 | 0.06 | 12.93 | 1.73 |
| 77 | K-mers | CCACCAGACTAGCGTC | 8.26 | 1.70 | -0.06 | 0.09 | 9.70 | 1.26 |
| 78 | K-mers | TCTACTTACATGAAGC | 9.35 | 2.35 | -0.21 | 0.08 | 9.46 | 2.22 |
| 79 | K-mers | GTGCAGGAAGGA | 5.56 | 1.16 | -0.01 | 0.09 | 7.42 | 1.60 |
| 80 | K-mers | CCCATGGCAATTTC | 5.43 | 2.15 | 0.03 | 0.15 | 3.98 | 1.41 |
| 81 | K-mers | CCCAACGACCGGACCCGTCGGT | 5.77 | 1.65 | 0.00 | 0.09 | 5.76 | 1.47 |
| 82 | K-mers | AGTGCCGGTCCTGAGA | 5.56 | 1.43 | -0.04 | 0.07 | 6.33 | 1.17 |
| 83 | K-mers | TGCTCTGATG | 9.96 | 1.93 | -0.32 | 0.14 | 11.74 | 1.66 |
| 84 | K-mers | TATGGCCAAAAA | 16.03 | 2.40 | -0.10 | 0.14 | 23.59 | 2.57 |
| 85 | K-mers | TAGTTAGGGTCTCTGCTTCTACACAGGAAA | 9.97 | 7.17 | 0.00 | 0.00 | 7.95 | 10.00 |
| 86 | K-mers | TAGTGAGGTTCTCTGCTTCTACACAAGCAA | 1.03 | 0.00 | 0.00 | 0.00 | 0.00 | 0.00 |
| 87 | K-mers | TACACACACGACATGTCTTTACGTTATTGC | 0.17 | 3.43 | 0.00 | 0.00 | 0.28 | 2.98 |
| 88 | K-mers | GCCATCTGCAAGTACGTGCCACAACGCGCG | 2.84 | 1.07 | 0.00 | 0.00 | 2.83 | 1.34 |
| 89 | Rational | GTGTGTGTGTGTGTGTGTGTGTGTGTGTGT | 1.22 | 0.26 | 0.00 | 0.02 | 1.19 | 0.18 |
| 90 | Rational | ATATATATATATATATATATATATATATAT | 1.13 | 0.41 | -0.01 | 0.01 | 1.34 | 0.31 |
| 91 | Rational | ACACACACACACACACACACACACACACAC | 1.09 | 0.43 | 0.01 | 0.03 | 0.80 | 0.28 |
| 92 | Rational | ATATATATATATAT | 1.48 | 1.51 | 0.05 | 0.03 | 1.59 | 1.30 |
| 93 | Rational | GTGTGTGTGTGTGT | 4.32 | 1.11 | -0.04 | 0.03 | 5.38 | 0.98 |
| 94 | Rational | GTGTGTGTGT | 4.32 | 1.11 | -0.04 | 0.03 | 5.38 | 0.98 |
| 95 | Rational | ACACACACACACACACACACACAC | 2.11 | 1.59 | 0.12 | 0.11 | 2.05 | 0.65 |
| 96 | Rational | ATATATATATATATATATATATAT | 2.30 | 1.82 | 0.06 | 0.05 | 2.72 | 1.44 |
| 97 | Rational | GTGTGTGTGTGTGTGTGTGTGTGT | 3.69 | 1.41 | 0.05 | 0.06 | 3.62 | 1.12 |
| 98 | Rational | ACACACACACACACACACACACACACAC | 2.46 | 0.52 | 0.10 | 0.09 | 2.49 | 0.12 |
| 99 | Rational | GTGTGTGTGTGTGTGTGTGTGTGTGTGTGT | 2.35 | 1.71 | 0.05 | 0.08 | 2.30 | 1.32 |
| 100 | Rational | ATATATATATATATATATATATATATATAT | 2.17 | 0.89 | 0.07 | 0.08 | 2.38 | 0.56 |
| 101 | Random | ATT CGC CCA GTT AAA AGA CAT TCC TCA TCC | 1.22 | 0.26 | 0.00 | 0.02 | 1.19 | 0.18 |
| 102 | Random | TTA ACC CGG GCT CAT AGG CGA TGC GCC AGC | 1.48 | 2.42 | 0.00 | 0.00 | 0.61 | 2.27 |
| 103 | Random | GTA TCT ACT CCG CCT AGG CCG CCG ACT GGG | 1.09 | 0.43 | 0.01 | 0.03 | 0.80 | 0.28 |
| 104 | Random | GAT CGT CAA AAA CCG CGG TTT GAC GGG TAT | 2.11 | 1.59 | 0.12 | 0.11 | 2.05 | 0.65 |
| 105 | Random | GGC TAC GGA TGA TAC GGT GGA TCC AAG AAG | 3.32 | 1.11 | -0.04 | 0.03 | 5.38 | 0.98 |
| 106 | Random | TCC CTA GAT AGA AGA ATA GGA GTG TGA CTC | 1.13 | 0.41 | -0.01 | 0.01 | 1.34 | 0.31 |
| 107 | Random | CAC TGC CAA ATT TAC TCT TAG TGC GCC TCT | 2.46 | 0.52 | 0.10 | 0.09 | 2.49 | 0.12 |
| 108 | Random | AAC TAG TTC GAG ATG AGG TAC GTA CTT GAA | 5.32 | 1.11 | -0.04 | 0.03 | 5.38 | 0.98 |
| 109 | Random | TAC AGC GAT CAT GCC CGG TCC CGT AGT AGA | 1.22 | 0.26 | 0.00 | 0.02 | 1.19 | 0.18 |
| 110 | Random | AGA CGT ATT AGT CAT GTG CCT CCC GGC TTT | 1.09 | 0.43 | 0.01 | 0.03 | 0.80 | 0.28 |
| 111 | Random | GAC GAT GGA GAT TGG TGT CAC GCA AAT CTA | 1.48 | 2.42 | 0.00 | 0.00 | 0.61 | 2.27 |
| 112 | Random | GGT AAA GTC CAA AGA TGC ATG AGG GAC AAA | 3.13 | 0.41 | -0.01 | 0.01 | 1.34 | 0.31 |

**Table S3. Experimental results of the sensor from diverse sequences selected through DNA mutation and clustering step.** Experimental (9,4) peak shift, (9,4) and (7,6) intensity changes in response to glucose detection at 7.5 mM using ssDNA-SWCNT for different diverse ssDNA sequences, including mutation steps and clustering information, along with the DNA sequence.

| Number | Status | DNA | Len | (7,6) Intensity | (7,6) Intensity_Std | (9,4) Shift | (9,4) Shift_Std | (9,4) Intensity | (9,4) Intensity_Std |
| --- | --- | --- | --- | --- | --- | --- | --- | --- | --- |
| 1 | Parent | GATGGACCAT | 10 | 42.18 | 3.84 | -2.55 | 0.30 | 67.54 | 7.56 |
| 2 | 1 mutation | GATGGTCCAT | 10 | 16.63 | 1.67 | 0.22 | 0.15 | 15.45 | 1.90 |
| 3 | 1 mutation | GATAGACCAT | 10 | 24.33 | 2.12 | -0.24 | 0.21 | 29.45 | 6.29 |
| 4 | 1 mutation | GATGGAGCAT | 10 | 5.44 | 2.04 | -0.05 | 0.24 | 16.03 | 2.15 |
| 5 | 1 mutation | GATGGAACCT | 10 | 19.39 | 2.90 | -0.18 | 0.22 | 26.48 | 1.56 |
| 6 | 2 mutation | GATGCATCAT | 10 | 2.94 | 2.92 | 0.10 | 0.16 | 2.84 | 2.23 |
| 7 | 2 mutation | GATGGAATAT | 10 | 12.80 | 1.91 | 0.17 | 0.16 | 11.04 | 2.65 |
| 8 | 2 mutation | GATGGACAAG | 10 | 13.57 | 2.17 | -0.02 | 0.17 | 16.79 | 2.21 |
| 9 | 2 mutation | GACGGACTAT | 10 | 36.68 | 3.55 | -1.12 | 0.31 | 62.18 | 10.42 |
| 10 | 2 mutation | GACGGACAAT | 10 | 40.16 | 9.61 | -0.25 | 0.21 | 50.58 | 10.89 |
| 11 | 2 mutation | GTTGGACCTT | 10 | 40.11 | 6.85 | -1.17 | 0.34 | 61.73 | 18.18 |
| 12 | 2 mutation | AATGGACCGC | 10 | 7.10 | 2.09 | -0.55 | 0.27 | 17.85 | 0.61 |
| 13 | 3 mutation | GATGAACTAA | 10 | 12.52 | 3.22 | -0.53 | 0.28 | 28.94 | 1.82 |
| 14 | 3 mutation | GACGGACAGT | 10 | 16.04 | 1.52 | -0.20 | 0.24 | 21.86 | 4.72 |
| 15 | 3 mutation | GACAGACCAA | 10 | 20.81 | 4.31 | -0.16 | 0.13 | 29.29 | 4.09 |
| 16 | 3 mutation | GGTGAATCAT | 10 | 5.37 | 3.12 | 0.10 | 0.21 | 6.49 | 2.96 |
| 17 | 3 mutation | GCTTGACCTT | 10 | 26.65 | 4.87 | -0.87 | 0.26 | 42.53 | 9.63 |
| 18 | 3 mutation | AGAGGACCAT | 10 | 23.75 | 3.28 | -0.89 | 0.29 | 48.61 | 9.59 |
| 19 | 3 mutation | GACAGACCAC | 10 | 7.51 | 4.85 | 0.32 | 0.23 | 5.13 | 4.35 |
| 20 | 3 mutation | GGTGGGTCAT | 10 | 45.39 | 14.09 | -0.15 | 0.21 | 50.21 | 16.66 |
| 21 | 3 mutation | GGTGGGCCTT | 10 | 10.67 | 3.72 | -0.09 | 0.19 | 15.56 | 3.44 |
| 22 | 3 mutation | GTGGGACCTT | 10 | 21.08 | 5.77 | -0.32 | 0.32 | 32.38 | 9.01 |
| 23 | 3 mutation | GCTGTACCCT | 10 | 15.41 | 4.47 | -0.02 | 0.22 | 17.77 | 6.23 |
| 24 | 3 mutation | GATGGGCCGC | 10 | 6.76 | 0.75 | -0.06 | 0.17 | 11.09 | 1.86 |
| 25 | 3 mutation | AGCGGACCAT | 10 | 16.34 | 1.84 | -1.04 | 0.43 | 25.60 | 4.27 |
| 26 | 3 mutation | GACGGTCCAA | 10 | 6.66 | 0.50 | -0.14 | 0.19 | 13.07 | 2.98 |
| 27 | 3 mutation | AATGGACTCT | 10 | 6.17 | 2.94 | 0.01 | 0.14 | 10.76 | 1.86 |
| 28 | 3 mutation | GGTTGACCCT | 10 | 8.91 | 2.32 | 0.02 | 0.16 | 10.43 | 3.42 |
| 29 | 3 mutation | GATGGACAGC | 10 | 6.12 | 2.96 | 0.04 | 0.15 | 7.60 | 2.25 |
| 30 | 3 mutation | GAGTGACCAA | 10 | 24.69 | 2.57 | -0.64 | 0.19 | 46.07 | 5.46 |
| 31 | Length mutations | GGATGGACCATC | 12 | 4.42 | 1.39 | 0.05 | 0.12 | 4.27 | 0.90 |
| 32 | Length mutations | GCAGATGGACCAT | 13 | 52.66 | 7.33 | -1.91 | 0.41 | 40.04 | 9.10 |
| 33 | Length mutations | CTTGATGGACCAT | 13 | 49.70 | 9.14 | 0.15 | 0.13 | 51.51 | 14.02 |
| 34 | Length mutations | TTGATGGACCATCC | 14 | 4.15 | 1.91 | 0.06 | 0.12 | 3.34 | 2.60 |
| 35 | Length mutations | ATAGATGGACCATTCG | 16 | 8.45 | 5.03 | 0.14 | 0.19 | 7.79 | 4.89 |
| 36 | Length mutations | GATGATGGACCATACT | 16 | 5.61 | 0.75 | 0.10 | 0.14 | 3.75 | 0.33 |
| 37 | Length mutations | AGAGATGGACCATTTC | 16 | 6.17 | 0.84 | -0.07 | 0.13 | 4.78 | 0.67 |
| 38 | Length mutations | CTCGATGGACCATGTG | 16 | 5.27 | 1.83 | 0.02 | 0.19 | 8.14 | 2.50 |
| 39 | Length mutations | CCTGATGGACCATTAC | 16 | 7.21 | 2.33 | -0.05 | 0.19 | 6.51 | 2.53 |
| 40 | Length mutations | CAAGATGGACCATTTA | 16 | 14.46 | 1.34 | 0.09 | 0.17 | 12.09 | 0.98 |
| 41 | Length mutations | GCGGATGGACCATGCG | 16 | 6.25 | 0.70 | 0.06 | 0.16 | 6.61 | 2.28 |
| 42 | Length mutations | AGGGATGGACCATAAT | 16 | 67.31 | 10.76 | -4.83 | 0.88 | 62.14 | 18.02 |
| 43 | Length mutations | GAGGATGGACCATTTA | 16 | 16.37 | 1.82 | -1.68 | 0.51 | 31.21 | 8.88 |
| 44 | Length mutations | CACGATGGACCATACG | 16 | 4.89 | 1.39 | 0.08 | 0.15 | 6.56 | 1.78 |
| 45 | Length mutations | GGAGATGGACCATCTC | 16 | 1.95 | 4.16 | 0.09 | 0.11 | 2.80 | 4.13 |
| 46 | Length mutations | TAAGATGGACCATTTT | 16 | 39.50 | 11.28 | -1.15 | 0.37 | 60.67 | 15.82 |
| 47 | Rational design | GATGGACCATGATGGACCAT | 20 | 33.01 | 4.59 | -0.24 | 0.10 | 35.68 | 6.18 |
| 48 | Rational design | GATGGACCATCGCGAGACTG | 20 | 7.06 | 2.07 | -0.13 | 0.25 | 10.49 | 2.38 |
| 49 | Rational design | GATGGACCATCTTACTAGGCCG | 22 | 6.73 | 0.71 | -0.05 | 0.21 | 9.97 | 2.05 |
| 50 | Rational design | CTTACTAGGCCGGATGGACCAT | 22 | 6.16 | 4.03 | -0.11 | 0.24 | 10.82 | 3.99 |
| 51 | Rational design | GATGGACCATGAACCTTTCGGG | 22 | 13.52 | 7.19 | -0.29 | 0.36 | 20.77 | 8.43 |
| 52 | Rational design | CTTACTAGGCCGCTTACTAGGCCG | 24 | 7.31 | 2.99 | -0.08 | 0.27 | 9.35 | 3.83 |
| 53 | Rational design | GATGGACCATATGTTAAAGCCCAAAGGA | 28 | 10.07 | 2.69 | 0.18 | 0.73 | 9.95 | 6.81 |
| 54 | Rational design | GATGGACCATGATGGACCATGATGGACCAT | 30 | 11.93 | 1.03 | 0.06 | 0.19 | 12.78 | 2.30 |

**Table S4: Hyperparameter optimization for Deep neural network training.** Hyperparameter optimization for the CNN model using Optuna search, including k-fold cross-validation. The computational analysis was executed using a custom script (nano run.sh) on the IZAR EPFL cluster. The script was configured to utilize the GPU partition, requesting one GPU, one compute node, 40 CPUs per task, and 120 GB of memory. The results for CNN optimization have been reported only for the CNN model to avoid overloading information in the paper. More detailed optimization processes for both GNN and CNN, along with the Python scripts, are available on GitHub.

| **Parameter** | **Type/Options** | **Description** | **Result** |
| --- | --- | --- | --- |
| num_filters1 | Categorical: [4,8,16, 32,64,128] | Number of filters in the first Conv2D layer. | 8 |
| num_filters2 | Categorical: [4,8,16, 32,64] | Number of filters in the second Conv2D layer. | 8 |
| kernel_size | Categorical: [(2, 2),(3, 3), (4, 4) ,(5, 5)] | Size of the convolution kernel. | (2,2) |
| dense_units | Categorical: [8,16,20,32,64,128,200] | Number of neurons in the dense layer. | 64 |
| dropout_rate | Uniform: 0.0 to 0.7 | Dropout rate for regularization. | 0.196429 |
| learning_rate | Loguniform: 1e-6 to 1e-1 | Learning rate for the optimizer. | 0.009916 |
| pooling_type | Categorical: ['max', 'average'] | Type of pooling layer. | Average |
| pool_size1 | Categorical: [(2, 1),(2, 2) ,(3, 1),(3, 2) ,(3, 3),(4, 1),(4, 2)] | Pool size for the first pooling layer. | (4,2) |
| pool_size2 | Categorical: [(2, 1),(2, 2) ,(3, 1),(3, 2) ,(3, 3),(4, 1),(4, 2)] | Pool size for the second pooling layer. | (4,2) |
| optimizer | Categorical: ['adam', 'sgd', 'rmsprop'] | Type of optimizer to use. | Adam |
| momentum (if SGD) | Uniform: 0.0 to 1.0 | Momentum for the SGD optimizer. | - |
| num_folds | Integer: 2 to 30 | Number of folds in K-fold cross-validation. | 10 |
| n_trials | Integer: 10000 | Number of trials for the Optuna study to optimize hyperparameters. | 10000 |

**Table S5.** **The model performance for deep neural network training**. The model performance achieved high accuracy for the test data with intensity responses (7,6) and (9,4), resulting in accuracies of 0.81 and 0.90, respectively.

|  | I(7,6) | I(9,4) |
| --- | --- | --- |
| Accuracy | **0.81** | **0.90** |
| Precision | **0.7** | **0.83** |
| Recall | **0.7** | **0.71** |
| F1 Score | **0.7** | **0.77** |
| AUC Score | **0.90** | **0.87** |

**Table S6. Pattern recognition for identifying different effective K-mers in relation to sensor response.** This table shows the K-mers with high frequency and high-intensity change at both chiralities (9,4) and (7,6), which led to the development of an ssDNA-SWCNT sensor responsive to detecting glucose at 7.5 mM.

| **Common K-mers** | **Frequency of K-mers** | **Mean (7,6)**  **Intensity change (%)** | **Mean (9,4)**  **Intensity change (%)** |
| --- | --- | --- | --- |
| GGA | 81 | 12.96 | 16.45 |
| GAC | 78 | 14.24 | 18.72 |
| TGG | 71 | 13.18 | 15.60 |
| ATG | 65 | 12.37 | 14.92 |
| ACC | 63 | 14.76 | 18.55 |
| CCA | 57 | 13.63 | 16.77 |
| GAT | 55 | 12.76 | 15.41 |
| CAT | 47 | 13.95 | 16.78 |

**Table S7. Discovery of the Responsive DNA Sequence for Glucose Detection.** This table shows the intensity changes at chiralities (7,6) and (9,4), and the peak shift at chirality (9,4) during different steps of our study. This study led to the identification of a responsive sequence that can detect glucose as a small molecule by changing the intensity or peak shift of our ssDNA-SWCNT sensor.

| **DNA** | **Status** | **len** | **(7,6) Intensity** | **(7,6) Intensity_Std** | **(9,4) Shift** | **(9,4) Shift_Std** | **(9,4) Intensity** | **(9,4) Intensity_Std** |
| --- | --- | --- | --- | --- | --- | --- | --- | --- |
| GATGGACCAT | Clustering | 10 | 42.17711799 | 3.838207438 | -2.551782585 | 0.295600044 | 67.54265896 | 7.562649912 |
| GACGCCCGCC | Clustering | 10 | 29.85177966 | 4.046013832 | -0.586047945 | 0.346767495 | 46.67095977 | 11.44792463 |
| CTTACTAGGCCG | Clustering | 12 | 22.50727018 | 2.557852783 | -1.016552069 | 0.125156583 | 33.95120501 | 3.466082469 |
| GACGGACTAT | Mutation & Clustering | 10 | 36.67839285 | 3.548616516 | -1.12347993 | 0.308592415 | 62.18338222 | 10.42167164 |
| AGGGATGGACCATAAT | Mutation & Clustering | 16 | 67.30595859 | 10.75788758 | -4.826981098 | 0.883933179 | 62.14140606 | 18.02280915 |
| GTTGGACCTT | Mutation & Clustering | 10 | 40.10750725 | 6.847686966 | -1.166701879 | 0.340117507 | 61.7274637 | 18.17893111 |
| TAAGATGGACCATTTT | Mutation & Clustering | 16 | 39.49694354 | 11.283478 | -1.152219496 | 0.374351886 | 60.66575548 | 15.81929997 |
| CTTGATGGACCAT | Mutation & Clustering | 13 | 49.69648004 | 9.137705404 | 0.151397038 | 0.128040122 | 51.51451095 | 14.01568782 |
| GACGGACAAT | Mutation & Clustering | 10 | 40.164008 | 9.607672356 | -0.254074099 | 0.205269416 | 50.57845002 | 10.88541709 |
| GGTGGGTCAT | Mutation & Clustering | 10 | 45.39307249 | 14.08878974 | -0.149443634 | 0.205141492 | 50.21413019 | 16.66008948 |
| AGAGGACCAT | Mutation & Clustering | 10 | 23.74972074 | 3.282494287 | -0.886961583 | 0.289226556 | 48.60582623 | 9.587056081 |
| GAGTGACCAA | Mutation & Clustering | 10 | 24.69344596 | 2.573827713 | -0.635623353 | 0.194422106 | 46.06875445 | 5.456515388 |
| GCTTGACCTT | Mutation & Clustering | 10 | 26.64884465 | 4.872511845 | -0.867169813 | 0.25848433 | 42.52701656 | 9.631318627 |
| GCAGATGGACCAT | Mutation & Clustering | 13 | 52.65826142 | 7.330842844 | -1.909669372 | 0.414024089 | 40.03982053 | 9.098686858 |
| GATGGACCATGATGGACCAT | Mutation & Clustering | 20 | 33.01199794 | 4.586525931 | -0.23642341 | 0.097681167 | 35.67836662 | 6.184584548 |
| GTGGGACCTT | Mutation & Clustering | 10 | 21.07680944 | 5.772903851 | -0.321447737 | 0.318246073 | 32.38437666 | 9.013237398 |
| GAGGATGGACCATTTA | Mutation & Clustering | 16 | 16.36617943 | 1.818185542 | -1.6784525 | 0.51251613 | 31.20509454 | 8.883571275 |
| GATGAACTAA | Mutation & Clustering | 10 | 12.52235718 | 3.219095427 | -0.52638995 | 0.281224601 | 28.9383315 | 1.820325359 |
| AGCGGACCAT | Mutation & Clustering | 10 | 16.3363101 | 1.840483113 | -1.040460523 | 0.430665134 | 25.60129064 | 4.272906866 |
| GATGGGCCAT | ML Prediction | 10 | 30.3971381 | 4.145318752 | 0.12160947 | 0.058558134 | 37.71593095 | 4.962273802 |
| GAAGGACCAT | ML Prediction | 10 | 41.0120955 | 2.580801518 | -1.49167755 | 0.044412777 | 76.96237371 | 2.952872815 |
| GAGGGACCAT | ML Prediction | 10 | 29.5830206 | 2.435428865 | -1.218742392 | 0.140773762 | 53.70538471 | 2.895051922 |
| GACGGGCCAT | ML Prediction | 10 | 26.40956484 | 5.704855812 | -0.017022793 | 0.145912152 | 30.42459387 | 5.747958051 |
| GGCGGACCAT | ML Prediction | 10 | 29.90246406 | 3.531602127 | -0.370209359 | 0.15389066 | 34.59435243 | 4.797768888 |
| AAAGATGGACCAT | ML Prediction | 13 | 42.76057381 | 2.952694134 | -3.265990252 | 0.409550937 | 55.87643496 | 4.164318568 |
| CAAGATGGACCATTTT | ML Prediction | 16 | 44.61286531 | 3.836329691 | -0.194049819 | 0.11023199 | 48.89730486 | 3.144934761 |
| AAGGATGGACCAT | ML Prediction | 13 | 24.41164352 | 12.84418849 | -0.676504927 | 0.405133438 | 40.82350948 | 16.15800996 |

**Table S8. Biochemical analysis of different serum samples** for sequence responsive 1, sequence responsive 2, and GT15. After 24 hours post-injection of nanoparticles, blood serum was collected from 3 animals per group and sent for serum biochemistry analysis. The results did not show any increase in toxicity markers according to the biochemical test (Group 1 – mice 1-3; Group 2 – mice 4-6; Group 4 – mice 7-9 in the serum analysis report). The results for animal ID 1 for sequence responsive 1, animal ID 4 for sequence responsive 2, and animal ID 9 for the GT sequence are presented as examples.

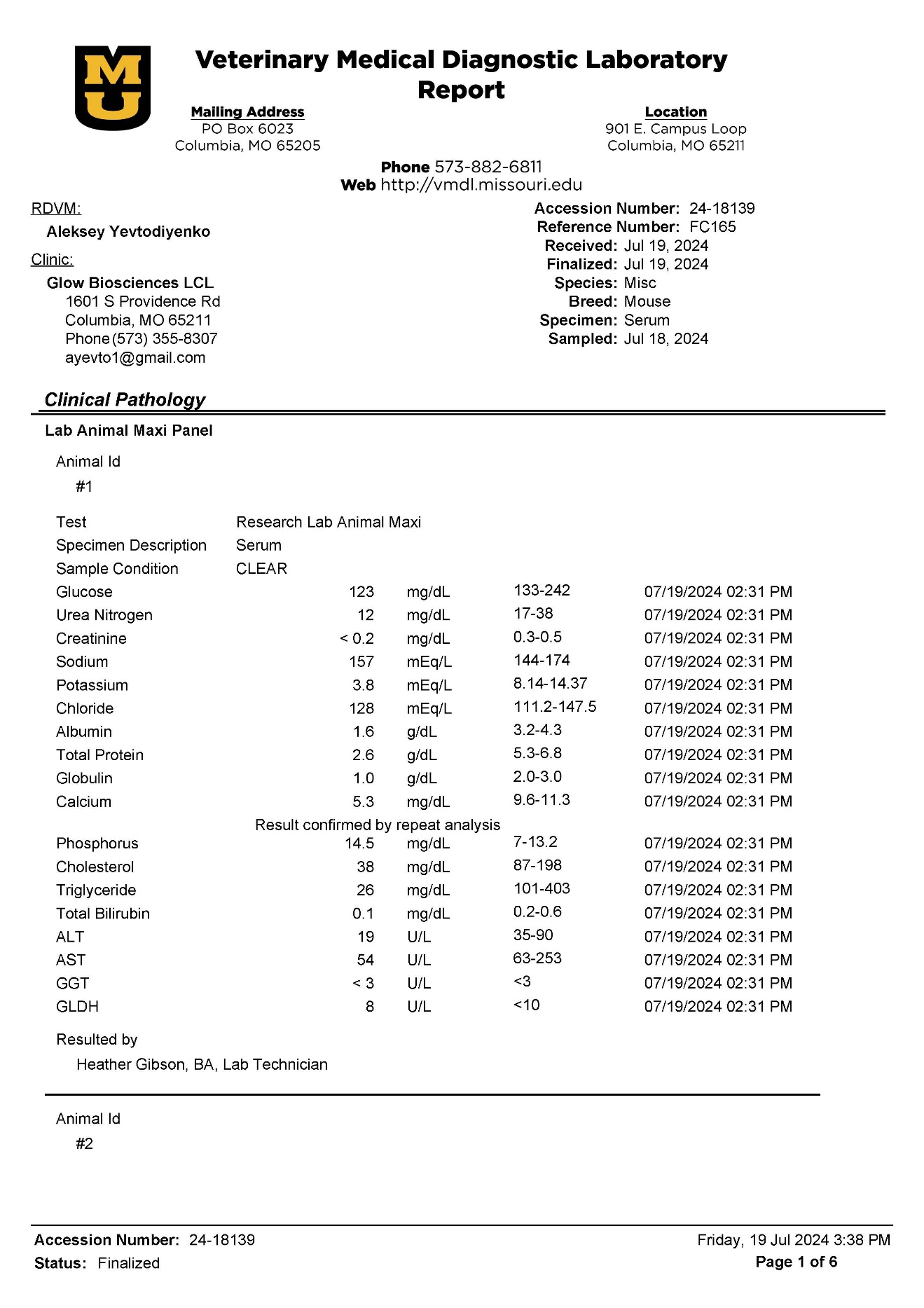

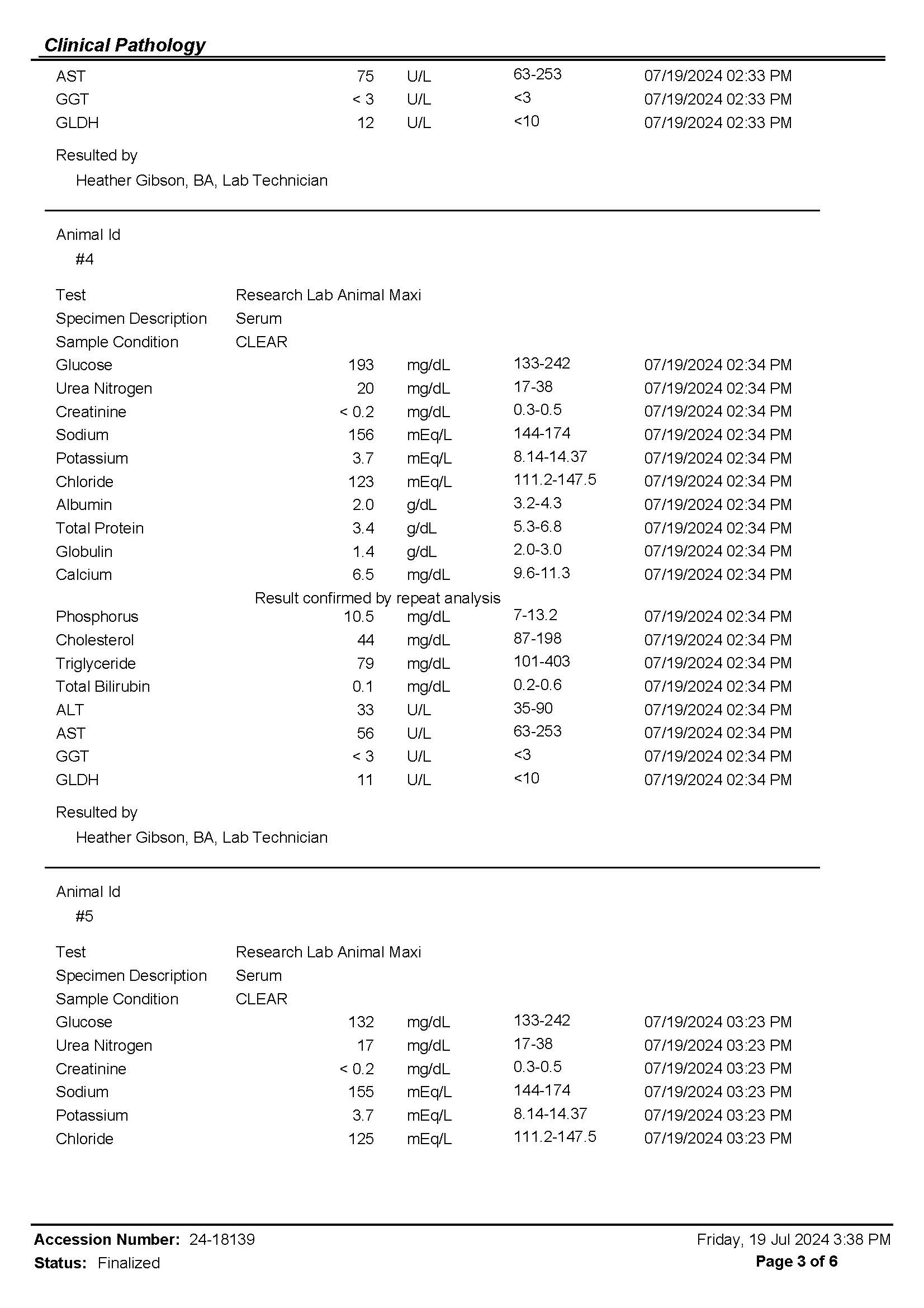

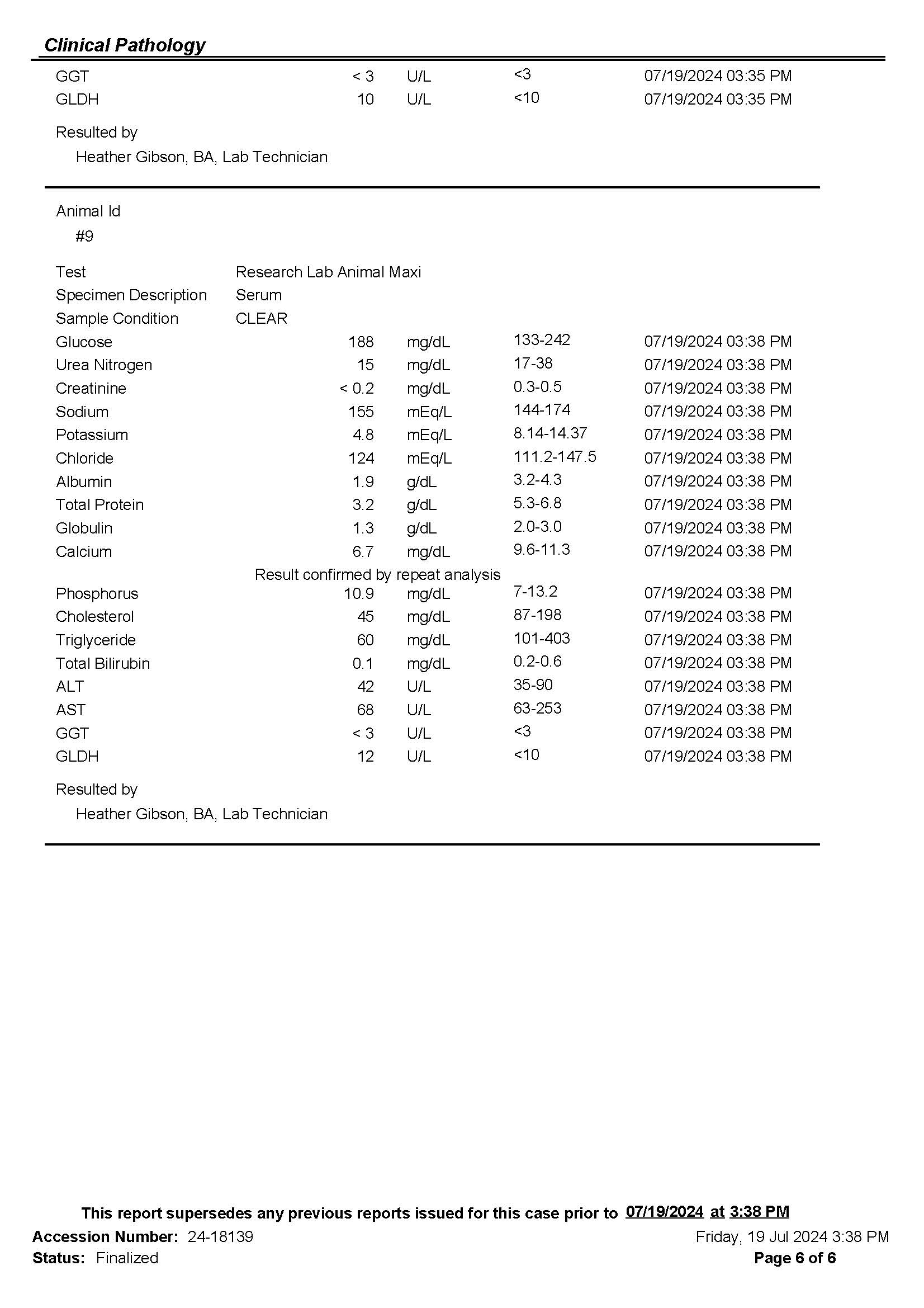

3. Di Filippo, D., Sunstrum, F. N., Khan, J. U. & Welsh, A. W. Non-Invasive Glucose Sensing Technologies and Products: A Comprehensive Review for Researchers and Clinicians. *Sensors*

*(Basel, Switzerland)* 23, 9130. ISSN: 1424-8220. [https://www.ncbi.nlm.nih.gov/pmc/ articles/PMC10674292/](https://www.ncbi.nlm.nih.gov/pmc/articles/PMC10674292/).

4. Tomic, D., Shaw, J. E. & Magliano, D. J. The burden and risks of emerging complications of diabetes mellitus. en. *Nature Reviews Endocrinology* 18. Publisher: Nature Publishing Group,

525–539. ISSN: 1759-5037. <https://www.nature.com/articles/s41574-022-00690-7> .

5. Yoo, E.-H. & Lee, S.-Y. Glucose Biosensors: An Overview of Use in Clinical Practice. *Sensors (Basel, Switzerland)* 10, 4558–4576. ISSN: 1424-8220. [https://www.ncbi.nlm.nih.gov/ pmc/articles/PMC3292132/](https://www.ncbi.nlm.nih.gov/pmc/articles/PMC3292132/).
6. Liu, Y. *et al.* Review of point-of-care platforms for diabetes: (1) sensing. *Sensors and Actuators Reports* 4, 100113. ISSN: 2666-0539. [https://www.sciencedirect.com/science/ article/pii/S2666053922000406](https://www.sciencedirect.com/science/article/pii/S2666053922000406).
7. Mathew, T. K., Zubair, M. & Tadi, P. eng. in *StatPearls* (StatPearls Publishing, Treasure Island (FL), 2024). <http://www.ncbi.nlm.nih.gov/books/NBK555976/>.
8. Yu, Z., Jiang, N., Kazarian, S. G., Tasoglu, S. & Yetisen, A. K. Optical sensors for continuous glucose monitoring. en. *Progress in Biomedical Engineering* 3, 022004. ISSN: 2516-1091. <https://iopscience.iop.org/article/10.1088/2516-1091/abe6f8>.
9. Ahmed, I. *et al.* Recent advances in optical sensors for continuous glucose monitoring. en. *Sensors & Diagnostics* 1. Publisher: Royal Society of Chemistry, 1098–1125. [https:// pubs.rsc.org/en/content/articlelanding/2022/sd/d1sd00030f](https://pubs.rsc.org/en/content/articlelanding/2022/sd/d1sd00030f) .
10. Jena, P. V., Cupo, C. & Heller, D. A. en. in *Near Infrared-Emitting Nanoparticles for Biomedical Applications* (eds Benayas, A., Hemmer, E., Hong, G. & Jaque, D.) 103–132 (Springer International Publishing, Cham, 2020). ISBN: 978-3-030-32036-2. [https://doi.org/10.](https://doi.org/10.1007/978-3-030-32036-2_6)

[1007/978-3-030-32036-2_6](https://doi.org/10.1007/978-3-030-32036-2_6).

11. Yoon, M. *et al.* A nIR fluorescent single walled carbon nanotube sensor for broad-spectrum diagnostics. en. *Sensors & Diagnostics* 3. Publisher: RSC, 203–217. ISSN: 2635-0998. [https:](https://pubs.rsc.org/en/content/articlelanding/2024/sd/d3sd00257h)

[//pubs.rsc.org/en/content/articlelanding/2024/sd/d3sd00257h](https://pubs.rsc.org/en/content/articlelanding/2024/sd/d3sd00257h).

12. Barone, P. W. & Strano, M. S. Reversible Control of Carbon Nanotube Aggregation for a Glucose Affinity Sensor. *Angewandte Chemie International Edition* 45, 8138–8141. ISSN:

1521-3773. <https://onlinelibrary.wiley.com/doi/abs/10.1002/anie.200603138> .

13. Lambert, B. P. J. G. *Directed evolution of DNA-wrapped single-walled carbon nanotube complexes for optical sensing* eng. PhD thesis (EPFL, Lausanne, 2021). 10.5075/epfl-thesis8406.
14. Kupis-Rozmysłowicz, J., Antonucci, A. & Boghossian, A. A. Review—Engineering the Selectivity of the DNA-SWCNT Sensor. en. *ECS Journal of Solid State Science and Technology* 5. Publisher: IOP Publishing, M3067. ISSN: 2162-8777. [https://iopscience.iop.org/ article/10.1149/2.0111608jss/meta](https://iopscience.iop.org/article/10.1149/2.0111608jss/meta).
15. Zhou, Y., Fang, Y. & Ramasamy, R. P. Non-Covalent Functionalization of Carbon Nanotubes for Electrochemical Biosensor Development. *Sensors (Basel, Switzerland)* 19, 392. ISSN:

1424-8220. <https://www.ncbi.nlm.nih.gov/pmc/articles/PMC6358788/>.

16. Yoon, H. *et al.* Periplasmic Binding Proteins as Optical Modulators of Single-Walled Carbon Nanotube Fluorescence: Amplifying a Nanoscale Actuator. *Angewandte Chemie International Edition* 50. _eprint: https://onlinelibrary.wiley.com/doi/pdf/10.1002/anie.201006167,

1828–1831. ISSN: 1521-3773. [https://onlinelibrary.wiley.com/doi/abs/10.1002/ anie.201006167](https://onlinelibrary.wiley.com/doi/abs/10.1002/anie.201006167).

17. Barone, P. W., Baik, S., Heller, D. A. & Strano, M. S. Near-infrared optical sensors based on single-walled carbon nanotubes. en. *Nature Materials* 4. Publisher: Nature Publishing Group,

86–92. ISSN: 1476-4660. <https://www.nature.com/articles/nmat1276> .

18. Karachevtsev, V. A., Glamazda, A. Y., Leontiev, V. S., Lytvyn, O. S. & Dettlaff-Weglikowska, U. Glucose sensing based on NIR fluorescence of DNA-wrapped single-walled carbon nanotubes. *Chemical Physics Letters* 435, 104–108. ISSN: 0009-2614. [https://www.sciencedir](https://www.sciencedirect.com/science/article/pii/S0009261406018616)ect. [com/science/article/pii/S0009261406018616](https://www.sciencedirect.com/science/article/pii/S0009261406018616).
19. Juska, Vuslat B., and Martyn E. Pemble.. "A Critical Review of Electrochemical Glucose Sensing: Evolution of Biosensor Platforms Based on Advanced Nanosystems" Sensors 20, no. 21: 6013.2020. <https://doi.org/10.3390/s20216013>
20. Zubkovs, V., Schuergers, N., Lambert, B., Ahunbay, E. & Boghossian, A. A. Mediatorless, Reversible Optical Nanosensor Enabled through Enzymatic Pocket Doping. en. *Small* 13.

_eprint: https://onlinelibrary.wiley.com/doi/pdf/10.1002/smll.201701654, 1701654. ISSN: 1613-

6829. <https://onlinelibrary.wiley.com/doi/abs/10.1002/smll.201701654> .

21. Gillen, A. J. & Boghossian, A. A. Non-covalent Methods of Engineering Optical Sensors Based on Single-Walled Carbon Nanotubes. English. *Frontiers in Chemistry* 7. Publisher:

Frontiers. ISSN: 2296-2646. [https://www.frontiersin.org/articles/10.3389/ fchem.2019.00612](https://www.frontiersin.org/articles/10.3389/fchem.2019.00612).

22. Beyene, A. G. et al. Ultralarge Modulation of Single Wall Carbon Nanotube Fluorescence Mediated by Neuromodulators Adsorbed on Arrays of Oligonucleotide Rings en. June 2018. <http://biorxiv.org/lookup/doi/10.1101/351627>.
23. Zhang, J., Landry, M., Barone, P. et al. Molecular recognition using corona phase complexes made of synthetic polymers adsorbed on carbon nanotubes. Nature Nanotech 8, 959–968 (2013). <https://doi.org/10.1038/nnano.2013.236>
24. Lambert, B., Gillen, A. J., Schuergers, N., Wu, S.-J. & Boghossian, A. A. Directed evolution of the optoelectronic properties of synthetic nanomaterials. en. *Chemical Communications* 55. Publisher: The Royal Society of Chemistry, 3239–3242. ISSN: 1364-548X. [https://pubs.](https://pubs.rsc.org/en/content/articlelanding/2019/cc/c8cc08670b)

[rsc.org/en/content/articlelanding/2019/cc/c8cc08670b](https://pubs.rsc.org/en/content/articlelanding/2019/cc/c8cc08670b).

25. Lambert, B. P. *et al. Directed evolution of nanosensors for the detection of mycotoxins* en. Pages: 2023.06.13.544576 Section: New Results. June 2023. [https://www.biorxiv.org/ content/10.1101/2023.06.13.544576v1](https://www.biorxiv.org/content/10.1101/2023.06.13.544576v1).
26. Gong, X., Renegar, N., Levi, R. & Strano, M. S. Machine learning for the discovery of molecular recognition based on single-walled carbon nanotube corona-phases. en. *npj Computational Materials* 8. Publisher: Nature Publishing Group, 1–13. ISSN: 2057-3960. [https://www. nature.com/articles/s41524-022-00795-7](https://www.nature.com/articles/s41524-022-00795-7).
27. Heller, D. A. *et al.* Multimodal optical sensing and analyte specificity using single-walled carbon nanotubes. en. *Nature Nanotechnology* 4. Publisher: Nature Publishing Group, 114–

120. ISSN: 1748-3395. <https://www.nature.com/articles/nnano.2008.369>.

28. Lambert, B. P., Gillen, A. J. & Boghossian, A. A. Synthetic Biology: A Solution for Tackling Nanomaterial Challenges. *The Journal of Physical Chemistry Letters* 11. Publisher: American

Chemical Society, 4791–4802. <https://doi.org/10.1021/acs.jpclett.0c00929> .

29. Alizadehmojarad, A. A. *et al.* Binding Affinity and Conformational Preferences Influence Kinetic Stability of Short Oligonucleotides on Carbon Nanotubes. en. *Advanced Materials Interfaces* 7. _eprint: https://onlinelibrary.wiley.com/doi/pdf/10.1002/admi.202000353.
30. Mann, F. A., Herrmann, N., Meyer, D. & Kruss, S. Tuning Selectivity of Fluorescent Carbon Nanotube-Based Neurotransmitter Sensors. *Sensors (Basel, Switzerland)* 17, 1521. ISSN: 1424-8220. <https://www.ncbi.nlm.nih.gov/pmc/articles/PMC5539566/>
31. Salila Vijayalal Mohan, H. K., An, J. & Zheng, L. Sequence-dependent electrical response of ssDNA-decorated carbon nanotube, field-effect transistors to dopamine. *Beilstein Journal of*

*Nanotechnology* 5, 2113–2121. ISSN: 2190-4286. [https://www.ncbi.nlm.nih.gov/pmc/ articles/PMC4273222/](https://www.ncbi.nlm.nih.gov/pmc/articles/PMC4273222/)

32. Jena, P. V., Safaee, M. M., Heller, D. A. & Roxbury, D. DNA-Carbon Nanotube Complexation Affinity and Photoluminescence Modulation Are Independent. *ACS Applied Materials & Interfaces* 9. Publisher: American Chemical Society, 21397–21405. ISSN: 1944-8244. [https: //doi.org/10.1021/acsami.7b05678](https://doi.org/10.1021/acsami.7b05678)
33. Beyene, A. G. *et al. Ultralarge Modulation of Single Wall Carbon Nanotube Fluorescence Mediated by Neuromodulators Adsorbed on Arrays of Oligonucleotide Rings* en. June 2018. <http://biorxiv.org/lookup/doi/10.1101/351627>
34. Zhang, J. *et al.* Single Molecule Detection of Nitric Oxide Enabled by d(AT)15 DNA Adsorbed to Near Infrared Fluorescent Single-Walled Carbon Nanotubes. *Journal of the American Chemical Society* 133. Publisher: American Chemical Society, 567–581. ISSN: 00027863. <https://doi.org/10.1021/ja1084942>.
35. Ronny Lorenz, Stephan H. Bernhart, Christian Höner zu Siederdissen, Hakim Tafer, Christoph Flamm, Peter F. Stadler, and Ivo L. Hofacker. ViennaRNA package 2.0. Algorithms for Molecular Biology, 6(1):26, 2011. https://doi.org/[10.1186/1748-7188-6-26](https://doi.org/10.1186/1748-7188-6-26)
36. I.L. Hofacker, W. Fontana, P.F. Stadler, L.S. Bonhoeffer, M. Tacker, and P. Schuster. Fast folding and comparison of RNA secondary structures. Monatshefte für Chemie/Chemical Monthly, 125(2):167–188, 1994. https://link.springer.com/article/10.1007/BF00818163
37. S. Kruss, M. P. Landry, E. V. Ende, B.M.A. Lima, N.F. Reuel, J. Zhang, J.Nelson, B.Mu, A.Hilmer, and M. Strano, Neurotransmitter Detection Using Corona Phase Molecular Recognition on Fluorescent Single-Walled Carbon Nanotube Sensors, Journal of the American Chemical Society 2014 136 (2), 713-724, <https://doi.org/10.1021/ja410433b> .
38. E. Polo and S. Kruss, Impact of Redox-Active Molecules on the Fluorescence of Polymer-Wrapped Carbon Nanotubes, *The Journal of Physical Chemistry C* **2016** *120* (5), 3061-3070. https://doi.org/10.1021/acs.jpcc.5b12183
39. Daniel A. Heller et al. Optical Detection of DNA Conformational Polymorphism on Single-Walled Carbon Nanotubes.Science311,508-511(2006). https://doi.org/[10.1126/science.1120792](https://doi.org/10.1126/science.1120792)
40. Hong Jin, Esther S. Jeng, Daniel A. Heller, Prakrit V. Jena, Robert Kirmse, Jörg Langowski, and Michael S. Strano, Divalent Ion and Thermally Induced DNA Conformational Polymorphism on Single-walled Carbon Nanotubes, *Macromolecules* **2007** *40* (18), 6731-6739. <https://doi.org/10.1021/ma070608t>
41. Albertorio F, Hughes ME, Golovchenko JA, Branton D. Base dependent DNA-carbon nanotube interactions: activation enthalpies and assembly-disassembly control. Nanotechnology. 2009 Sep 30;20(39):395101. <https://doi.org/10.1088/0957-4484/20/39/395101>
42. [Esther S. Jeng](https://onlinelibrary.wiley.com/authored-by/Jeng/Esther%E2%80%85S.), [Paul W. Barone](https://onlinelibrary.wiley.com/authored-by/Barone/Paul%E2%80%85W.), [John D. Nelson](https://onlinelibrary.wiley.com/authored-by/Nelson/John%E2%80%85D.), [Michael S. Strano Prof](https://onlinelibrary.wiley.com/authored-by/Strano/Michael%E2%80%85S.), [Hybridization Kinetics and Thermodynamics of DNA Adsorbed to Individually Dispersed Single‐Walled Carbon Nanotubes - Jeng - 2007 - Small - Wiley Online Library](https://onlinelibrary.wiley.com/doi/full/10.1002/smll.200700141?msockid=3fc8229cb9316df832333611b83a6c32) , <https://doi.org/10.1002/smll.200700141>
43. Jaroslav Kypr, Iva Kejnovská, Daniel Renčiuk, Michaela Vorlíčková, Circular dichroism and conformational polymorphism of DNA, *Nucleic Acids Research*, Volume 37, Issue 6, 1 April 2009, Pages 1713–1725, <https://doi.org/10.1093/nar/gkp026>
44. G. Dukovic, Milan Balaz, Peter Doak, Nina D. Berova, Ming Zheng, Robert S. Mclean, and Louis E. Brus, Racemic Single-Walled Carbon Nanotubes Exhibit Circular Dichroism When Wrapped with DNA, Journal of the American Chemical Society 2006 128 (28), 9004-9005 <https://doi.org/10.1021/ja062095w>
45. Kypr J, Kejnovská I, Renciuk D, Vorlícková M. Circular dichroism and conformational polymorphism of DNA. Nucleic Acids Res. 2009 Apr;37(6):1713-25. https://doi.org/ 10.1093/nar/gkp026.
46. Adrien Marchand, Valérie Gabelica, Folding and misfolding pathways of G-quadruplex DNA, *Nucleic Acids Research*, Volume 44, Issue 22, December 2016, Pages 10999–11012, <https://doi.org/10.1093/nar/gkw970>.
47. Jaroslav Kypr, Iva Kejnovská, Daniel Renčiuk, Michaela Vorlíčková, Circular dichroism and conformational polymorphism of DNA, *Nucleic Acids Research*, Volume 37, Issue 6, 1 April 2009, Pages 1713–1725, <https://doi.org/10.1093/nar/gkp026>.
